# A spatially resolved EGFR signaling model predicts the length scale of GAB1-SHP2 complex persistence

**DOI:** 10.1101/2021.11.08.467801

**Authors:** Paul J. Myers, Christopher M. Furcht, Itziar Pinilla-Macua, Alexander Sorkin, William M. Deen, Matthew J. Lazzara

## Abstract

Activation of receptor tyrosine kinases (RTKs) promotes the intracellular assembly of protein complexes formed through phosphotyrosine-SH2 domain linkages. These interactions are weak and reversible, and the dephosphorylation of uncomplexed phosphorylated proteins can be rapid enough to reduce the time and length scales of protein complex persistence distal from the receptor. This limitation may be overcome by cytosolic kinases that rephosphorylate constituent complex proteins as they diffuse. To develop quantitative understanding of the spatiotemporal regulation of these processes, we generated a reaction-diffusion model describing how membrane-bound epidermal growth factor receptor (EGFR) activates the SH2 domain-containing protein tyrosine phosphatase SHP2 by assembling SHP2 complexes with the adaptor GRB2-associated binder 1 (GAB1). Src family kinases (SFKs) were included as a diffusible EGFR-activated intermediary kinase for GAB1. Contrary to the textbook picture of EGFR-mediated SHP2 activation, the model predicts that GAB1-SHP2 complexes can remain intact distal from membrane-retained EGFR at ∼80% of their plasma membrane concentration over the entire intracellular length scale. Parameter sensitivity analysis identified SFK inactivation as one of the most important kinetic processes controlling GAB1-SHP2 spatial persistence, revealing the key role of repeated adaptor phosphorylation in maintaining SHP2 activity. A simple order of magnitude analysis of the controlling rate processes supports the predictions of the coupled differential equation model. Furthermore, proximity ligation assays and immunofluorescence microscopy in carcinoma cells demonstrate that GAB1-SHP2 complexes are increasingly cytoplasmic and distal from EGFR as time proceeds after acute EGFR activation, consistent with model predictions. These results provide quantitative insight into a mechanism that may be alternately tuned for different receptors and downstream effector complexes to enable receptor control of signaling at a distance.

**STATEMENT OF SIGNIFICANCE:** The textbook understanding of receptor-mediated signaling involves the assembly of protein complexes at the receptor cytosolic tail and complex movement within the cell by protein trafficking. However, reaction-diffusion processes can also play an important role in distributing signaling proteins throughout the cell, potentially quite distal from signal-initiating receptors. Indeed, receptors can activate cytosolic kinases that may maintain the phosphorylation-dependent complexation of downstream effector complexes through many rounds of complex dissociation and protein dephosphorylation. Here, we develop quantitative understanding of this mode of signal regulation through development of a mechanistic computational model and demonstrate, with experimental validation, that membrane-bound receptors can regulate signaling through downstream effector complexes over the entire intracellular length scale.

## INTRODUCTION

Cell signaling by receptor tyrosine kinases (RTKs) involves the recruitment of SH2 and PTB domain-containing cytosolic adapter proteins to phosphorylated receptor cytoplasmic tyrosines. Phospho-dependent linkages between receptors and adapters and among proteins in larger complexes nucleated upon receptor activation are highly reversible, with low micromolar affinities and dissociation kinetics so rapid that complexes must reform repeatedly to sustain signaling^1, 2^. Protein tyrosine phosphatase (PTP) activity is also high enough that phosphotyrosines on constituent proteins can be appreciably dephosphorylated while not protected by SH2 domains^3, 4^. These effects may limit the access of functional, intact protein complexes to some intracellular locations, even though protein diffusion time scales are small (∼10 sec).

While some protein complexes form in direct association with signal-initiating receptors^5^, RTKs also activate cytosolic kinases that can promote the persistent complexation of proteins distal from the membrane. These effects can be hidden but present in co-immunoprecipitation experiments. Indeed, the presence of an RTK in the pulldown of a complex of downstream effectors does not necessarily indicate that all effector complexes are RTK-bound. However, the intracellular length scale over which intermediary RTK-activated kinases can maintain intact protein complexes and the most important determinants of that length scale are unknown. Here, we investigate this type of signaling regulation for the epidermal growth factor receptor (EGFR) and downstream protein complexes containing SH2 domain-containing phosphatase 2 (SHP2) and GRB2-associated binder 1 (GAB1).

SHP2 is encoded by the proto-oncogene *PTPN11* and is required for complete activation of ERK downstream of most RTKs^6–8^. SHP2 is basally catalytically inhibited by an intramolecular interaction between its N-terminal SH2 domain and its phosphatase domain^9^, but this tethering is relieved when SHP2 N- and C-terminal SH2 domains engage phosphorylated tyrosines including GAB1 Y627 and Y659^10^. Thus, GAB1-SHP2 complexes represent an active form of SHP2. In response to EGFR activation, GAB1 is phosphorylated by Src family kinases (SFKs), which reside both at the membrane and in the cytoplasm^10–12^. SHP2 binding to GAB1 has conventionally been viewed as a recruitment of cytosolic SHP2 to the plasma membrane through RTK-containing complexes such as EGFR-GRB2-GAB1-SHP2^13^ or through binding to PIP_3_-bound GAB1^14^. RTKs including Ret, HER2, and MET can also recruit SHP2 to the plasma membrane^15–17^. Interestingly, MET promotes more substantial SHP2 redistribution toward the plasma membrane than EGFR^18^. This difference may be based in part on the ability of EGFR to maintain cytosolic GAB1-SHP2 complexes by repeated SFK-mediated GAB1 phosphorylation^12^. Importantly, the ability for SFKs to phosphorylate cytosolic GAB1 will be influenced by the spatiotemporal regulation of SFK activity. SFKs can be activated at the plasma membrane in response to EGFR activation^19–23^, and they are inactivated throughout the cell by PTPs and c-SRC kinase-mediated phosphorylation of negative regulatory tyrosines on SFKs^24^. Based on this, SFK activity may rapidly decline distal from the plasma membrane.

Here, we develop a reaction-diffusion model to predict the temporal and spatial persistence of GAB1-SHP2 complexes in EGF-treated cells containing plasma membrane and cytosolic compartments. The fitted model predicts that GAB1-SHP2 complexes persist at nearly 80% of their plasma membrane concentration over the entire intracellular length scale. That is, EGFR in the plasma membrane can drive the activity of SHP2 throughout the cell interior. A parameter sensitivity analysis reveals that this complex persistence depends strongly on rates of SHP2-GAB1 association and dissociation, SFK inactivation, and GAB1 dephosphorylation. Model predictions are supported by a simple order of magnitude analysis of the competing reaction and diffusion processes and by experimental measurements based on proximity ligation assays and immunofluorescence microscopy. Our results demonstrate that cytosolic SFK activity enables EGFR to maintain active GAB1-SHP2 complexes throughout the intracellular domain and establish a quantitative framework for considering related phenomena in other receptor systems and cell settings.

## METHODS

### Model Development

#### General model description

Model equations comprise a system of coupled ordinary and partial differential equations for membrane and cytosolic species, respectively. Equations account for protein diffusion, reversible binding, and reversible tyrosine phosphorylation among EGFR, SFKs, GRB2, GAB1, and SHP2 in a spherical cell with radius *R*. Reaction rate expressions follow mass action kinetics. A model schematic is provided in **Fig. 1**, and model parameters with their baseline values (described below) are summarized in **Table 1**.

**Fig. 1.**
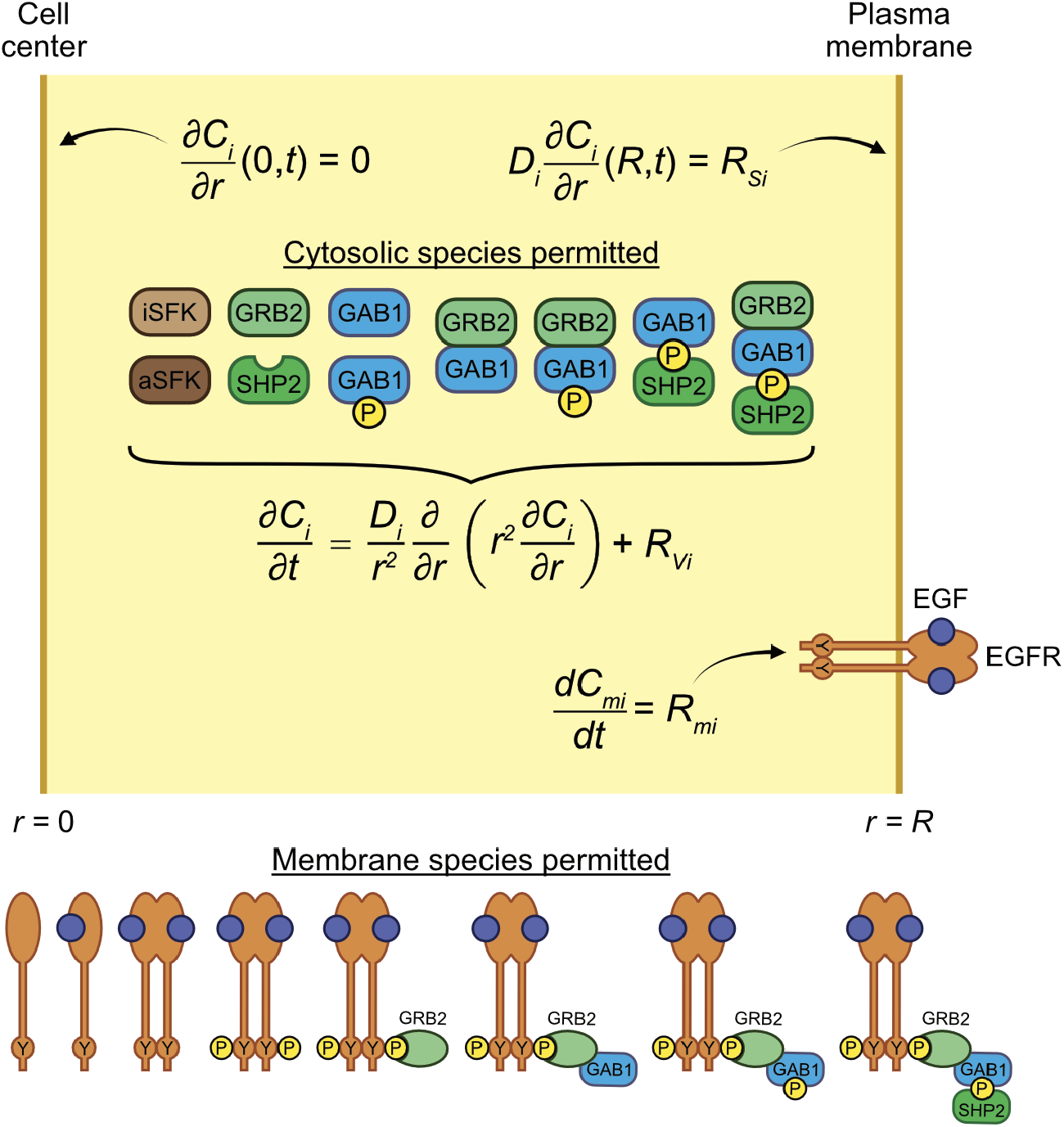
Model schematic. The cell cytosolic compartment is modeled as a sphere of radius *R* = 10 µm. The dependence of protein and protein complex concentrations on position and time after EGFR activation was modeled using standard reaction-diffusion equations. No-flux boundary conditions were imposed at the cell center consistent with symmetry of the problem, and Robin boundary conditions were imposed at the cell membrane to reflect the equivalence between reaction and diffusion of species at *r* = *R*. Membrane-bound species were assumed to be homogenously distributed in that compartment and were therefore modeled only as a function of time.

**Table 1.** Base model parameter values.

| Parameter (units) | Description | Base value |
| --- | --- | --- |
| $k_{E,f}$ ($\mu\text{M}^{-1} \text{min}^{-1}$) | EGF binding to EGFR, forward | 55.84 |
| $k_{E,r}$ ($\text{min}^{-1}$) | EGF binding to EGFR, reverse | 0.1301 |
| $k_{dE,f}$ ($\mu\text{m}^2 \text{molec}^{-1} \text{min}^{-1}$) | EGFR dimerization, forward | 1.2 |
| $k_{dE,r}$ ($\text{min}^{-1}$) | EGFR dimerization, reverse | 0.456 |
| $k_{catE}$ ($\text{min}^{-1}$) | EGFR phosphorylation, EGF-occupied dimer | 13.84 |
| $k_{dp}$ ($\text{min}^{-1}$) | EGFR dephosphorylation | 41.21 |
| $k_{S,a}$ ($\mu\text{m}^3 \text{molec}^{-1} \text{min}^{-1}$) | SFK activation | 0.7924 |
| $k_{S,i}$ ($\text{min}^{-1}$) | SFK inactivation | 4.67 |
| $k_{G2,f}$ ($\mu\text{m}^3 \text{molec}^{-1} \text{min}^{-1}$) | GRB2 binding to EGFR, forward | 1.594 |
| $k_{G2,r}$ ($\text{min}^{-1}$) | GRB2 binding to EGFR, reverse | 480 |
| $k_{G1,f}$ ($\mu\text{m}^3 \text{molec}^{-1} \text{min}^{-1}$) | GAB1 binding to GRB2, forward | $2.4 \times 10^3$ |
| $k_{G1,r}$ ($\text{min}^{-1}$) | GAB1 binding to GRB2, reverse | $3.9 \times 10^{-2}$ |
| $k_{G1p}$ ($\mu\text{m}^3 \text{molec}^{-1} \text{min}^{-1}$) | GAB1 phosphorylation | 1.267 |
| $k_{G1dp}$ ($\text{min}^{-1}$) | GAB1 dephosphorylation | 3.118 |
| $k_{S2,f}$ ($\mu\text{m}^3 \text{molec}^{-1} \text{min}^{-1}$) | SHP2 binding to phosphorylated GAB1, forward | 1.594 |
| $k_{S2,r}$ ($\text{min}^{-1}$) | SHP2 binding to phosphorylated GAB1, reverse | 480 |
| $D_{SFK}$ ($\mu\text{m}^2 \text{min}^{-1}$) | Diffusivity of SFK molecules | 84 |
| $D_{G2}$ ($\mu\text{m}^2 \text{min}^{-1}$) | Diffusivity of GRB2 molecules | 136 |
| $D_{G2G1}$ ($\mu\text{m}^2 \text{min}^{-1}$) | Diffusivity of GRB2-GAB1 molecules | 62 |
| $D_{G2G1S2}$ ($\mu\text{m}^2 \text{min}^{-1}$) | Diffusivity of GRB2-GAB1-SHP2 molecules | 56 |
| $D_{G1}$ ($\mu\text{m}^2 \text{min}^{-1}$) | Diffusivity of GAB1 molecules | 67 |
| $D_{G1S2}$ ($\mu\text{m}^2 \text{min}^{-1}$) | Diffusivity of GAB1-SHP2 molecules | 57 |
| $D_{S2}$ ($\mu\text{m}^2 \text{min}^{-1}$) | Diffusivity of SHP2 molecules | 80 |
| $C_{o,EGF}$ ($\mu\text{M}$) | Extracellular EGF concentration | $1.7 \times 10^{-3}$ |
| $C_{o,EGFR}$ ($\text{cell}^{-1}$) | EGFR molecules per cell | $6.0 \times 10^5$ |
| $C_{o,GRB2}$ ($\text{cell}^{-1}$) | GRB2 molecules per cell | $6.0 \times 10^5$ |
| $C_{o,GAB1}$ ($\text{cell}^{-1}$) | GAB1 molecules per cell | $6.0 \times 10^5$ |
| $C_{o,SHP2}$ ($\text{cell}^{-1}$) | SHP2 molecules per cell | $6.0 \times 10^5$ |
| $C_{o,SFK}$ ($\text{cell}^{-1}$) | SFK molecules per cell | $6.0 \times 10^5$ |
| $R$ ($\mu\text{m}$) | Cell radius | 10 |

Within the cytosolic compartment, conservation of species *i* (*C_i_*) was modeled as

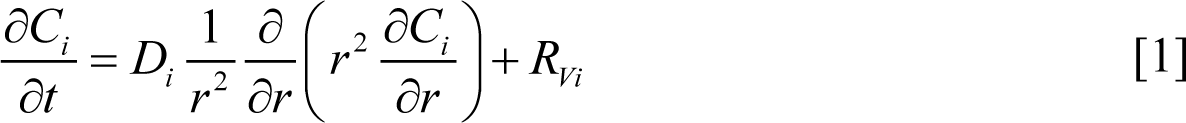

where *D_i_* is the diffusivity and *R_Vi_* is the net volumetric rate of generation. No-flux (symmetry) boundary conditions were implemented at the cell center for all species such that

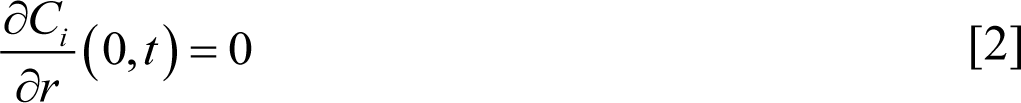

Robin boundary conditions were applied at the membrane to equate the diffusive flux from the membrane (into the cytosol) to the rate of generation of species *i* (*R_si_*) at the membrane. Thus,

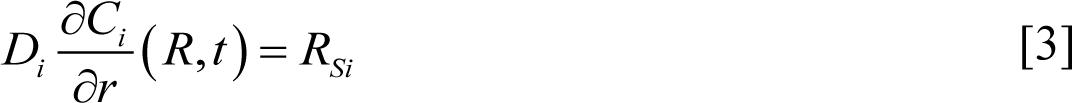

Membrane species were treated as uniformly distributed in the membrane and modeled only as a function of time as

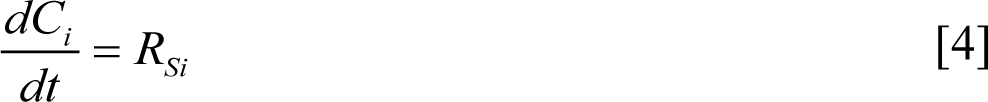

#### Finite difference solution

Partial differential equations were discretized using an explicit finite difference method, wherein first- and second-order derivatives were approximated using forward and central differences, respectively. At each time point, the system of algebraic equations resulting from discretizing over the entire spatial domain was explicitly solved using concentrations from the prior time point. The coupled equations for membrane proteins and boundary conditions for cytosolic proteins were simultaneously solved using the semi-implicit Euler method. Discretized boundary conditions for cytosolic proteins were evaluated first, using initial guesses for membrane protein concentrations at the next time step. Subsequently, discretized conservation equations for membrane proteins were evaluated using cytosolic protein concentrations from the explicit finite difference calculation. The membrane protein concentrations calculated in this step were then used as initial guesses for the boundary conditions in the next iteration of the semi-implicit scheme. This process was repeated iteratively (up to a maximum of 100 iterations) until the fractional error between the previous and updated concentrations for membrane species and cytosolic proteins at the boundary was < 1×10^-6^. The process above was then successively repeated for each time point until the end of the time domain considered was reached. Additional details are provided in Appendix S1.

#### Method of lines solution

To overcome stability issues of the explicit finite difference scheme encountered in some model sensitivity analysis calculations, during which the time step required to maintain solution stability becomes prohibitively small (primarily due to large values of diffusivities), the model was also solved by converting it into a system of time-dependent ordinary differential equations (ODEs) using the method of lines^25^. This was performed by symbolically defining the model equations and boundary conditions using ModelingToolkit.jl^26^, discretizing them in the spatial dimension and converting to a system of ODEs using MethodOfLines.jl (https://github.com/SciML/MethodOfLines.jl), and solving the system with DifferentialEquations.jl^27^ using the QNDF stiff ODE solver. The model implemented in this way was used to perform eFAST sensitivity analysis calculations involving kinetic parameters and protein diffusivities (see below).

#### Simplified, steady-state model

A simplified steady-state model was developed by limiting the role of EGFR to SFK activation only (i.e., no adaptor binding) and setting the concentration of phosphorylated EGFR to the value predicted by the full model for the 5-min response to EGF. Solving the conservation equations for active and inactive SFKs analytically, the *r*-dependent concentration of active SFKs is

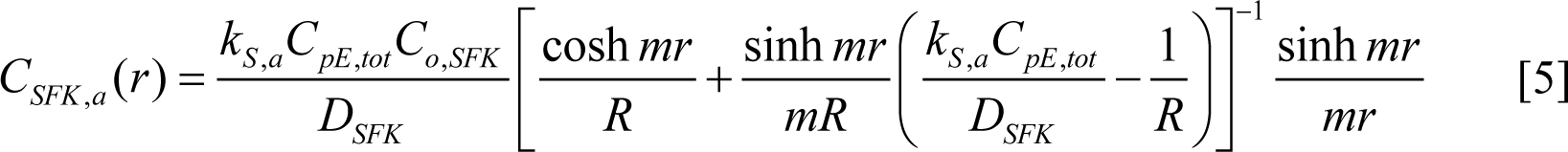

where *C_pE_* is the concentration of phosphorylated EGFR, and other parameters are as defined in **Table 1**. The concentration of inactive SFKs is *C_SFK,i_* = *C_o,SFK_* – *C_SFK,a_*, where *C_o,SFK_* is the initial concentration of SFKs.

Without binding to EGFR and GRB2 at the cell membrane, the boundary conditions for the four remaining cytosolic species (GAB1, pGAB1, pGAB1-SHP2, and SHP2) are of the form

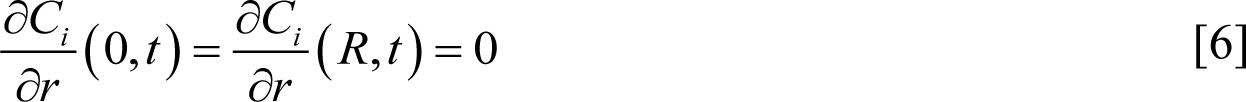

Subject to Eq. 6, the conservation equations for these species were solved using a modified, steady-state finite difference method without time steps.

To create a further simplified model, the governing equations for GAB1, pGAB1, pGAB1-SHP2, and SHP2 were reduced to just two equations for pGAB1 and pGAB1-SHP2 using conserved scalar quantities. The resulting equations were solved using the modified finite difference scheme. Finally, the system was reduced yet further to a single conservation equation for pGAB1 by assuming that GAB1 and SHP2 binding was everywhere at equilibrium, which enabled the elimination of all other species from the differential equation by substitution. The resulting conservation equation for pGAB1 was also solved by modified finite difference. Appendix S2 provides additional details on model simplifications, governing equations, and the modified steady-state finite difference approach.

### Model Implementation

#### General considerations

Model codes were written and compiled primarily in MATLAB R2020b (The MathWorks, Natick, MA) and Julia v1.8.3^28^ and are available on GitHub under the username “Lazzara Lab” (https://github.com/lazzaralab). In total, the model consists of 18 distinct protein species (10 cytosolic and eight membrane) and 25 reactions. Model parameters are described in **Table 1**. For all finite difference calculations, except those for sensitivity analyses, the spatial dimension was discretized using a step = 0.1 µm. For all sensitivity analyses, whether using the explicit finite difference or method of lines schemes, a spatial step = 0.2 µm was used to increase calculation efficiency. For all explicit finite difference calculations, the time step was set based on the explicit finite difference stability criterion described in Eq. 8 by Bieniasz^29^ for diffusion problems with homogeneous reactions. Typical time steps ranged between ∼3×10^-4^ sec to ∼3×10^-6^ sec, depending on the values of model parameters and the spatial step size.

An additional version of the model was developed in Virtual Cell (VCell)^30, 31^ to provide accessibility to VCell users. The VCell model “Myers&Furcht_spatial_GAB1_SHP2” was constructed as a BioModel in VCell 7.5 and is available in the public domain through the Public BioModels portal in VCell and at http://vcell.org/vcell-models under the shared username “LazzaraLab.” Solution geometries were constructed in VCell by defining the cytoplasmic volume compartment as a sphere with a radius of 10 µm or 100 µm.

The cell radius *R* was set equal to 10 μm^32^ for most calculations. When the model was used to compute length scales for species persistence that could exceed 10 μm (e.g., with perturbed parameters), *R* was set to 100 μm. For most calculations, EGFR, GRB2, GAB1, SHP2, and SFKs were assumed to be at cellular concentrations of 6×10^5^ per cell, as described in a previous, purely dynamic model^12^. However, in some calculations, stoichiometries based on absolute quantification of protein expression in HeLa cells were used^33^. Total concentrations of EGFR, SFKs, GRB2, GAB1, and SHP2 were converted to units of molecules/µm^3^ using the appropriate volume and surface area conversion factors such that the total number of molecules of each protein per cell did not change between calculations whether the cell radius was set to 10 or 100 μm.

#### Kinetic and diffusion processes considered

EGF binding at the plasma membrane was modeled as a reversible process characterized by association and dissociation rate constants^34^. EGF was treated as present at a constant concentration of 10 ng/mL, which is approximately equal to EGF’s affinity for EGFR. The EGFR dimerization rate constant was used as calculated previously^35^, and the dimer uncoupling rate constant in the presence of EGF was calculated using this forward dimerization rate constant and a previously estimated dissociation constant for EGF-bound EGFR dimers^36^. All dimer species were assumed to be symmetric, i.e., no coupling of ligand-bound and -unbound EGFR monomers. EGFR phosphorylation was modeled as occurring between EGF-bound EGFR receptors in a dimer, where both receptors are simultaneously phosphorylated at a representative tyrosine (Y1068) that binds GRB2. EGFR dephosphorylation was modeled as zeroth-order with respect to protein tyrosine phosphatases based on previously described rate constants^12, 37^.

As in prior work^12, 38^, SFK activation was modeled as a first-order process wherein phosphorylated EGFR activates SFK at the plasma membrane. This approach is based on knowledge that Src SH2 domains can bind EGFR^39^ and that Src SH2 engagement promotes autophosphorylation and activation of Src^40^. SFK inactivation was modeled as zeroth order volumetric process occurring throughout the cytosol.

GAB1 phosphorylation at a representative tyrosine that binds SHP2 (e.g., Y627) was modeled as a process catalyzed by active SFKs throughout the cytosol. GAB1 dephosphorylation was modeled as a zeroth order volumetric reaction in the cytosol. Binding interactions among GRB2, GAB1, and SHP2 were modeled as described previously^12^.

GRB2 was modeled as being able to bind phosphorylated EGFR using experimentally derived rate constants for association and dissociation^2^. Lacking direct measurements of SHP2 binding to phosphorylated GAB1, the association and dissociation rate constants for GRB2-EGFR were used to describe GAB1-SHP2 binding, assuming SHP2 follows similar phosphotyrosine– SH2 domain–mediated interactions to GAB1 as GRB2 has with EGFR. GAB1 binding to GRB2 was modeled as occurring between GAB1’s proline-rich domain (PRD) and GRB2’s SH3 domain. Again lacking direct measurements of this interaction, association and dissociation rate constants for GAB1-GRB2 binding were estimated based on experimental measurements for GRB2 and c-Src SH3 domains binding to the PRDs of dynamin-1/2^41^.

Diffusivities for cytosolic proteins and complexes were estimated based on the diffusivity of tubulin^42^ and an adjustment factor based on differences in the hydrodynamic radii of tubulin and a given protein monomer or complex. The hydrodynamic radii of protein monomers and complexes were estimated based on their molecular weights^43^. See Appendix S4 for details.

#### Generation of fixed and prior parameter distributions

Using reported parameter values and uncertainties (if available) obtained from the literature sources described above, distributions of probable values were generated for each model parameter according to the protocol described by Tsigkinopoulou et al.^44^. For processes involving forward and reverse rates (e.g., EGF binding to EGFR, GRB2-GAB1 binding, etc.), parameters were modeled using multivariate log-normal distributions to maintain thermodynamic consistency. All other parameters were modeled using univariate log-normal distributions. This approach yielded a distribution of plausible values for each parameter, except for the parameters describing SFK activation/inactivation (*k_S,a_*, and *k_S,i_*) and GAB1 phosphorylation/dephosphorylation (*k_G1p_* and *k_G1dp_*); to maximize the model’s ability to recapitulate experimental measurements of GAB1-SHP2 association, prior and posterior distributions of plausible values for these four parameters were determined through model fitting, as described in the next section. The distributions generated in this step for all other (non-fitted) parameters were fixed and used in all subsequent model calculations. The parameter values and uncertainties used to generate parameter distributions are reported in **Table S1**, and the distributions generated from these values are shown in **Fig. S1A**. As advocated by Tsigkinopoulou et al.^44^, as many appropriate parameter estimates as possible were used from the literature to generate the distributions for the fixed model parameters. For example, experimental measurements and associated uncertainties of the rates of phosphorylation for multiple intracellular EGFR tyrosine residues were used to estimate an overall EGFR phosphorylation rate constant for use in model calculations. In cases where parameter uncertainties were unavailable, multiplicative uncertainties of 10% of the reported parameter value were assumed^44^. Custom Julia scripts used to generate these distributions were adapted from the original MATLAB scripts written by Tsigkinopoulou et al.^44^ and are available with the rest of the model codes on GitHub.

#### Parameter fitting and ensemble generation using Bayesian inference

The values for the parameters for *k_S,a_*, *k_S,i_*, *k_G1p_* and *k_G1dp_* were obtained by fitting model predictions for the fraction of SHP2-bound GAB1 after 5 min of 10 ng/mL EGF treatment to experimental measurements from EGF-treated H1666 cells^12^. Rather than computing a single best-fit parameter set, Bayesian inference was used to generate distributions of parameter values consistent with the uncertainty in the experimental data and other model parameters. As described in detail below, this was accomplished by first obtaining a best-fit point estimate for these four fitted parameters, followed by using this estimate to generate the prior distributions needed for Bayesian inference calculations. The result of the inference process is a set of values (posterior distribution) for each parameter that is consistent with the fitting data and its associated uncertainty. By taking simultaneous draws from these posterior distributions and the distributions of the fixed parameters described above, an ensemble of parameter values is obtained that can be used for model simulations. Using ensemble model calculations, credible intervals (estimates of model prediction uncertainty) were obtained for the majority of model calculations except in certain cases where this was impractical or unhelpful (see the following section, *Definition of “base” parameter values*, for additional detail).

The initial fit for the four fitted parameters was obtained by minimizing the sum-of-squares error between model and experiment for the fraction of SHP2-bound GAB1 using the Optimization.jl meta-package (https://github.com/SciML/Optimization.jl). The upper and lower bounds for *k_S,a_* and *k_G1p_* were set by multiplying and dividing the value for *k_G1p_* from a prior model^12^ by a factor of 100; the bounds for *k_S,i_* and *k_G1dp_* were similarly set based on the value for *k_G1dp_* from the same prior model. These bounds were set based on the values for *k_G1p_* and *k_G1dp_* from the prior model due to topological differences in the descriptions of SFK inactivation between that model and the reaction-diffusion model here. The values of all other parameters during initial model fitting were taken to be the modes from their respective parameter distributions (**Fig. S1A;** see also the values for these parameters reported in **Table 1**). 100 optimization start sites (sets of initial guesses) were generated within these bounds using the TikTak multi-start optimization algorithm^45^ from the MultistartOptimization.jl package (https://github.com/tpapp/MultistartOptimization.jl). Optimization was performed for each start site using the limited-memory Broyden-Fletcher-Goldfarb-Shanno algorithm (L-BFGS)^46, 47^. L-BFGS was accessed with NLopt.jl, a Julia interface to the NLopt library (http://github.com/stevengj/nlopt). Optimization efficiency was improved by computing the gradient of the objective function using forward-mode automatic differentiation with ForwardDiff.jl^48^. The best-fit parameter values were taken from the optimization run with the minimum error. These values are included in **Table S1** as the initial values used to generate the prior distributions for these four parameters that are needed to perform Bayesian inference.

After generating initial point estimates, Bayesian inference was then performed for *k_S,a_*, *k_S,i_*, *k_G1p_*, and *k_G1dp_* using Turing.jl, a probabilistic programming language written in Julia^49^. The same fitting data used to generate the initial point estimates were assumed to follow a log-normal distribution with variance calculated based on the experimental measurement uncertainty. Log-normal prior distributions for *k_S,a_*, *k_S,i_*, *k_G1p_*, and *k_G1dp_* were centered around the best-fit point estimates described above with an assumed multiplicative uncertainty of 10% of their values, as suggested by Tsigkinopoulou et al.^44^. The uncertainty in other model parameters was incorporated in the inference process by drawing from their fixed distributions (generated as described in the preceding section) during posterior sampling for *k_S,a_*, *k_S,i_*, *k_G1p_*, and *k_G1dp_*. Inference was performed with the No U-turn sampler (NUTS), an adaptive Hamiltonian Monte Carlo method, using the recommended acceptance rate of 0.65^50^. 5,000 posterior samples (parameter sets) were generated with NUTS by obtaining 1,000 samples each from 5 independent chains. Chain convergence was assessed graphically and using the Gelman-Rubin convergence diagnostic^51, 52^. The ensemble of model fits accurately recapitulates the experimentally observed fraction of SHP2-bound GAB1 in response to EGF (**Fig. S1B**) when sampling from the generated parameter posteriors (**Fig. S1C**). Furthermore, the average net rates of SFK activation and GAB1 phosphorylation are consistent with experimentally measured phosphorylation rates for multiple EGFR substrates^53^ and are thus within the range with physically plausible values for kinase substrate phosphorylation/activation.

#### Definition of base parameter values

For certain calculations in which full ensemble predictions were impractical or uninformative, as noted specifically in figure legends or the text, model predictions were made using a single set of baseline or base parameter values. These were defined as the modes (most probable values) of the fixed (**Fig. S1A)** and posterior distributions (**Fig. S1C**) for the fixed and fitted model parameters, respectively. These were also used as the baseline values around which parameters were allowed to vary as part of model sensitivity analysis calculations. The base values of each model parameter and the base abundances for each model species are described in **Table 1**.

#### Model sensitivity analysis

Model sensitivity to simultaneous changes in all parameter values was computed using the extended Fourier amplitude sensitivity test (eFAST)^54, 55^. eFAST calculates first-order (S_1_) and total-order sensitivity indices (S_T_) for each input parameter by simultaneously varying all parameters between manually specified upper and lower bounds according to sine waves with distinct frequencies and using Fourier decomposition to calculate the variance in model output explained by the input parameters. S_1_, also known as the main-effect index, accounts for the fraction of overall model output variance that can be attributed solely to an individual parameter, whereas S_T_ indicates the fraction of the overall model output variance that can be attributed to a parameter both individually and when accounting for all possible interactions with the other model parameters being varied. Thus, the variance in model output that can be attributed solely to interactions of an input parameter with other model parameters is calculated as the difference S_T_– S_1_. eFAST calculations were performed with the Julia version of the model using the GlobalSensitivity.jl package^56^. Diffusivities and kinetic parameters were varied by a factor ≤ 1,000 above or below their base values, as described in **Table 1**. For the fitted parameters these values are taken as their modes from the full set of 5,000 posterior samples. Protein concentrations were allowed to vary between 100 and 1,000,000 molecules per cell. In all eFAST calculations, 1,000 simulations were run per varied model parameter.

### Length-scale arguments for active SFKs and GAB1-SHP2 complexes

Estimates for the length scales over which concentrations of active SFKs and GAB1-SHP2 complexes change were developed by considering the orders of magnitude of the rate processes involved in the reaction-diffusion mechanism. The length scale obtained for GAB1-SHP2 was assumed to apply to GRB2-GAB1-SHP2 because the diffusivities of these two species are nearly identical (**Table 1**) and GRB2 binding does not prevent GAB1-SHP2 association in the current model topology. See Appendix S3 for additional details regarding these derivations.

The length scale of active SFKs can be derived from the steady conservation equation for *C_SFK,a_*, which is

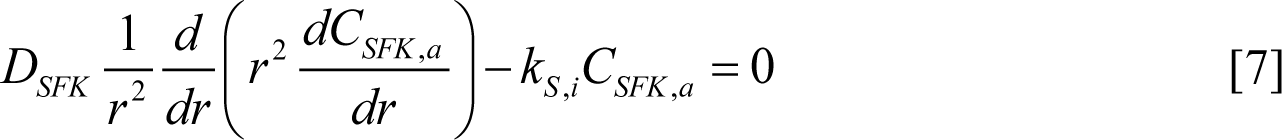

Note that *R_Vi_* for active SFKs does not include a forward (activation) rate because SFK activation occurs only at the plasma membrane. Dimensionless *C_SFK,a_* and *r* are defined as

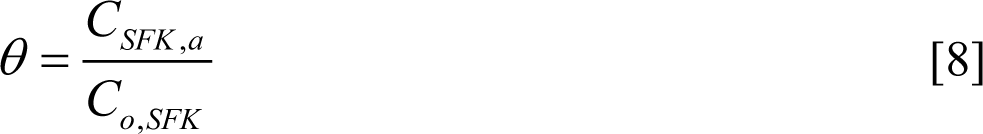

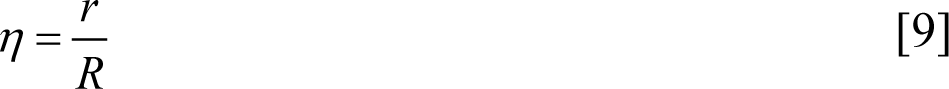

Substituting these definitions into Eq. 7 yields

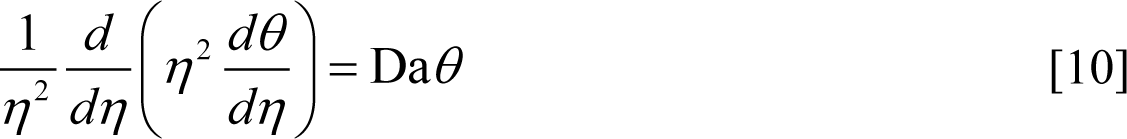

where the Damköhler number Da = *k_S,i_R*^2^/*D_SFK_* is a dimensionless quantity describing the competition between SFK inactivation and SFK diffusion.

Based on the parameter values in **Table 1**, Da ≈ 6. Thus, the inactivation of SFKs occurs rapidly compared to SFK diffusion over the length scale *R*, and a sharp gradient in *C_SFK,a_* is expected. The large value of Da also indicates that Eq. 10 is not properly scaled (i.e., not all terms are ∼1), which is a consequence of the geometric length *R* (used to define *η*) overestimating the length scale for changes in *C_SFK,a_*. Redefining the radial coordinate by incorporating Da as

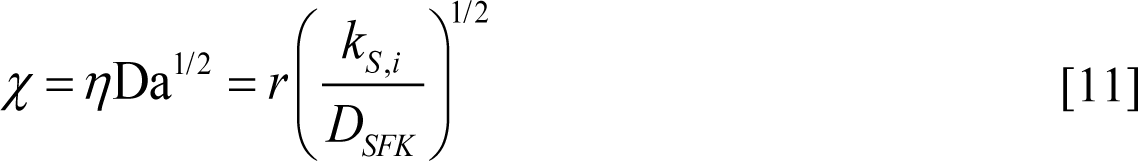

eliminates Da from the conservation equation, resulting in this scaled result:

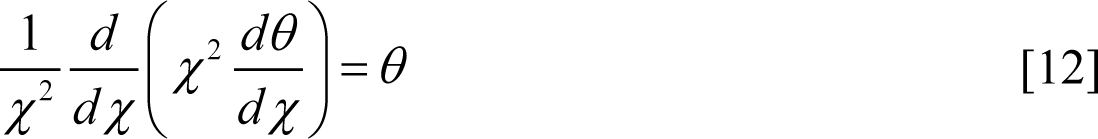

Based on Eq. 11, the length scale for changes in *C_SFK,a_* (*δ_SFK_*_,*a*_) is

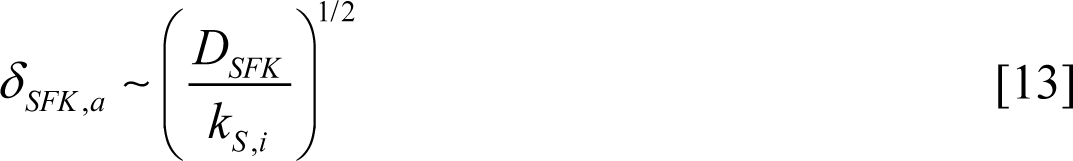

where “∼” denotes order-of-magnitude equality.

While *δ_SFK_*_,*a*_ is readily derived through standard scaling analysis^57^, the length scale for GAB1-SHP2 complex persistence (*δ_G_*_1*S*2_) is more challenging to obtain due to the coupled kinetic processes involved. An alternative approach is to estimate and sum the length scales for the processes that cooperate to determine *δ_G_*_1*S*2_: i. SFK-mediated GAB1 phosphorylation, ii. GAB1-SHP2 dissociation, and iii. GAB1 dephosphorylation. This approach recognizes that complexes will form wherever GAB1 is actively being phosphorylated, that intact complexes require a finite time to dissociate, and that complexes can reform where GAB1 remains phosphorylated.

The length scale for SFK-mediated GAB1 phosphorylation is just *δ_SFK_*_,*a*_. Based on the same reasoning used to derive *δ_SFK_*_,*a*_, the length scale for GAB1-SHP2 dissociation (*δ_dis_*) is

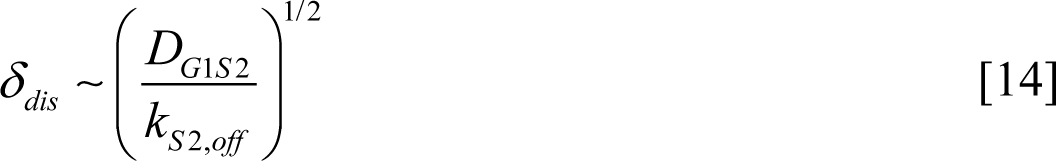

and the length scale for GAB1 dephosphorylation (*δ_dep_*) is

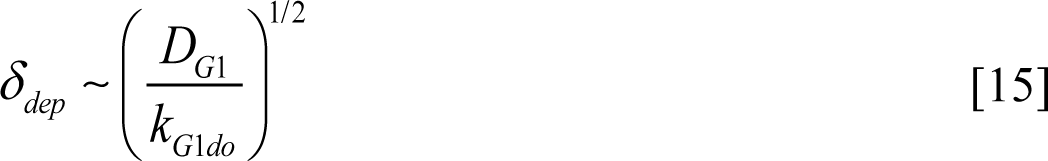

Thus, *δ_G_*_1*S*2_ can be estimated as

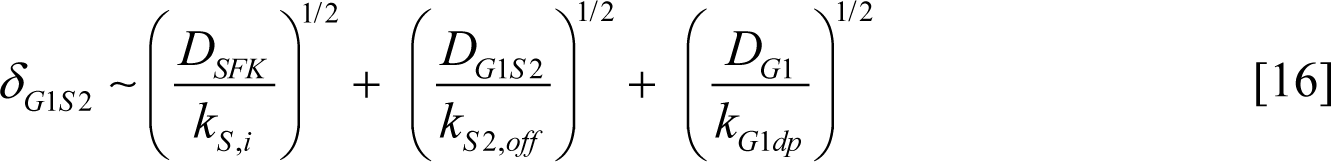

### Experimental Methods

#### Cell culture

Gene-edited HeLa cells expressing mVenus-HRAS (HeLa/mV-HRAS) from a single endogenous *HRAS* locus were described previously^58^. Unless otherwise noted, cells were cultured in Dulbecco’s Modified Eagle’s Medium (DMEM) supplemented with 10% fetal bovine serum (FBS) (Avantor, Radnor, PA), 1% penicillin-streptomycin (10,000 µg/mL), and 1% L-glutamine (200 mM). Cells were confirmed to be mycoplasma-negative using the MycoAlert Mycoplasma Detection Kit (LT07-318; Lonza, Basel, Switzerland).

#### Proximity ligation assay and immunofluorescence

HeLa/mV-HRAS cells were plated on 12-mm coverslips in six-well tissue culture dishes at 3×10^5^ cells per well. ∼24 hr after plating, cells were serum-starved by switching them to DMEM containing 0.1 % FBS for at least 16 hr. Serum-starved cells were treated with recombinant human EGF (AF-100-15; PeproTech, Cranbury, NJ) at 10 ng/mL for ≤ 15 min. Medium was then aspirated, and cells were fixed in 4% paraformaldehyde in PBS for 20 min, followed by permeabilization with 0.25% Triton X-100 in PBS for 5 min. Coverslips were blocked in Duolink Blocking Solution (Sigma Aldrich, Burlington, MA) for 1 hr at 37°C and were then incubated with primary antibodies overnight at 4°C. All antibodies were diluted in Odyssey Blocking Buffer (OBB; LI-COR Biosciences, Lincoln, NE). For GAB1-SHP2 proximity ligation assays (PLA), mouse SHP2 antibody (sc-7384, 1:50; Santa Cruz Biotechnology, Santa Cruz, CA) and rabbit GAB1 antibody (HPA049599, 1:50, Sigma-Aldrich) were used. For EGFR-GRB2 PLA, mouse EGFR antibody (CST D381B, 1:100; Cell Signaling Technology, Danvers, MA) and rabbit GRB2 antibody (BD 610111, 1:100; BD Biosciences, Franklin Lakes, NJ) were used. PLA probes were anti-rabbit PLUS (DUO92002, Sigma-Aldrich) and anti-mouse MINUS (DUO92004, Sigma-Aldrich) and were used and detected using Duolink In Situ Far Red Detection Reagents (Sigma-Aldrich) according to the manufacturer’s instructions. After staining with PLA probes, coverslips were incubated overnight at 4°C with rat EGFR antibody (ab231, 1:200; Abcam, Cambridge, United Kingdom). EGFR was detected by incubating coverslips for 1 hr at 37°C with anti-rat Alexa Fluor 594 (A-11007, 1:750; Invitrogen, Waltham, MA) and Hoechst nuclear stain (1:2,000). Coverslips were mounted on glass slides using ProLong Gold Antifade Mounting Medium (P36930; Thermo Fisher Scientific, Waltham, MA). Epifluorescence images were obtained with a Zeiss AxioObserver Z1 using 20×, 40× oil immersion, and 63× oil immersion objective lenses.

Fluorescence images were analyzed using CellProfiler (version 4.2.5)^59^ and assorted third-party CellProfiler plugins (https://github.com/CellProfiler/CellProfiler-plugins). The CellProfiler pipelines used in this work are available on GitHub (https://github.com/lazzaralab) in the same repository containing the PDE model codes. Quantification of PLA puncta and colocalization of PLA and EGFR puncta was performed using an approach based on the protocols described by Horzum et al.^60^ and Moser et al.^61^. Briefly, maximum-intensity z-projections of PLA and EGFR immunofluorescence (IF) images were corrected for uneven illumination and background using the CorrectIlluminationCalculate and CorrectIlluminationApply modules. The Laplacian of Gaussian (LoG) filter was applied to illumination-corrected IF and PLA images to enhance individual puncta. Puncta in LoG-filtered images were further enhanced using the EnhanceOrSuppressFeatures module and thresholded using the IdentifyPrimaryObjects module to identify individual PLA and EGFR puncta. PLA and EGFR primary objects were related with the RelateObjects module to determine signal colocalization. Nuclear stain images were Gaussian-filtered to reduce background noise, followed by thresholding to identify individual nuclei as primary objects. Cell bodies were identified from mVenus-HRAS images by first adjusting image contrast with the HistogramEqualization module, followed by noise and background reduction with the ReduceNoise module. After noise reduction, mVenus-HRAS images were thresholded with the IdentifySecondaryObjects module to identify cell bodies as secondary objects of nuclear primary objects. PLA objects were assigned to individual cells based on overlap with mVenus-HRAS secondary objects calculated using the RelateObjects module. The distances of individual PLA objects to the nearest primary EGFR object and the nearest edge of the enclosing mVenus-HRAS secondary object (i.e., the plasma membrane) was estimated using the MeasureObjectNeighbors module.

### Statistical analyses and plotting

The ggstatsplot R package^62^ was used to perform the Kruskal-Wallis test and the Dunn post-hoc test. Bayes factors comparing model ensemble predictions were calculated using the BayesFactor R package^63^ and interpreted using the effectsize R package^64^. The R plotting library ggplot2^65^ and the Julia plotting libraries Makie.jl^66^ and AlgebraOfGraphics.jl were used throughout for figure generation.

## RESULTS

### Base Model Predictions

As detailed in *Methods*, the model accounts for EGFR phosphorylation, SFK activation, GAB1 phosphorylation, and GAB1-SHP2 complex formation in a spherical cytosolic compartment bounded by a plasma membrane (**Fig. 1**). Unless otherwise stated, the concentration of GAB1-SHP2 complexes is calculated as the sum of the species pGAB1-SHP2 and GRB2-pGAB1-SHP2. Simulations were limited to a 5-min response to EGF because peak EGFR activation and adapter binding typically occur within this time^67, 68^ and because EGFR endocytosis and degradation—processes that are computationally prohibitive and difficult to accurately model spatiotemporally with available modeling tools—are prominent after 5 min^5, 58, 69, 70^. Thus, model calculations provide a conservative estimate of the ability of the membrane-localized receptors to regulate signaling via SHP2 throughout the cytoplasmic domain.

Ensemble calculations with the full model predict that SFKs rapidly activate at the plasma membrane in response to EGF but that the concentration of active SFKs decays substantially to about 20% of the membrane concentration over a length scale that is small compared to the cell radius, though for some ensemble predictions they decay to nearly zero at *t* = 5 min (**Fig. 2A**; note the dashed red lines at *t* = 5 min). In contrast, the concentration of pGAB1 is nearly constant moving away from the membrane and toward the cell interior (**Fig. 2B**). The difference between the length scales for active SFKs and pGAB1 decay is related in part to the time required for GAB1 dephosphorylation, an issue examined in greater detail later. The difference in length scales is also related to an effective signaling amplification step from EGFR to GAB1, wherein ∼10-20 pGAB1 molecules result via SFKs for every pEGFR (**Fig. 2C**). As with pGAB1, the abundance of GAB1-SHP2 complexes is largely unchanged as a function of the radial coordinate *r*, even with uncertainty in the overall complex concentration across model ensemble predictions, suggesting that SHP2 remains active over the entire intracellular length scale when activated by membrane-retained EGFR (**Fig. 2D**). The predicted response of GAB1-SHP2 complexes to treatment with the EGFR inhibitor gefitinib after an initial pulse of EGF demonstrated a slower decay of GAB1-SHP2 complexes (∼2 min; **Fig. 2E**) than the rate of initial complex formation (∼30 sec; **Fig. 2D**). The slower rate of decay is the result of the time scale required for GAB1 dephosphorylation, which cannot occur until the effect of EGFR inactivation propagates to an inactivation of SFKs. The finite time scale for this propagation explains why the predicted loss of EGFR activity occurs more quickly than does the loss of GAB1-SHP2 complexes.

**Fig. 2.**
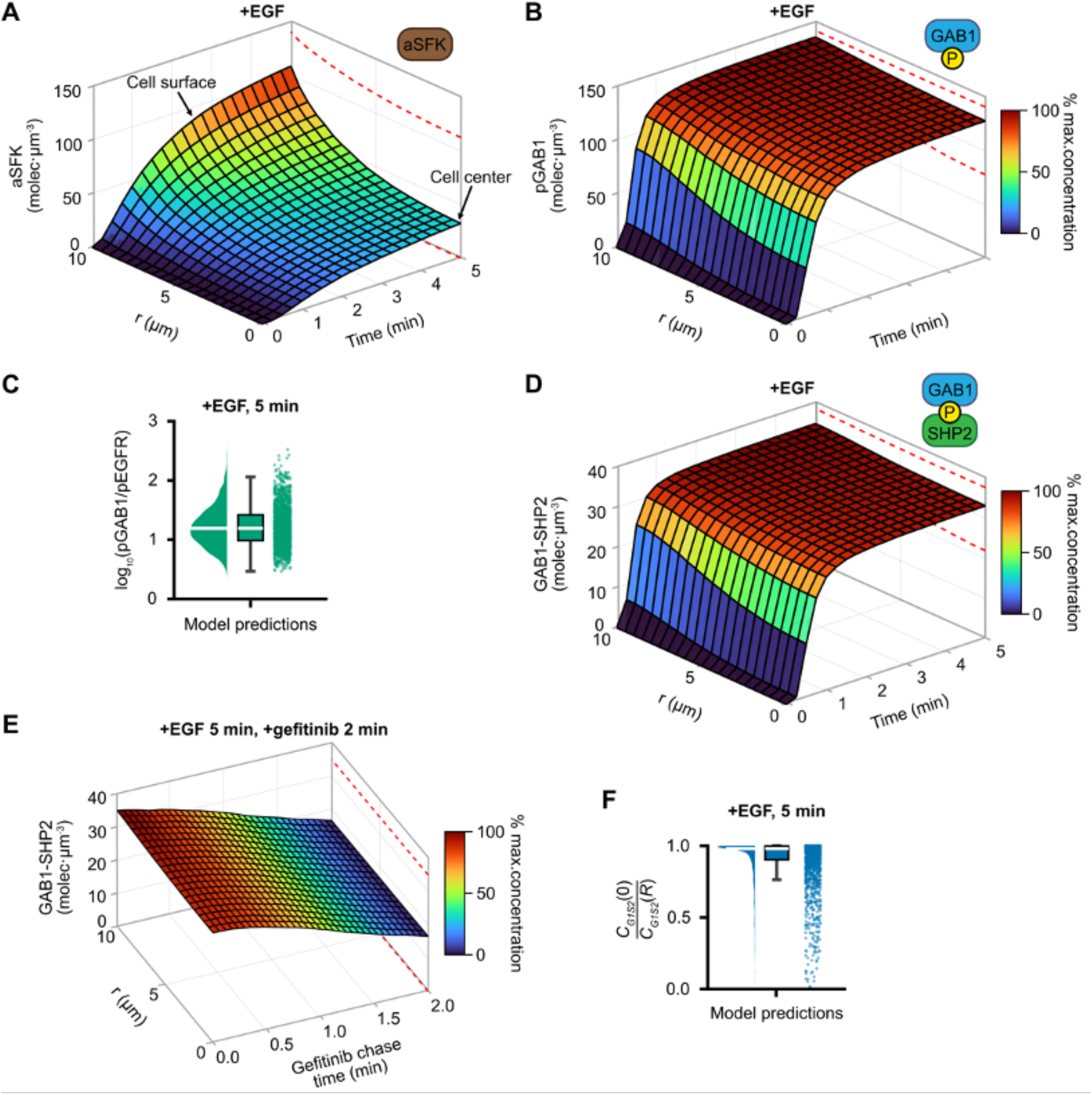
GAB1-SHP2 complexes are well distributed throughout cells in response to EGFR activation. **(A-B)** Model predictions are shown for the concentrations of active SFKs and phosphorylated GAB1 (pGAB1) in response to 10 ng/mL EGF as a function of distance from the cell center (*r*) and time for an ensemble of 2,000 draws from model parameter distributions. Surfaces represent the median of ensemble predictions at each space-time node, and dashed red lines denote the 68.2% credible interval for ensemble predictions centered around the median at *t* = 5 min. (**C**) The distribution for the ratio of pGAB1 and phosphorylated EGFR (pEGFR) in response to 10 ng/mL EGF at *t* = 5 min was computed using 2,000 draws from model parameter distributions and plotted as a raincloud plot. Individual model simulations are shown as dots at right, and the distribution of the simulations is described by the box and half-violin plots at left. (**D**) Model predictions are shown for the concentration of GABl-SHP2 complexes in response to 10 ng/mL EGF as a function of *r* and time for 2,000 draws from model parameter distributions. The surface represents the median of ensemble predictions, and dashed red lines enclose the 68.2% credible interval centered around the median at *t* = 5 min. (**E**) For 2,000 draws from model parameter distributions, the concentration of GAB1-SHP2 complexes was predicted as a function of *r* and time for an initial 5-min pulse of 10 ng/mL EGF chased by a 2-min treatment with the EGFR inhibitor gefitinib. Gefitinib treatment was simulated by setting the rate constant for EGFR phosphorylation to 0. Model predictions are shown for the 2-min chase time course. The surface represents the median of ensemble predictions, and dashed red lines enclose the 68.2% credible interval centered around the median at *t* = 5 min. (**F**) The ratio of GAB1-SHP2 concentrations at the cell center to the cell surface (*C_G_*_1*S*2_(0)/*C_G_*_1*S*2_(*R*)) after 5 min of 10 ng/mL EGF was computed using 2,000 draws from model parameter distributions and plotted as a raincloud plot. In panels **B** and **F**, distribution medians are indicated by solid white lines.

The robustness of the model prediction is further indicated by ensemble predictions, i.e., model simulations using many draws from fixed and posterior parameter distributions, for the ratio of the center versus surface concentrations of GAB1-SHP2. For the vast majority of ensemble simulations, the ratio of the cell center and surface GAB1-SHP2 concentrations is greater than 0.75 (**Fig. 2F**). Therefore, the model prediction of GAB1-SHP2 complexes being maintained distal from the cell surface appears robust even when accounting for uncertainty in all model parameters.

### Steady-state Model Predictions

Predictions of the full model were compared to a simplified steady-state model in which EGFR was considered only for its ability to activate SFKs, with the abundance of pEGFR chosen as that predicted by the full model for a 5-min response to 10 ng/mL EGF. This version of the model allowed an assessment of the effect of GRB2-GAB1 binding on the persistence of GAB1-SHP2 and whether the system is characterized by pseudo-steady behavior. See *Methods* for details. Solutions for active and inactive SFKs matched the solutions from the full reaction-diffusion model nearly identically (**Fig. S2A**). Predictions of the simplified model for GAB1-SHP2, pGAB1, GAB1, and SHP2 agreed with predictions of the full model to within ∼2% error or less (**Fig. S2B-E**). In addition to supporting the conclusion that the length scale for GAB1-SHP2 complex persistence is ≥ *R*, these results demonstrate that GAB1-SHP2 complexes are governed by nearly pseudo-steady behavior with respect to EGFR dynamics. These results also demonstrate that GRB2-GAB1 binding, which was neglected in the simplified model, has little influence on GAB1-SHP2 complex persistence.

### Parameter Sensitivity Analysis

To identify rate processes that strongly influence the concentration and distribution of GAB1-SHP2 complexes, the extended Fourier amplitude sensitivity test (eFAST) approach was used to conduct a multivariate parameter sensitivity analysis. Model parameters were varied by up to a factor of 1,000 from their base values (see *Methods*), and a 5-min response to 10 ng/mL EGF was considered (**Fig. 3A**). For most parameters, variance in model outputs was largely explained by interactions among parameters rather than the main effects of parameters in isolation. The average GAB1-SHP2 concentration was most sensitive to perturbations to SHP2 binding to (*k_S2,f_*) and unbinding from GAB1 (*k_S2,r_*). To determine parameters that control the distribution of GAB1-SHP2 complexes, we calculated the distance from the cell surface where the concentration of GAB1-SHP2 reaches 50% of its membrane value (*r*_1/2_). *r*_1/2_ was most sensitive to changes in the diffusivities of SFKs, GAB1, and pGAB1-SHP2. *r*_1/2_ was also sensitive, to a lesser degree, to changes in SHP2 binding kinetics, dephosphorylation kinetics of GAB1 (*k_G1dp_*), and the inactivation kinetics of SFKs (*k_S,i_*). The ratio of GAB1-SHP2 at the cell center versus cell surface was sensitive to the same perturbations. Consistent with results in **Fig. S2**, perturbations to rate constants governing EGFR-GRB2 interactions had little effect on GAB1-SHP2 complex abundance or persistence. Univariate perturbations to the most important parameters from the eFAST analysis confirmed their relative influence on the spatial distribution of GAB1-SHP2 complexes and demonstrated expected qualitative trends (**Fig. 3, B-E**). For example, a 100-fold increase in *k_G1dp_* limited the concentration of GAB1-SHP2 at the cell center to ∼60% of its value at the membrane, a 40% decrease from the base model prediction. Notably, the sensitivity of the GAB1-SHP2 length scale to changes in the kinetics of SHP2 binding were most apparent when decreasing the diffusivity of SFKs (**Fig. 3, D and E**). This indicates that SHP2 binding becomes significantly more important as the mobility of SFKs becomes limited, an effect that is explored in further detail later.

**Fig. 3.**
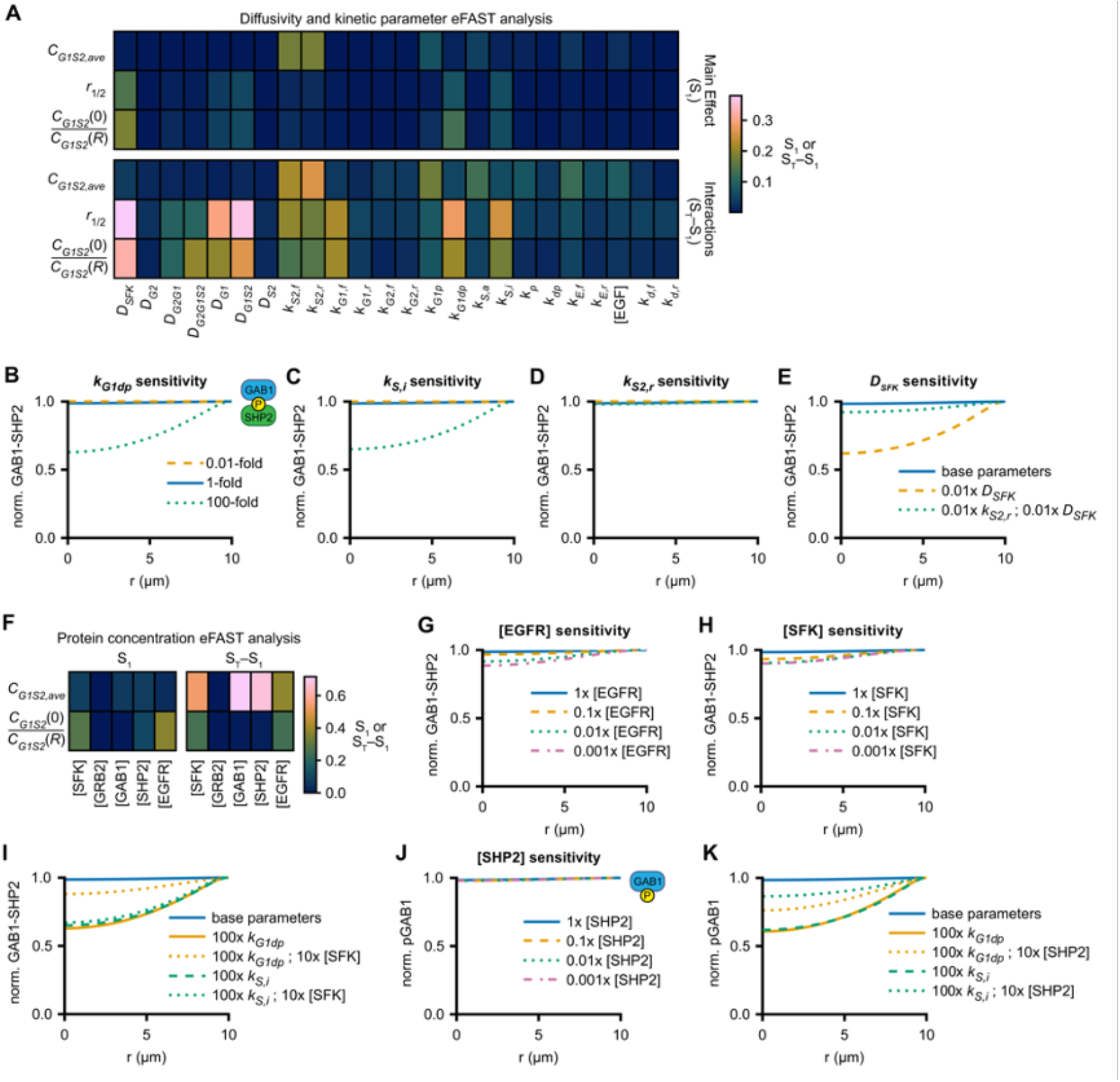
Model sensitivity analysis identifies processes controlling the length scale of GAB1-SHP2 persistence. (**A**) Extended Fourier amplitude sensitivity test (eFAST) was used to perform parameter sensitivity analysis of model predictions for the spatially averaged concentration of GAB1-SHP2 complexes (*C_G1S2,ave_*), the distance over which the GAB1-SHP2 concentration decays to 1/2 of the maximum GAB1-SHP2 concentration at the cell surface (*r_1/2_*), and the ratio of GAB1-SHP2 concentrations at the cell center to the cell surface (*C_G_*_1*S*2_(0)*/C_G_*_1*S*2_(*R*)) after a 5-min EGF treatment. Model sensitivities were calculated as first- and total-order eFAST indices (S_1_ and S_T_, respectively), followed by calculating the interaction index as S_T_−S_1_. Model parameters were permitted to vary by up to a factor of 1,000 above or below their base values (see below) over 24,000 simulations (1,000 simulations per parameter varied). (**B-E**) Model predictions for the normalized concentration of GAB1-SHP2 after a 5-min EGF treatment (10 ng/mL) were compared with predictions when *k_G1dp_*, *k_S2,r_*, *k_S,i_,* or *D_SFK_* were varied by a factor of 100 as indicated. (**F**) eFAST was used to assess the model sensitivity to changes in initial protein concentrations when allowing the abundance of each protein to vary between 100 and 1,000,000 molecules per cell over 5,000 total simulations. (**G-K**) Model predictions for the normalized concentration of GAB1-SHP2 or pGAB1 after 5-min EGF treatment were compared with predictions when the concentration of SFKs or the indicated kinetic parameters were varied by up to a factor of 1,000 as indicated. For all panels, the base values for diffusivities and kinetic parameters were used as the as described in Table 1. For eFAST calculations, 10 ng/mL was used as the baseline EGF concentration.

We next determined the model’s sensitivity to perturbations in initial protein concentrations, again using the eFAST approach. This analysis revealed that the total concentration and the center-to-surface ratio of GAB1-SHP2 were highly sensitive to changes in SFK and EGFR expression, yet only the total concentration of complexes was sensitive to changes in SHP2 expression (**Fig. 3F**). Surprisingly, however, large decreases in the abundances of EGFR and SFKs had very minor effects on the center-to-surface ratio of GAB1-SHP2 (**Fig. 3, G and H**). We investigated this further by analyzing whether SFK overexpression could compensate for increases in SFK inactivation and GAB1 dephosphorylation when determining the length scale for GAB1-SHP2 complex persistence. A ten-fold increase in SFK abundance can substantially offset the effect of an elevated GAB1 dephosphorylation rate (**Fig. 3I**), but the same increase cannot offset a faster rate of SFK inactivation. Collectively, the observations in **Fig. 3G-I** are most likely explained by the fact that SFKs persist appreciably at the cell center (**Fig. 2A**) and that the length scale for active SFKs—which directly affects the length scale of GAB1-SHP2 persistence—is independent of concentration but not the rate of SFK inactivation (Eq. 13). In the case of increased GAB1 dephosphorylation, the length scale for active SFKs remains unchanged, and SFK overexpression compensates by significantly increasing the average rate of GAB1 phosphorylation over the length scale where SFKs are active. Thus, increasing SFK expression can compensate for perturbations to the rate constants for some processes but not others. The characteristic length scales of actives SFKs and GAB1-SHP2 are described in further detail later.

Based on the protein concentration sensitivity analysis (**Fig. 3F**) and previously reported findings^12, 71^, we next sought to determine if SHP2 binding could protect GAB1 from being dephosphorylated. Consistent with the results in **Fig. 3D**, GAB1 phosphorylation is almost completely insensitive to changes in SHP2 expression alone (**Fig. 3J**). However, SHP2 overexpression significantly compensates for the effect of increasing the rate of GAB1 dephosphorylation or SFK inactivation on the persistence phospho-GAB1 (**Fig. 3K**). This occurs because unbound and dephosphorylated GAB1 near the cell center is more isolated from active SFKs than at the cell surface. This physical separation magnifies the protective effect of SHP2’s binding on phosphorylated GAB1. Based on these results, elevated SHP2 concentration can be expected to promote GAB1-SHP2 persistence by sustaining GAB1 phosphorylation through shielding it from phosphatases in the scenario when SFKs do not persist appreciably away from the cell surface or EGFR.

### Effect of SFK Activity Confinement to the Membrane

While we and other investigators have found active SFKs in the cytosolic compartment^12, 35^, some investigators report that active SFKs are confined primarily to the membrane due to palmitoylation or other post-translational modifications^72^. To further investigate the effect of SFK compartmentalization on the GAB1-SHP2 persistence length scale, we set the diffusivity of active SFKs to nearly zero, confining their activity to the membrane. This change caused the center-to-surface ratio of GAB1-SHP2 to drop significantly to ∼0.64 on average compared to ∼0.98 with the base model (**Fig. 4, A and B**). Overall, confining SFK activity to the cell membrane significantly decreased the length scale of GAB1-SHP2 persistence, with the center-to-surface ratio of GAB1-SHP2 falling well below 0.5 for a large number of model ensemble predictions. This occurs because *δ_SFK,a_*, the length scale of active SFK persistence, is comparable to *R* for the base model parameter values. However, the GAB1-SHP2 center-to-surface ratio is still greater than 0.5 for > ∼60% of model ensemble predictions, indicating the complex is still expected to persist appreciably in the case of membrane-localized SFK activity.

**Fig. 4.**
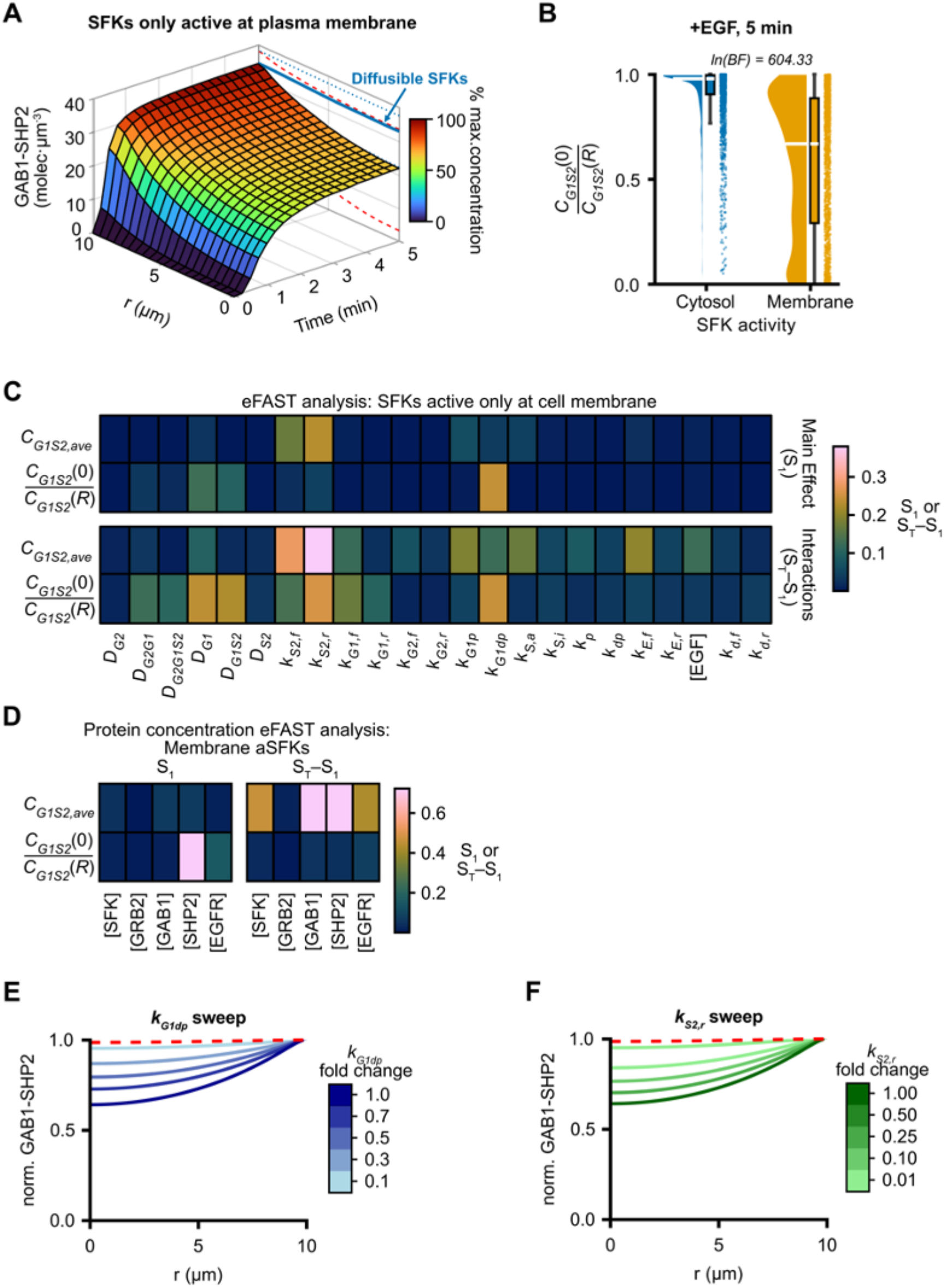
Confining SFK activity to the plasma membrane modestly reduces GAB1-SHP2 complex persistence. (**A**) Model predictions are shown for the concentrations GAB1-SHP2 complexes as a function of distance from the cell center (*r*) and time following 10 ng/mL EGF treatment when SFKs were only permitted to be active at the cell surface for 2,000 draws from model parameter distributions. The surface indicates the median ensemble prediction for SFKs only active at the cell surface, and red dashed lines indicate the 68.2% credible interval around this prediction at *t* = 5 min. The solid and dotted blue lines denote the median and 68.2% credible intervals, respectively, for the model-predicted GAB1-SHP2 concentration profile at *t* = 5 min with diffusible active SFKs for the same ensemble of parameter values. (**B**) The ratio of GAB1-SHP2 concentrations at the cell center to the cell surface (*C_G_*_1*S*2_(0)/*C_G_*_1*S*2_(*R*)) after 5 min of 10 ng/mL EGF was computed for the 2,000 parameter sets from **A** when active SFKs are allowed to diffuse in the cytosol or are confined to the cell surface. The distribution medians are indicated by solid white lines, The natural logarithm of the Bayes factor (BF) indicates strong evidence in favor of the alternative hypothesis that the difference in the distribution means is different than zero. (**C**) eFAST was used to perform parameter sensitivity analysis of *C_G1S2,ave_*, *r_1/2_*, and *C_G_*_1*S*2_(0)*/C_G_*_1*S*2_(*R*) after a 5-min EGF treatment when active SFKs were confined to the plasma membrane. Model sensitivities were calculated as first- and total-order eFAST indices (S_1_ and S_T_) where model parameters were permitted to vary by up to a factor of 1,000 above or below their base values (1,000 simulations per parameter varied). (**D**) eFAST was used to assess the model sensitivity to changes in initial protein concentrations when active SFKs were confined to the plasma membrane when allowing the abundance of each protein to vary between 100 and 1,000,000 molecules per cell over 5,000 total simulations. (**E, F**) Model predictions are shown for the normalized concentration of GAB1-SHP2 after a 5-min 10 ng/mL EGF treatment when active SFKs were confined to the plasma membrane and where *k_G1dp_* and *k_S2,r_* were varied by the indicated fold changes from their base values (see below). The concentration of GAB1-SHP2 was normalized to the maximum value within each curve. For comparison, the median model prediction for the normalized GAB1-SHP2 concentration profile at *t* = 5 min with diffusible active SFKs (solid blue line in **A**) is shown as a dashed red line in panels **E** and **F**. Base values for all parameters were used as described in Table 1. 10 ng/mL was used as the baseline EGF concentration for eFAST calculations.

We next performed another eFAST analysis to determine whether the model parameter sensitivities changed due to active SFK membrane confinement (**Fig. 4C**). The sensitivity of the overall GAB1-SHP2 concentration in this version of the model was relatively unchanged compared to the model with diffusible SFKs. However, the GAB1-SHP2 center-to-surface ratio was significantly more sensitive to the kinetics of SHP2-to-GAB1 binding and was entirely insensitive to the rate constants for SFK activation and inactivation. Furthermore, the GAB1-SHP2 center-to-surface ratio was now almost entirely sensitive to perturbations in the concentration of SHP2 alone. We assessed how much the rate constants for these processes needed to be varied to compensate for confinement of SFK activity to the cell membrane by performing model calculations over a range of values for the GAB1 dephosphorylation and GAB1-SHP2 dissociation rate constants (**Fig. 4, E and F**). These calculations indicated that a 0.1-fold change in the GAB1 dephosphorylation rate constant (*k_G1dp_*) is needed to obtain a similar GAB1-SHP2 center-to-surface ratio as for the base parameter values and compensate for active SFK membrane confinement. This fold-change falls well within the range of plausible values for *k_G1dp_* based on its posterior distribution (**Fig. S1B**). For the rate of GAB1-SHP2 dissociation (*k_S2,r_*), a 0.01-fold change from the base value was required to match the base model center-to-surface ratio of GAB1-SHP2, but we note that this change actually led to an overall increase in the cell-center concentration of cytosolic GAB1-SHP2 above that predicted by the base parameter values. This fold-change also falls within the physically plausible range of off-rates for SH2 domain binding to phospho-tyrosines^73, 74^. Thus, even if our base model assumption that active SFKs are diffusible is incorrect, these results and model ensemble simulations indicate that GAB1-SHP2 complexes can be expected to persist appreciably throughout cells even when SFK activity is confined to the cell membrane.

### Estimating Length Scales by Order of Magnitude Analysis

To validate model predictions, we developed an independent estimate for the length scale of GAB1-SHP2 complex persistence based on scaling analyses of the conservation equations for key species. The analysis is described in *Methods*, and the conceptual model describing why specific length scales were added is presented in **Fig. 5A**. Using samples from the model parameter ensemble, the order of magnitude analysis predicts that the median length scale of GAB1-SHP2 persistence is *δ_G_*_1*S*2_ ∼ 11 µm, with approximately one-half of that determined by the length scale of which SFKs are active *δ_SFK,a_* (median ∼5 µm). Of the two remaining length scales that contribute to *δ_G_*_1*S*2_, the length scale for GAB1 dephosphorylation *δ_dep_* is the larger (median ∼5 µm), and the length scale for GAB1-SHP2 dissociation *δ_dis_* is the smaller (median < 1 µm) (**Fig. 5B**). These estimates are consistent with the results shown in **Fig. 2A** and **Fig. 2D**.

**Fig. 5.**
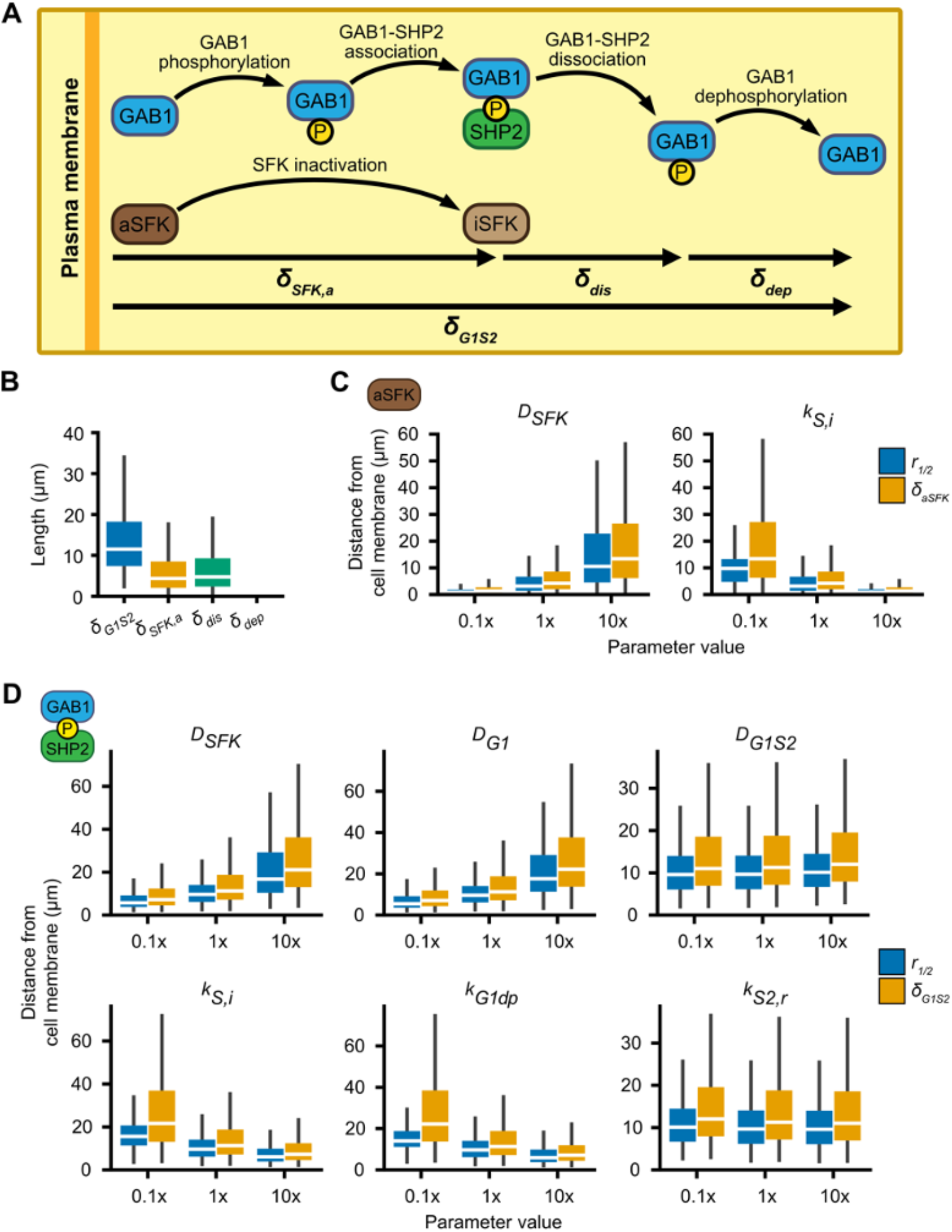
Order of magnitude length scale estimates support the conclusions of the full model. (**A**) The key reaction-diffusion processes used to estimate an order-of-magnitude length scale estimate for GAB1-SHP2 association are illustrated schematically. Over the length scale for which SFKs are active (*δ_SFK,a_*), GAB1 is phosphorylated, which enables its binding to SHP2. Beyond the length scale over which SFKs remain active distal from EGFR, GAB1-SHP2 association is extended by the length scales over which GAB1-SHP2 dissociates (*δ_dis_*) and GAB1 dephosphorylation occurs (*δ_dep_*). (**B**) Distributions for the order-of-magnitude length scale estimates *δ_G1S2_, δ_SFK,a_*, *δ_dis_*, and *δ_dep_* were obtained using 2,000 samples from model parameter distributions. (**C, D**) Model predictions for the distance over which the concentrations of active SFKs and GAB1-SHP2 fall to half their concentrations at the cell surface (*r*_1/2_) were compared to order-of-magnitude estimates for the length scales (*δ_aSFK_* and *δ_G1S2_*, respectively) over which these species persist distal from the cell surface when the indicated parameters were varied by a factor of 10 from their base values, as described in Table 1. Model predictions for *r*_1/2_ were made by setting the cell radius to 100 µm because length scale values > 10 µm could not be obtained with the base cell radius of 10 µm. *δ_aSFK_* and *δ_G1S2_* values were calculated according to Eqs. 13 and 16, respectively.

We also compared the length scale estimates for active SFKs (Eq. 13) and GAB1-SHP2 (Eq. 16) against the model-predicted length scales *r*_1/2_. Because *r*_1/2_ ≥ *R* for the base model, the cell radius was set to 100 µm for the calculations shown here. Distributions for the length scales were calculated with parameter ensemble samples using the base ensemble values and for 10-fold variations, up or down, in the parameters included in Eq. 16 (**Fig. 5C**). The length scale predictions agreed well for both active SFKs and GAB1-SHP2 across all parameter variations. The strong agreement indicates that Eqs. 13 and 16 capture the essential processes for determining the process length scales. Note that effects of cell curvature are neglected in our length scale estimates, and that curvature effects are anticipated to reduce the GAB1-SHP2 length scale, *δ_G1S2_*, by ∼10-20% based on model ensemble simulations for spherical and rectangular coordinates (**Fig. S3**). Overall, though, the length scale estimates generated concur with the predictions of the full model and support the conclusion that GAB1-SHP2 complexes persist over the entire intracellular length scale in response to EGFR activation, even given the uncertainty in model parameter values.

### Visualizing EGFR-driven GAB1-SHP2 complexes in carcinoma cells

To validate the central model prediction that GAB1-SHP2 complexes persist throughout the cell in response to EGFR activation, we used proximity ligation assays (PLA) and immunofluorescence (IF) microscopy in HeLa cells and updated model calculations for known HeLa protein abundances. The stoichiometry and expression of pathway proteins in HeLa cells diverge substantially from the assumed expression levels in our base model^33^. Notably, GAB1 is expressed at just ∼1,500 copies/cell. While the overall concentrations of model species vary greatly across the ensemble of model predictions (**Fig. 6A-C**; note the dashed red lines at *t* = 5 min), calculations based on HeLa cell protein abundances nonetheless suggest that GAB1-SHP2 complexes should be distributed throughout the cell interior. Furthermore, the median prediction for the center-to-surface ratio of GAB1-SHP2 decreases by only ∼10% compared to the base model, and the ratio is greater than or equal to 0.5 for the vast majority of HeLa simulations (**Fig. 6D**). Since active SFKs still persist appreciably in these simulations and because SHP2 is expressed in excess compared to GAB1 in HeLa cells (∼300,000 versus ∼1,500 molecules, respectively) and also protects GAB1 from dephosphorylation, it is not surprising that the length scale of GAB1-SHP2 persistence is relatively unchanged despite the low abundance of GAB1 in these cells. Thus, the reaction-diffusion mechanism described by the model distributes even relatively small numbers of GAB1-SHP2 complexes throughout the cell interior. The equivalence of predicted length scales for the base model and HeLa protein abundances is consistent with the lack of species concentration dependence in the estimate for *δ_G_*_1*S*2_ in Eq. 16.

**Fig. 6.**
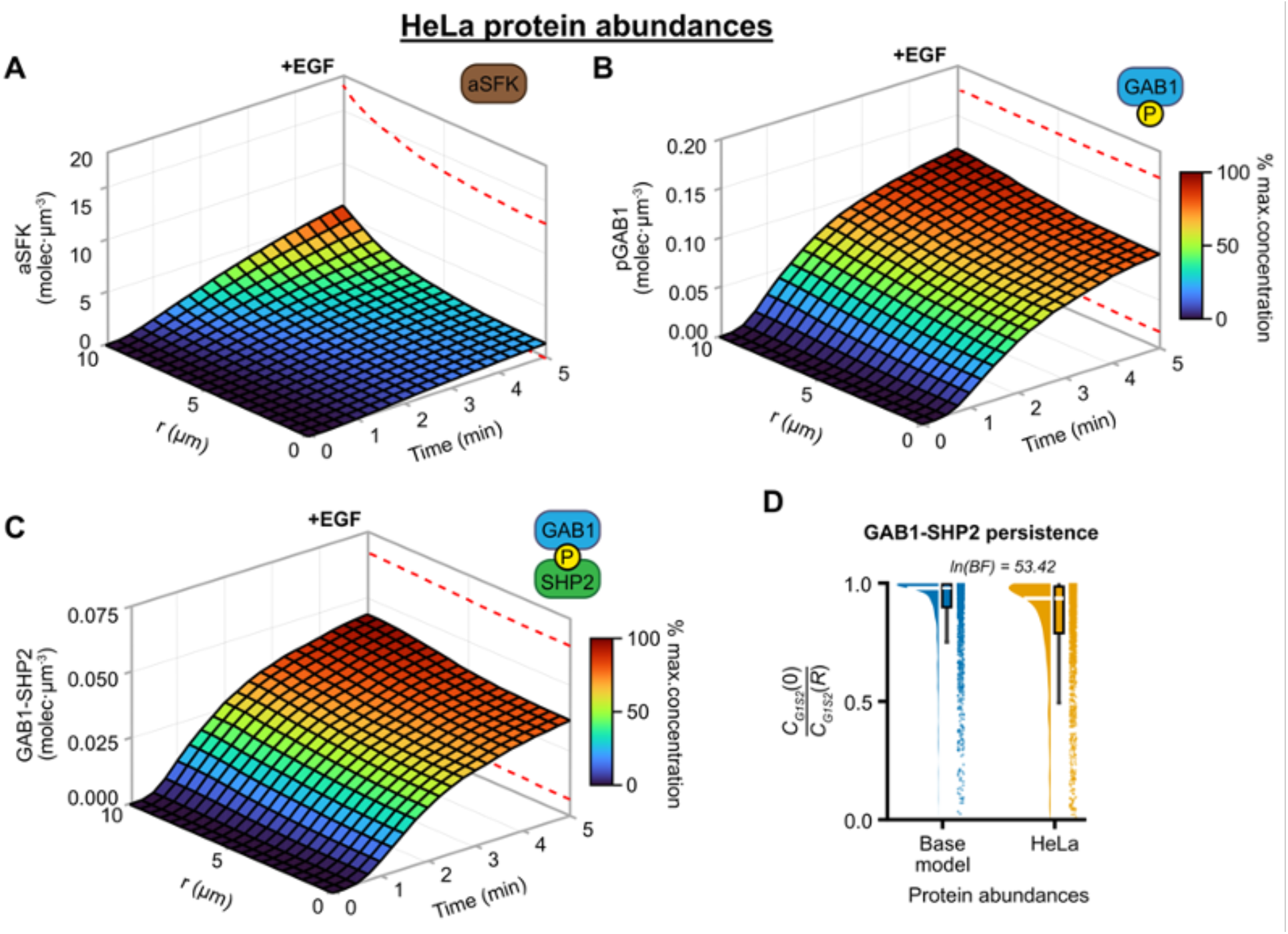
Qualitative model predictions hold for HeLa cell protein abundances, which include a stoichiometrically limiting concentration of GAB1. (**A-C**) Model predictions using HeLa protein abundances for the concentrations of active SFKs, phosphorylated GAB1, and GAB1-SHP2 complexes were plotted as a function of distance from the cell center (*r*) and time following a 5-min 10 ng/mL EGF treatment using 2,000 samples from model parameter distributions. The surface indicates the median ensemble prediction, and red dashed lines indicate the 68.2% credible interval around this prediction at *t* = 5 min. (**D**) Model predictions for *C_G_*_1*S*2_(0)/*C_G_*_1*S*2_(*R*) were calculated for a 5-min EGF treatment (10 ng/mL) when using the base model protein abundances and HeLa protein abundances for 2,000 samples from model parameter distributions. The natural logarithm of the Bayes factor (BF) indicates strong evidence in favor of the alternative hypothesis that the difference in the distribution means is different than zero

As a specific cell background for visualizing GAB1-SHP2 complexes relative to EGFR, we used gene-edited HeLa cells expressing an mVenus-HRAS fusion from one endogenous *HRAS* locus^58^, which enables membrane visualization without use of stains. We first validated the use of PLA for determining the subcellular protein localization by performing EGFR-GRB2 PLA and immunofluorescence for EGFR (**Fig. 7A**). EGFR and GRB2 strongly associate upon EGFR phosphorylation. EGFR-GRB2 PLA signals should thus be almost entirely colocalized with an EGFR IF signal generated using an independent primary antibody. As expected, exogenous EGF produced the obvious formation of EGFR-positive puncta by IF and EGFR-GRB2 PLA signals, each of which were visible throughout cells (**Fig. 7, B and C**). EGFR puncta form due to EGFR association with clathrin-coated pits and endocytosis, effects not explicitly considered in our model but described in more detail in the *Discussion*. PLA signals seen with primary antibody controls indicated that some of the apparent basal EGFR-GRB2 association in serum-starved cells may result from non-specific binding of PLA probes (**Fig. S4A**). Most importantly, colocalization analysis of EGFR-GRB2 PLA and EGFR IF signals indicated that nearly all EGFR-GRB2 puncta were colocalized with EGFR identified independently (**Fig. 7, D and E**). This result provides confidence in the ability of PLA to accurately represent the subcellular localization of protein complexes.

**Fig. 7.**
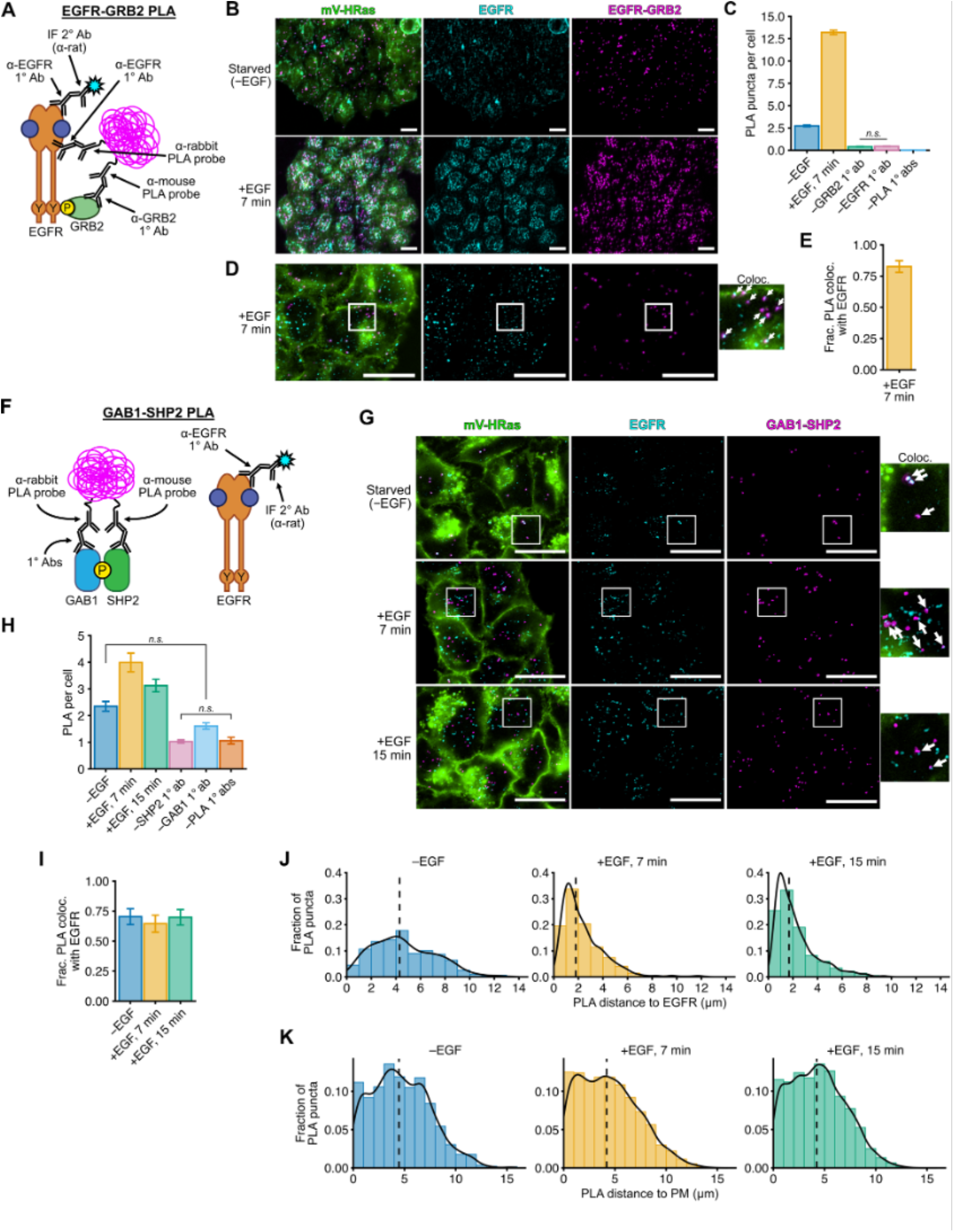
GAB1-SHP2 complexes are distributed throughout the cytoplasm and distal from EGFR in EGF-treated cancer cells. (**A**) Schematic representation of the proximity ligation assay (PLA) and experimental setup. EGFR-GRB2 complexes were stained via PLA and EGFR was stained via standard immunofluorescence (IF) using an alternative EGFR antibody to confirm the colocalization of the two signals. (**B**) Images of HeLa/mVenus-HRAS that were treated with 10 ng/mL EGF as indicated, fixed, and stained for EGFR and EGFR-GRB2 by IF and PLA, respectively. Scale bars, 20 μm. Images are representative of n = 6 to 7 independent coverslips per condition. (**C**) Quantification of PLA signals per cell for the conditions shown in **B** and PLA primary antibody controls (Fig. S2-12A) was determined for individual cells (n = 3,909 to 10,658 cells per condition). The Kruskal-Wallis test indicated a statistically significant difference among groups (χ^2^ = 14,082.64, *p* = 0.00), and the Dunn post-hoc test was used to determine *p* values for pairwise comparisons. *p* < 0.05 for all comparisons except the one indicated comparison. Error bars indicate 95% confidence intervals around the mean. (**D**) High-magnification images of HeLa/mV-HRAS cells treated as indicated and stained via IF and PLA for EGFR and EGFR-GRB2, respectively. The far-right panel (“Coloc.”) shows 2.5× magnification of the region marked by the white square. White arrows indicate colocalization between EGFR and EGFR-GRB2 puncta. Scale bars, 20 μm. Images are representative of n = 6 independent coverslips. (**E**) Fraction of EGFR-GRB2 PLA puncta co-localized with EGFR puncta in treated HeLa/mV-HRAS cells stained using the full PLA reaction for the images shown in **D** (n = 6 coverslips). Error bars indicate mean ± S.E.M. (**F**) A combination of proximity ligation assay (PLA) for GAB1 and SHP2 and standard immunofluorescence (IF) for EGFR was used to determine the localization of GAB1-SHP2 complexes relative to EGFR. The schematic shows the basic elements of the approach for using three primary and three secondary antibodies. (**G**) HeLa cells expressing mVenus-HRAS were treated with 10 ng/mL EGF for *t* ≤ 15 min, fixed, and stained for PLA and IF. Maximum-intensity *z*-projections from epifluorescence imaging are shown. The rightmost column (“colocalization”) shows 2.5× magnifications of the regions indicated by white rectangles. White arrows indicate colocalization (overlap) of EGFR and GAB1-SHP2 signals (i.e., EGFR and GAB1-SHP2 colocalization). Scale bars, 20 μm. Images are representative of n = 5 to 7 independent coverslips per condition. (**H**) The number of PLA puncta per cell for the conditions in **G** and PLA primary antibody controls (Fig. S2-12B) was determined for individual cells (n = 532 to 811 cells per condition). Histograms for the distributions of PLA signals per cell are indicated to the right of the box plots. The Kruskal-Wallis test indicated a statistically significant difference among groups (χ^2^ = 2243.55, *p* = 0.00). The Dunn post-hoc test was used to determine between-group significance. *p* < 0.05 for all pairwise comparisons except the two comparisons indicated. Error bars indicate 95% confidence intervals around the mean. (**I**) The fraction of GAB1-SHP2 puncta co-localized with EGFR puncta was computed for the conditions indicated in **G** (n = 5 to 7 coverslips per condition). The Kruskal-Wallis test indicated there was not a statistically significant difference among the groups (χ^2^ = 0.28, *p* = 0.87). Error bars indicate mean ± S.E.M. (**J**) The distances of non-EGFR-colocalized GAB1-SHP2 PLA signals to the nearest EGFR signal were computed for the conditions in **G** using cells with at least one PLA signal (n = 343 to 756 PLA puncta per condition, measured from 291 to 394 cells per condition). (**K**) The distances of GAB1-SHP2 PLA signals to the plasma membrane (PM) were computed for the conditions in **G** using cells with at least one PLA signal (n = 343 to 756 PLA puncta per condition, measured from 291 to 394 cells per condition). In **J** and **K**, the median value of each distribution is indicated with a dashed vertical line.

We next performed GAB1-SHP2 PLA and EGFR IF in parallel (**Fig. 7F**). As expected, exogeneous EGF produced clear GAB1-SHP2 PLA signals and EGFR puncta throughout the cells (**Fig. 7G-H**). Primary antibody controls indicated that the presence of low baseline PLA signal in serum-starved cells is likely due to a combination of some basal GAB1-SHP2 association and non-specific binding of the PLA anti-rabbit probe to epitopes in proximity to the SHP2 primary antibody. The observed increase in GAB1-SHP2 PLA signal with EGF is consistent with prior co-immunoprecipitation measurements^12^. Quantitative image analysis revealed that a significant fraction of GAB1-SHP2 PLA puncta are not colocalized with EGFR at all time points (**Fig. 7I**). Additionally, GAB1-SHP2 PLA signals can be seen both proximal to the cell membrane (mV-HRAS signal) and throughout the cell interior. We note that the experimentally observed fraction of EGFR-bound GAB1-SHP2 is significantly greater than that predicted by the model, but increasing the amount of complex association with EGFR to match PLA experiments does not significantly impact the length scale of GAB1-SHP2 persistence (**Fig. S5**; compare to **Fig. 7I**), which is consistent with model sensitivity analyses (**Fig. 3A** and **Fig. 4C**). Quantification of the distances of GAB1-SHP2 PLA signals to EGFR and mVenus-HRAS signals indicate that these complexes indeed persist throughout the cytoplasm over relatively large distances with respect to the cell membrane and EGFR (**Fig. 7, J and K**). Thus, the PLA results provide experimental support for the model prediction that GAB1-SHP2 complexes can exist separately from the signal-initiating EGFR and throughout the cytosol in response to EGF.

## DISCUSSION

The predictions of our reaction-diffusion model suggest that membrane-bound EGFR regulates signaling over the entire intracellular length scale through GAB1-SHP2 complexes that are maintained by the intermediary action of SFKs (**Fig. 8**). Furthermore, this prediction is robust regardless of whether SFKs are membrane-anchored or diffusible. This conceptual model stands in contrast to the more conventional view that receptor-bound complexes, such as GAB1-SHP2, access different subcellular compartments through directed membrane localization and trafficking processes. This new understanding may help explain prior apparently contradictory observations. For example, recent work demonstrates that RAS activity and ERK phosphorylation paradoxically remain dependent on EGFR activity even after EGFR and RAS become physically separated due to EGFR endocytosis and membrane retention of RAS^58, 67, 75^. Based on our work, GAB1-SHP2 complexes, generated by active endocytosed EGFR, may diffuse back to the cell membrane and thereby promote RAS activity. SHP2 promotes RAS activity via dephosphorylation of GAB1 at Y317, a p120-RASGAP (RASA1) binding site^13^, or by dephosphorylating RASGAP binding sites directly on EGFR. Other mechanisms of RAS regulation by SHP2 have been proposed^6, 76–78^, including the possibility of SHP2 acting as a scaffold that recruits GRB2-SOS complexes to the cell membrane^7, 79^. Aside from regulating RAS activity, SHP2 has also been reported to regulate the phosphorylation of paxillin^80, 81^ and the transmembrane adaptor protein PAG/CBP^76^. GAB1 also mediates signal transduction through other EGFR-downstream pathways, most notably by localizing phosphoinositide 3-kinase (PI3K) to the membrane and thereby promoting activation of the PI3K-AKT pathway^10^. Furthermore, GAB1 can mediate these interactions at the plasma membrane in an EGFR-independent manner via PH domain-mediated binding to membrane-bound PIP_3_. Thus, there are several GAB1 and SHP2 signaling targets that our analysis suggests EGFR can distally regulate via the reaction-diffusion mechanism described here.

**Fig. 8.**
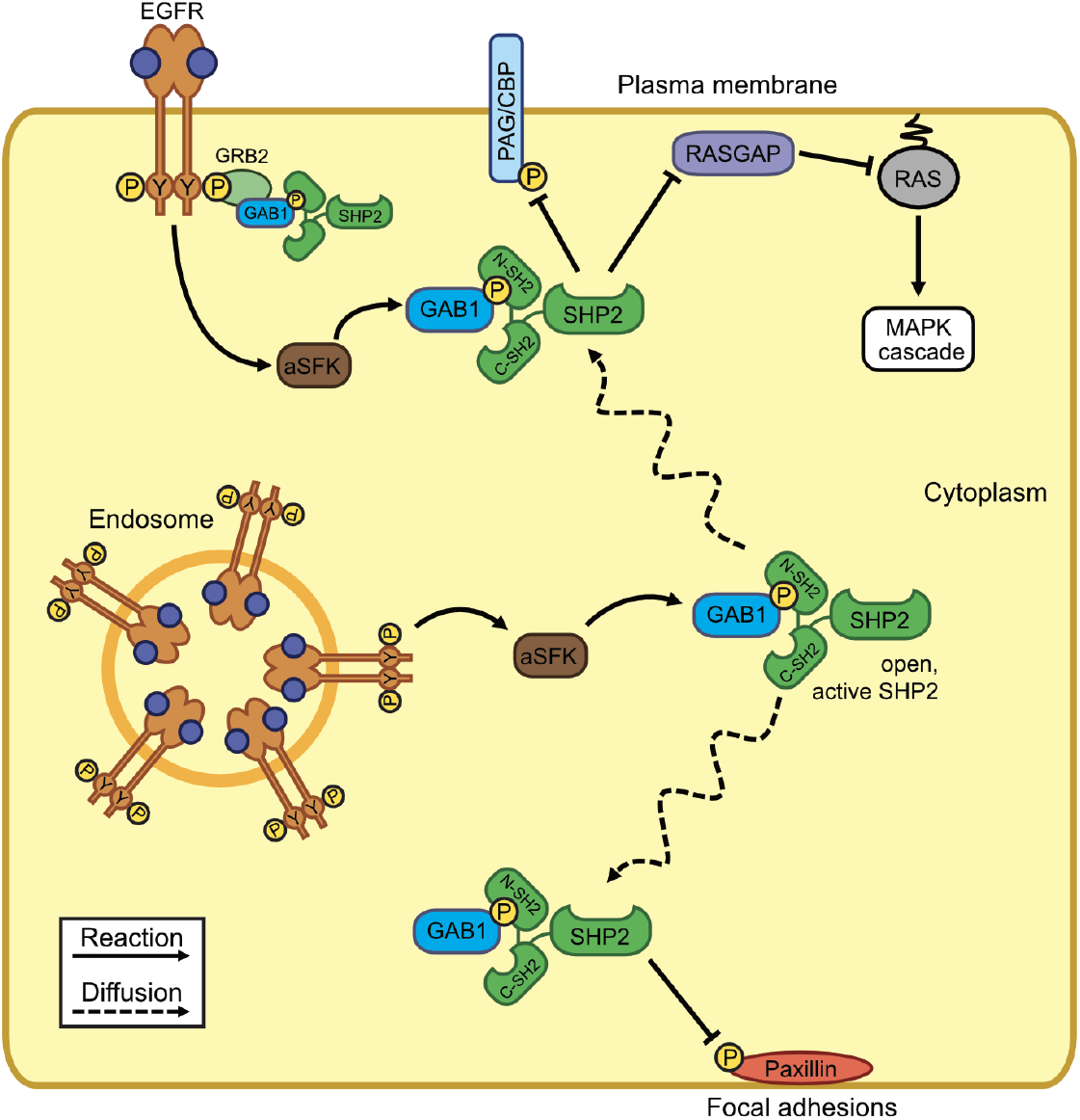
Diffusible GAB1-SHP2 complexes enable EGFR to regulate signaling processes distal from the receptor. Through the intermediate action of SFKs, EGFR promotes the formation of GAB1-SHP2 complexes, and hence active SHP2, which can then diffuse to different cellular locations and act on substrates involved in signaling pathways and cytoskeletal regulation. In cell backgrounds where the majority of EGFR is rapidly internalized after receptor activation, EGFR may still be able to influence signaling processes at the plasma membrane and in other cellular compartments via the activation of diffusible signaling complexes such as GAB1-SHP2.

Our analysis reveals that GAB1-SHP2 complex maintenance throughout the cytosol is promoted by SFK phosphorylation of GAB1 tyrosines. Interestingly, a prior study noted impaired EGF-mediated SRC Y418 phosphorylation in the non-small cell lung cancer (NSCLC) cell line H3255, which expresses a kinase-activated EGFR mutant that confers sensitivity to EGFR inhibition^82^. H3255 cells also have a functional impairment in EGFR-mediated SHP2 activity and inability to drive ERK activity as efficiently as NSCLC cells expressing wild-type EGFR^83^. Thus, EGFR may rely heavily on SFKs to activate ERK through SHP2, and disruption of this mechanism may occur with certain EGFR mutations that render cells susceptible to EGFR inhibitors.

Our analysis indicates that the GAB1-SHP2 persistence length scale is sensitive to perturbations in the rate constants for GAB1-SHP2 association and dissociation, SFK inactivation, and GAB1 dephosphorylation, but is surprisingly much less sensitive to the rate constants for SFK activation and GAB1 phosphorylation (**Fig. 3A** and **Fig. 4C**). In fact, the rate constant for SFK activation had essentially no control over the steady-state spatial distribution of GAB1-SHP2 complexes, regardless of whether active SFKs were diffusible or membrane-anchored. This is likely the result of a model topology wherein SFKs are activated only at the cell surface by membrane-bound EGFR. When allowed to diffuse, SFKs will still be rapidly inactivated throughout the cytosol even if SFK activation is made more rapid. The appearance of the rate constant for SFK inactivation but not activation in our order-of-magnitude estimate for active SFKs (Eq. 13) is consistent with this understanding. This view of the system is also consistent with our finding that EGFR only weakly controls the spatial gradient of GAB1-SHP2 complexes within the cell (**Fig. 3, A and G**).

Characteristic time scales for the reaction-diffusion mechanism modeled here suggest that GAB1 may require phosphorylation at least 5-6 times on average by SFKs throughout the cytosol to permit an individual GAB1-SHP2 complex to diffuse to the cell center from the cell surface, but the time scale of SFK inactivation suggests that GAB1 can only be phosphorylated once before SFKs become inactivated. The time scales for SHP2 binding and unbinding from GAB1, however, are much smaller than the time scale for GAB1 dephosphorylation. For a pGAB1 molecule to diffuse to the cell center, more than 100 SHP2 binding events may occur prior to GAB1 dephosphorylation. This is reflected in the strong predicted sensitivity of GAB1-SHP2 persistence and abundance to changes in SHP2 expression and GAB1-SHP2 binding kinetics, particularly for the scenario when SFKs are membrane-anchored. Thus, even with active SFKs primarily localized at or near the cell membrane, protective SHP2 binding to pGAB1 promotes the maintenance of GAB1-SHP2 complexes away from the cell surface.

The future refinement and verification of the model developed here would be greatly aided by time-resolved measurements of the distribution of GAB1-SHP2 complexes within a cell, such as could be obtained from live-cell FRET microscopy. While a GAB1-SHP2 FRET reporter pair is not presently available, our PLA experiments (**Fig. 7**) and previous findings using cell fractionation^12^ show that GAB1-SHP2 complexes exist appreciably in the cytosol and at significant distances distal from the cell membrane and EGFR. PLA experiments capture only snapshots of protein-protein interactions from fixed cells and may not fully capture all reversible protein-protein binding events occurring within cells. Model calculations with HeLa abundances predict that ∼70-250 GAB1-SHP2 molecules are formed per cell in response to EGF treatment on average, which is about an order of magnitude greater than the amount measured by PLA. Live-cell FRET experiments may more accurately capture the dynamics and spatial distribution of GAB1-SHP2 association in cells. It would also be beneficial to measure the diffusivity of all the cytosolic proteins in the model (e.g., by fluorescence recovery after photobleaching), as our current predictions rely on using empirical relationships to estimate diffusivities based on protein and complex molecular weights^42, 43^. We note, however, that the diffusivities used here are consistent with the ranges of experimentally observed values in cells^42, 84^. In addition to refining model parameters, additional cellular processes can be incorporated into the model to more completely describe the steps leading to cytosolic protein complex assembly, including receptor endocytosis^85–87^ and coordinated protein complex assembly via cytosolic scaffold proteins^88^. RTKs including EGFR and TrkA can nucleate signaling complex formation from endosomes, but endocytosis also downregulates signaling by receptors such as EGFR through degradation in lysosomes^85–87, 89^. On one hand, receptor internalization may increase GAB1-SHP2 complex persistence throughout the cell as EGFR localization shifts to the cell interior, especially given the possibility that EGFR can directly phosphorylate GAB1^53, 90^ (an interaction neglected in our model). However, receptor degradation will also terminate signaling and antagonize GAB1-SHP2 complex formation, potentially counteracting any increase in complex formation initially induced by EGFR internalization. Thus, consideration of these processes in future spatial models will be important for more completely understanding the ability of receptors to regulate signaling events at a distance via reaction-diffusion mechanisms such as the one described here.

Mechanisms similar to that described for EGFR may exist for other receptors, including other ErbB receptor family members, capable of activating diffusible non-receptor kinases that phosphorylate protein complex constituents. ErbB2/HER2, for example, activates SFKs^91^ and promotes GAB1 phosphorylation and association with SHP2^92^. SHP2 is also required for complete activation of ERK and other pathways downstream of MET and fibroblast growth factor receptors (FGFRs)^17, 93, 94^, but the spatiotemporal regulation of SHP2 activity differs between these receptor systems and compared to EGFR. In some cell contexts, MET may drive lower but more sustained and membrane-localized GAB1-SHP2 association^12, 18^. In response to FGFR activation, SHP2 binds preferentially to the membrane-bound adaptor FRS2^95^. These observations highlight the possibility that different receptors induce SHP2 activity with different dynamics and subcellular localization of GAB1-SHP2, and that not all receptors may be capable of regulating signaling over the entire cytosolic domain via GAB1-SHP2 complexes. We hypothesize that endocytosed EGFR, for example, may continue to regulate RAS signaling at the membrane via diffusible GAB1-SHP2, but such a mechanism may not exist or be necessary for other receptors, particularly if the trafficking dynamics of those receptors differ substantially from EGFR. Computational modeling is an invaluable tool in investigating such hypotheses, and our model provides a platform upon which future models could be built to study the spatiotemporal regulation of cytosolic phospho-proteins or protein complexes initiated by the activity of membrane-associated receptors and RTKs.

## AUTHOR CONTRIBUTIONS

C.M.F and M.J.L. conceptualized the project and model scope. C.M.F. and P.J.M. performed model calculations. P.J.M., W.M.D., and M.J.L. developed the order-of-magnitude scaling analysis. I.P. and A.S. developed the HeLa/mVenus-HRAS cell line. P.J.M. performed PLA experiments and associated image analysis. All authors contributed to writing of the final manuscript.

## ACKNOWLEDGEMENTS

This work was supported by the National Science Foundation 1716537 (MJL), the University of Virginia Biomedical Data Sciences Training Program NIH T32 LM012416, University of Pennsylvania Training Program in Cancer Pharmacology (R25 CA101871), and the Ashton Foundation. The Virtual Cell is supported by NIH Grant Number P41 GM103313 from the National Institute for General Medical Sciences.

## REFERENCES

1. Mayer, B.J. (2012). Perspective: Dynamics of receptor tyrosine kinase signaling complexes. FEBS Lett 586, 2575–2579. 10.1016/j.febslet.2012.05.002.

2. Morimatsu, M., Takagi, H., Ota, K.G., Iwamoto, R., Yanagida, T., and Sako, Y. (2007). Multiple-state reactions between the epidermal growth factor receotor and Grb2 as observed by using single-molecule analysis. P Natl Acad Sci USA 104, 18013–18018. 10.1073/pnas.0701330104.

3. Zhou, M.M., Harlan, J.E., Wade, W.S., Crosby, S., Ravichandran, K.S., Burakoff, S.J., and Fesik, S.W. (1995). Binding affinities of tyrosine-phosphorylated peptides to the COOH-terminal SH2 and NH2-terminal phosphotyrosine binding domains of Shc. J Biol Chem 270, 31119–31123. DOI 10.1074/jbc.270.52.31119.

4. Offterdinger, M., Georget, V., Girod, A., and Bastiaens, P.I.H. (2004). Imaging phosphorylation dynamics of the epidermal growth factor receptor. J Biol Chem 279, 36972–36981. 10.1074/jbc.M405830200.

5. Sorkin, A., McClure, M., Huang, F.T., and Carter, R. (2000). Interaction of EGF receptor and Grb2 in living cells visualized by fluorescence resonance energy transfer (FRET) microscopy. Curr Biol 10, 1395–1398. Doi 10.1016/S0960-9822(00)00785-5.

6. Ahmed, T.A., Adamopoulos, C., Karoulia, Z., Wu, X.W., Sachidanandam, R., Aaronson, S.A., and Poulikakos, P.I. (2019). SHP2 Drives Adaptive Resistance to ERK Signaling Inhibition in Molecularly Defined Subsets of ERK-Dependent Tumors. Cell Rep 26, 65-+. 10.1016/j.celrep.2018.12.013.

7. Grossmann, K.S., Rosario, M., Birchmeier, C., and Birchmeier, W. (2010). The Tyrosine Phosphatase Shp2 in Development and Cancer. Adv Cancer Res 106, 53–89. 10.1016/S0065-230x(10)06002-1.

8. Dance, M. The molecular functions of Shp2 in the Ras/Mitogen-activated protein kinase (ERK1/2) pathway. Cell Signal 20, 453–459.

9. Neel, B.G., Gu, H.H., and Pao, L. (2003). The ‘Shp’ing news: SH2 domain-containing tyrosine phosphatases in cell signaling. Trends Biochem Sci 28, 284–293. 10.1016/S0968-0004(03)00091-4.

10. Gu, H.H., and Neel, B.G. (2003). The ‘Gab’ in signal transduction. Trends Cell Biol 13, 122–130. 10.1016/S0962-8924(03)00002-3.

11. Daub, H., Wallasch, C., Lankenau, A., Herrlich, A., and Ullrich, A. (1997). Signal characteristics of G protein-transactivated EGF receptor. Embo J 16, 7032–7044. DOI 10.1093/emboj/16.23.7032.

12. Furcht, C.M., Buonato, J.M., and Lazzara, M.J. (2015). EGFR-activated Src family kinases maintain GAB1-SHP2 complexes distal from EGFR. Science Signaling 8. 10.1126/scisignal.2005697.

13. Montagner, A., Yart, A., Dance, M., Perret, B., Salles, J.P., and Raynal, P. (2005). A novel role for Gab1 and SHP2 in epidermal growth factor-induced ras activation. J Biol Chem 280, 5350–5360. 10.1074/jbc.M410012200.

14. Rodrigues, G.A., Falasca, M., Zhang, Z.T., Ong, S.H., and Schlessinger, J. (2000). A novel positive feedback loop mediated by the docking protein Gab1 and phosphatidylinositol 3-kinase in epidermal growth factor receptor signaling. Mol Cell Biol 20, 1448–1459. Doi 10.1128/Mcb.20.4.1448-1459.2000.

15. Incoronato, M., D’Alessio, A., Paladino, S., Zurzolo, C., Carlomagno, M.S., Cerchia, L., and de Franciscis, V. (2004). The Shp-1 and Shp-2, tyrosine phosphatases, are recruited on cell membrane in two distinct molecular complexes including Ret oncogenes. Cell Signal 16, 847–856. 10.1016/j.cellsig.2004.01.002.

16. Zhou, X.D., and Agazie, Y.M. (2009). Molecular Mechanism for SHP2 in Promoting HER2-induced Signaling and Transformation. J Biol Chem 284, 12226–12234. 10.1074/jbc.M900020200.

17. Schaeper, U., Gehring, N.H., Fuchs, K.P., Sachs, M., Kempkes, B., and Birchmeier, W. (2000). Coupling of Gab1 to c-Met, Grb2, and Shp2 mediates biological responses. J Cell Biol 149, 1419-1432. 10.1083/jcb.149.7.1419.

18. Maroun, C.R., Naujokas, M.A., and Park, M. (2003). Membrane targeting of Grb2-associated binder-1 (Gab1) scaffolding protein through src myristoylation sequence substitutes for Gab1 pleckstrin homology domain and switches an epidermal growth factor response to an invasive morphogenic program. Mol Biol Cell 14, 1691–1708. 10.1091/mbc.E02-06-0352.

19. Abram, C.L., and Courtneidge, S.A. (2000). Src family tyrosine kinases and growth factor signaling. Exp Cell Res 254, 1–13. DOI 10.1006/excr.1999.4732.

20. Lu, K.V., Zhu, S.J., Cvrljevic, A., Huang, T.T., Sarkaria, S., Ahkavan, D., Dang, J., Dinca, E.B., Plaisier, S.B., Oderberg, I., et al. (2009). Fyn and Src Are Effectors of Oncogenic Epidermal Growth Factor Receptor Signaling in Glioblastoma Patients. Cancer Res 69, 6889–6898. 10.1158/0008-5472.Can-09-0347.

21. Osherov, N., and Levitzki, A. (1994). Epidermal-Growth-Factor-Dependent Activation of the Src-Family Kinases. Eur J Biochem 225, 1047–1053. DOI 10.1111/j.1432-1033.1994.1047b.x.

22. Bromann, P.A., Korkaya, H., and Courtneidge, S.A. (2004). The interplay between Src family kinases and receptor tyrosine kinases. Oncogene 23, 7957–7968. 10.1038/sj.onc.1208079.

23. Maa, M.C., Leu, T.H., McCarley, D.J., Schatzman, R.C., and Parsons, S.J. (1995). Potentiation of epidermal growth factor receptor-mediated oncogenesis by c-Src: implications for the etiology of multiple human cancers. Proceedings of the National Academy of Sciences 92, 6981–6985. 10.1073/pnas.92.15.6981.

24. Okada, M., and Nakagawa, H. (1989). A Protein Tyrosine Kinase Involved in Regulation of Pp60c-Src Function. J Biol Chem 264, 20886–20893.

25. Schiesser, W.E. (1991). The numerical method of lines : integration of partial differential equations (Academic Press).

26. Ma, Y., Gowda, S., Anantharaman, R., Laughman, C., Shah, V., and Rackauckas, C. (2021). ModelingToolkit: A Composable Graph Transformation System For Equation-Based Modeling. arXiv. 10.48550/arxiv.2103.05244.

27. Rackauckas, C. (2017). DifferentialEquations.jl – A Performant and Feature-Rich Ecosystem for Solving Differential Equations in Julia. Journal of open research software 5.

28. Bezanson, J., Edelman, A., Karpinski, S., and Shah, V.B. (2017). Julia: A Fresh Approach to Numerical Computing. SIAM Review 59, 65–98. 10.1137/141000671.

29. Bieniasz, L.K. (1993). The von Neumann stability of finite-difference algorithms for the electrochemical kinetic simulation of diffusion coupled with homogeneous reactions. Journal of Electroanalytical Chemistry 345, 13–25.

30. Resasco, D.C., Gao, F., Morgan, F., Novak, I.L., Schaff, J.C., and Slepchenko, B.M. (2012). Virtual Cell: computational tools for modeling in cell biology. Wires Syst Biol Med 4, 129–140. 10.1002/wsbm.165.

31. Blinov, M.L., Schaff, J.C., Vasilescu, D., Moraru, I.I., Bloom, J.E., and Loew, L.M. (2017). Compartmental and Spatial Rule-Based Modeling with Virtual Cell. Biophys J 113, 1365–1372. 10.1016/j.bpj.2017.08.022.

32. Kholodenko, B.N. (2002). MAP kinase cascade signaling and endocytic trafficking: a marriage of convenience? Trends Cell Biol 12, 173–177. 10.1016/s0962-8924(02)02251-1.

33. Kulak, N.A., Pichler, G., Paron, I., Nagaraj, N., and Mann, M. (2014). Minimal, encapsulated proteomic-sample processing applied to copy-number estimation in eukaryotic cells. Nat Methods 11, 319–U300. 10.1038/Nmeth.2834.

34. French, A.R., Tadaki, D.K., Niyogi, S.K., and Lauffenburger, D.A. (1995). Intracellular Trafficking of Epidermal Growth-Factor Family Ligands Is Directly Influenced by the Ph Sensitivity of the Receptor-Ligand Interaction. J Biol Chem 270, 4334–4340. DOI 10.1074/jbc.270.9.4334.

35. Shan, Y.B., Eastwood, M.P., Zhang, X.W., Kim, E.T., Arkhipov, A., Dror, R.O., Jumper, J., Kuriyan, J., and Shaw, D.E. (2012). Oncogenic Mutations Counteract Intrinsic Disorder in the EGFR Kinase and Promote Receptor Dimerization. Cell 149, 860–870. 10.1016/j.cell.2012.02.063.

36. Hajdu, T., Váradi, T., Rebenku, I., Kovács, T., Szöllösi, J., and Nagy, P. (2020). Comprehensive Model for Epidermal Growth Factor Receptor Ligand Binding Involving Conformational States of the Extracellular and the Kinase Domains. Front Cell Dev Biol 8, 776. 10.3389/fcell.2020.00776.

37. Young, M.W. (2019). Dephosphorylation of Epidermal Growth Factor Receptor by Protein Tyrosine Phosphatase 1B. The FASEB journal 33.

38. Borisov, N., Aksamitiene, E., Kiyatkin, A., Legewie, S., Berkhout, J., Maiwald, T., Kaimachnikov, N.P., Timmer, J., Hoek, J.B., and Kholodenko, B.N. (2009). Systems-level interactions between insulin–EGF networks amplify mitogenic signaling. Molecular Systems Biology 5, 256. 10.1038/msb.2009.19.

39. Biscardi, J.S., Maa, M.C., Tice, D.A., Cox, M.E., Leu, T.H., and Parsons, S.J. (1999). c-Src-mediated phosphorylation of the epidermal growth factor receptor on Tyr845 and Tyr1101 is associated with modulation of receptor function. J Biol Chem 274, 8335–8343. 10.1074/jbc.274.12.8335.

40. Roskoski, R., Jr. (2015). Src protein-tyrosine kinase structure, mechanism, and small molecule inhibitors. Pharmacol Res 94, 9–25. 10.1016/j.phrs.2015.01.003.

41. Solomaha, E., Szeto, F.L., Yousef, M.A., and Palfrey, H.C. (2005). Kinetics of Src Homology 3 Domain Association with the Proline-rich Domain of Dynamins. J Biol Chem 280, 23147–23156. 10.1074/jbc.m501745200.

42. Pepperkok, R., Bre, M.H., Davoust, J., and Kreis, T.E. (1990). Microtubules are stabilized in confluent epithelial cells but not in fibroblasts. J Cell Biol 111, 3003–3012. 10.1083/jcb.111.6.3003.

43. Erickson, H.P. (2009). Size and Shape of Protein Molecules at the Nanometer Level Determined by Sedimentation, Gel Filtration, and Electron Microscopy. Biol Proced Online 11, 32–51. 10.1007/s12575-009-9008-x.

44. Tsigkinopoulou, A., Hawari, A., Uttley, M., and Breitling, R. (2018). Defining informative priors for ensemble modeling in systems biology. Nature Protocols 13, 2643–2663. 10.1038/s41596-018-0056-z.

45. Arnoud, A., Guvenen, F., Kleineberg, T., and National Bureau of Economic, R. (2019). Benchmarking global optimizers (National Bureau of Economic Research).

46. Nocedal, J. (1980). Updating Quasi-Newton Matrices With Limited Storage. Mathematics of Computation 35, 773–782.

47. Liu, D.C., and Nocedal, J. (1989). On the limited memory BFGS method for large scale optimization. Mathematical Programming 45, 503–528. 10.1007/BF01589116.

48. Revels, J., Lubin, M., and Papamarkou, T. (2016). Forward-Mode Automatic Differentiation in Julia. arXiv:1607.07892 [cs.MS].

49. Ge, H., Xu, K., and Ghahramani, Z. (2018). Turing: A Language for Flexible Probabilistic Inference.

50. Hoffman, M.D., and Gelman, A. (2011). The No-U-Turn Sampler: Adaptively Setting Path Lengths in Hamiltonian Monte Carlo.

51. Brooks, S.P., and Gelman, A. (1998). General Methods for Monitoring Convergence of Iterative Simulations. Journal of Computational and Graphical Statistics 7, 434–455. 10.1080/10618600.1998.10474787.

52. Gelman, A., and Rubin, D.B. (1992). Inference from Iterative Simulation Using Multiple Sequences. Statistical Science 7, 457–472. 10.1214/ss/1177011136.

53. Fan, Y.X., Wong, L., Deb, T.B., and Johnson, G.R. (2004). Ligand regulates epidermal growth factor receptor kinase specificity - Activation increases preference for GAB1 and SHC versus autophosphorylation sites. J Biol Chem 279, 38143–38150. 10.1074/jbc.M405760200.

54. Marino, S., Hogue, I.B., Ray, C.J., and Kirschner, D.E. (2008). A methodology for performing global uncertainty and sensitivity analysis in systems biology. J Theor Biol 254, 178–196. 10.1016/j.jtbi.2008.04.011.

55. Saltelli, A., Tarantola, S., and Chan, K.P.S. (1999). A quantitative model-independent method for global sensitivity analysis of model output. Technometrics 41, 39–56. Doi 10.2307/1270993.

56. Dixit, V.K., and Rackauckas, C. (2022). GlobalSensitivity.jl: Performant and Parallel Global Sensitivity Analysis with Julia. Journal of Open Source Software 7, 4561. 10.21105/joss.04561.

57. Deen, W.M. (1998). Analysis of Transport Phenomena, 2 Edition (Oxford University Press New York).

58. Pinilla-Macua, I., Watkins, S.C., and Sorkin, A. (2016). Endocytosis separates EGF receptors from endogenous fluorescently labeled HRas and diminishes receptor signaling to MAP kinases in endosomes. Proc Natl Acad Sci U S A 113, 2122–2127. 10.1073/pnas.1520301113.

59. Stirling, D.R., Swain-Bowden, M.J., Lucas, A.M., Carpenter, A.E., Cimini, B.A., and Goodman, A. (2021). CellProfiler 4: improvements in speed, utility and usability. BMC Bioinformatics 22. 10.1186/s12859-021-04344-9.

60. Horzum, U., Ozdil, B., and Pesen-Okvur, D. (2014). Step-by-step quantitative analysis of focal adhesions. MethodsX 1, 56–59. 10.1016/j.mex.2014.06.004.

61. Moser, B., Hochreiter, B., Herbst, R., and Schmid, J.A. (2017). Fluorescence colocalization microscopy analysis can be improved by combining object-recognition with pixel-intensity-correlation. Biotechnol J 12. 10.1002/biot.201600332.

62. Patil, I. (2021). Visualizations with statistical details: The ‘ggstatsplot’ approach. Journal of Open Source Software 6, 3167.

63. Morey, R., Rouder, J., Love, J., and Marwick, B. (2015). BayesFactor: 0.9.12-2 CRAN. Zenodo.

64. Ben-Shachar, M., Lüdecke, D., and Makowski, D. (2020). effectsize: Estimation of Effect Size Indices and Standardized Parameters. Journal of Open Source Software 5, 2815. 10.21105/joss.02815.

65. Wickham, H. (2016). ggplot2: Elegant Graphics for Data Analysis.

66. Danisch, S., and Krumbiegel, J. (2021). Makie.jl: Flexible high-performance data visualization for Julia. Journal of Open Source Software 6, 3349. 10.21105/joss.03349.

67. Surve, S., Watkins, S.C., and Sorkin, A. (2021). EGFR-RAS-MAPK signaling is confined to the plasma membrane and associated endorecycling protrusions. J Cell Biol 220. 10.1083/jcb.202107103.

68. Reddy, R.J., Gajadhar, A.S., Swenson, E.J., Rothenberg, D.A., Curran, T.G., and White, F.M. (2016). Early signaling dynamics of the epidermal growth factor receptor. P Natl Acad Sci USA 113, 3114–3119. 10.1073/pnas.1521288113.

69. Roepstorff, K., Grandal, M.V., Henriksen, L., Knudsen, S.L.J., Lerdrup, M., Grovdal, L., Willumsen, B.M., and van Deurs, B. (2009). Differential Effects of EGFR Ligands on Endocytic Sorting of the Receptor. Traffic 10, 1115–1127. 10.1111/j.1600-0854.2009.00943.x.

70. Huang, F.T., Kirkpatrick, D., Jiang, X.J., Gygi, S., and Sorkin, A. (2006). Differential regulation of EGF receptor internalization and degradation by multiubiquitination within the kinase domain. Mol Cell 21, 737–748. 10.1016/j.molcel.2006.02.018.

71. Rotin, D., Margolis, B., Mohammadi, M., Daly, R.J., Daum, G., Li, N., Fischer, E.H., Burgess, W.H., Ullrich, A., and Schlessinger, J. (1992). Sh2 Domains Prevent Tyrosine Dephosphorylation of the Egf Receptor - Identification of Tyr992 as the High-Affinity Binding-Site for Sh2 Domains of Phospholipase C-Gamma. Embo J 11, 559–567. DOI 10.1002/j.1460-2075.1992.tb05087.x.

72. Sandilands, E., Brunton, V.G., and Frame, M.C. (2007). The membrane targeting and spatial activation of Src, Yes and Fyn is influenced by palmitoylation and distinct RhoB/RhoD endosome requirements. J Cell Sci 120, 2555–2564. 10.1242/jcs.003657.

73. Ladbury, J.E., Lemmon, M.A., Zhou, M., Green, J., Botfield, M.C., and Schlessinger, J. (1995). Measurement of the binding of tyrosyl phosphopeptides to SH2 domains: a reappraisal. Proc Natl Acad Sci U S A 92, 3199–3203. 10.1073/pnas.92.8.3199.

74. Ottinger, E.A., Botfield, M.C., and Shoelson, S.E. (1998). Tandem SH2 domains confer high specificity in tyrosine kinase signaling. J Biol Chem 273, 729–735. 10.1074/jbc.273.2.729.

75. Surve, S.V., Myers, P.J., Clayton, S.A., Watkins, S.C., Lazzara, M.J., and Sorkin, A. (2019). Localization dynamics of endogenous fluorescently labeled RAF1 in EGF-stimulated cells. Mol Biol Cell 30, 506–523. 10.1091/mbc.E18-08-0512.

76. Zhang, S.Q., Yang, W.T., Kontaridis, M.I., Bivona, T.G., Wen, G.Y., Araki, T., Luo, J.C., Thompson, J.A., Schraven, B.L., Philips, M.R., and Neel, B.G. (2004). Shp2 regulates Src family kinase activity and Ras/Erk activation by controlling Csk recruitment. Mol Cell 13, 341–355. Doi 10.1016/S1097-2765(04)00050-4.

77. Nichols, R.J., Haderk, F., Stahlhut, C., Schulze, C.J., Hemmati, G., Wildes, D., Tzitzilonis, C., Mordec, K., Marquez, A., Romero, J., et al. (2018). RAS nucleotide cycling underlies the SHP2 phosphatase dependence of mutant BRAF-, NF1-and RAS-driven cancers. Nat Cell Biol 20, 1064-+. 10.1038/s41556-018-0169-1.

78. Batth, T.S., Papetti, M., Pfeiffer, A., Tollenaere, M.A.X., Francavilla, C., and Olsen, J.V. (2018). Large-Scale Phosphoproteomics Reveals Shp-2 Phosphatase-Dependent Regulators of Pdgf Receptor Signaling. Cell Rep 22, 2784–2796. 10.1016/j.celrep.2018.02.038.

79. Dance, M., Montagner, A., Salles, J.P., Yart, A., and Raynal, P. (2008). The molecular functions of Shp2 in the Ras/Mitogen-activated protein kinase (ERK1/2) pathway. Cell Signal 20, 453–459. 10.1016/j.cellsig.2007.10.002.

80. Ren, Y., Chen, Z., Chen, L., Fang, B., Win-Piazza, H., Haura, E., Koomen, J.M., and Wu, J. (2010). Critical role of Shp2 in tumor growth involving regulation of c-Myc. Genes Cancer 1, 994–1007. 10.1177/1947601910395582.

81. Ren, Y., Meng, S., Mei, L., Zhao, Z.J., Jove, R., and Wu, J. (2004). Roles of Gab1 and SHP2 in paxillin tyrosine dephosphorylation and Src activation in response to epidermal growth factor. J Biol Chem 279, 8497–8505. 10.1074/jbc.M312575200.

82. Lazzara, M.J., Lane, K., Chan, R., Jasper, P.J., Yaffe, M.B., Sorger, P.K., Jacks, T., Neel, B.G., and Lauffenburger, D.A. (2010). Impaired SHP2-Mediated Extracellular Signal-Regulated Kinase Activation Contributes to Gefitinib Sensitivity of Lung Cancer Cells with Epidermal Growth Factor Receptor-Activating Mutations. Cancer Res 70, 3843–3850. 10.1158/0008-5472.Can-09-3421.

83. Furcht, C.M., Rojas, A.R.M., Nihalani, D., and Lazzara, M.J. (2013). Diminished functional role and altered localization of SHP2 in non-small cell lung cancer cells with EGFR-activating mutations. Oncogene 32, 2346–2355. 10.1038/onc.2012.240.

84. Kumar, M., Mommer, M.S., and Sourjik, V. (2010). Mobility of Cytoplasmic, Membrane, and DNA-Binding Proteins in Escherichia coli. Biophys J 98, 552–559. 10.1016/j.bpj.2009.11.002.

85. Wang, Y., Pennock, S., Chen, X.M., and Wang, Z.X. (2002). Endosomal signaling of epidermal growth factor receptor stimulates signal transduction pathways leading to cell survival. Mol Cell Biol 22, 7279–7290. 10.1128/Mcb.22.20.7279-7290.2002.

86. Miaczynska, M., Pelkmans, L., and Zerial, M. (2004). Not just a sink: endosomes in control of signal transduction. Curr Opin Cell Biol 16, 400–406. 10.1016/j.ceb.2004.06.005.

87. Ye, H.H., Kuruvilla, R., Zweifel, L.S., and Ginty, D.D. (2003). Evidence in support of signaling endosome-based retrograde survival of sympathetic neurons. Neuron 39, 57–68. Doi 10.1016/S0896-6273(03)00266-6.

88. Good, M.C., Zalatan, J.G., and Lim, W.A. (2011). Scaffold Proteins: Hubs for Controlling the Flow of Cellular Information. Science 332, 680–686. 10.1126/science.1198701.

89. Fortian, A., and Sorkin, A. (2014). Live-cell fluorescence imaging reveals high stoichiometry of Grb2 binding to the EGF receptor sustained during endocytosis. J Cell Sci 127, 432–444. 10.1242/jcs.137786.

90. Lehr, S., Kotzka, J., Herkner, A., Klein, E., Siethoff, C., Knebel, B., Noelle, V., Brüning, J.C., Klein, H.W., Meyer, H.E., et al. (1999). Identification of Tyrosine Phosphorylation Sites in Human Gab-1 Protein by EGF Receptor Kinase in Vitro. Biochemistry-Us 38, 151–159. 10.1021/bi9818265.

91. Moasser, M.M. (2007). The oncogene HER2: its signaling and transforming functions and its role in human cancer pathogenesis. Oncogene 26, 6469–6487. 10.1038/sj.onc.1210477.

92. Fan, Y.-X., Wong, L., Marino, M.P., Ou, W., Shen, Y., Wu, W.J., Wong, K.-K., Reiser, J., and Johnson, G.R. (2013). Acquired Substrate Preference for GAB1 Protein Bestows Transforming Activity to ERBB2 Kinase Lung Cancer Mutants. J Biol Chem 288, 16895–16904. 10.1074/jbc.m112.434217.

93. Matalkah, F., Martin, E., Zhao, H., and Agazie, Y.M. (2016). SHP2 acts both upstream and downstream of multiple receptor tyrosine kinases to promote basal-like and triple-negative breast cancer. Breast Cancer Research 18. 10.1186/s13058-015-0659-z.

94. Lu, H., Liu, C., Velazquez, R., Wang, H., Dunkl, L.M., Kazic-Legueux, M., Haberkorn, A., Billy, E., Manchado, E., Brachmann, S.M., et al. (2019). SHP2 Inhibition Overcomes RTK-Mediated Pathway Reactivation in KRAS-Mutant Tumors Treated with MEK Inhibitors. Molecular Cancer Therapeutics 18, 1323–1334. 10.1158/1535-7163.mct-18-0852.

95. Kontaridis, M.I., Liu, X., Zhang, L., and Bennett, A.M. (2002). Role of SHP-2 in Fibroblast Growth Factor Receptor-Mediated Suppression of Myogenesis in C2C12 Myoblasts. Mol Cell Biol 22, 3875–3891. 10.1128/mcb.22.11.3875-3891.2002.

