## Supplement for "A spatially resolved EGFR signaling model predicts the length scale of GAB1-SHP2 complex persistence"

### Appendix S1: Equations for the full reaction-diffusion model and finite difference scheme.

#### Reaction-diffusion equations and initial and boundary conditions

The conservation equations and initial and boundary conditions for all protein monomers and complexes included in the model are described according to the following abbreviations:

Concentrations of cytosolic protein monomers and complexes:
$C_{SFK,i}$ = inactive SFK
$C_{SFK,a}$ = active SFK
$C_{G1}$ = GAB1
$C_{PG1}$ = pGAB1
$C_{G2}$ = GRB2
$C_{G2G1}$ = GRB2-GAB1
$C_{G2PG1}$ = GRB2-pGAB1
$C_{S2}$ = SHP2
$C_{PG1S}$ = pGAB1-SHP2
$C_{G2PG1S}$ = GRB2-pGAB1-SHP2

Concentrations of membrane protein monomers and complexes:
$C_{mE}$ = EGFR monomer
$C_{mES}$ = EGF-bound EGFR monomer
$C_{mESmES}$ = EGF-bound EGFR dimer
$C_{E}$ = EGF-bound pEGFR dimer
$C_{EG2}$ = EGF-bound, GRB2-bound pEGFR dimer
$C_{EG2G1}$ = EGF-bound, GRB2-GAB1-bound pEGFR dimer
$C_{EG2PG1}$ = EGF-bound, GRB2-pGAB1-bound pEGFR dimer
$C_{EG2PG1S}$ = EGF-bound, GRB2-pGAB1-SHP2-bound pEGFR dimer
$C_{pE,tot}$ = Total number of phosphorylated EGFR monomers

Model parameters:
$D_{SFK}$ = SFK diffusivity
$D_{G2}$ = GRB2 diffusivity
$D_{G2G1}$ = GRB2-GAB1 diffusivity
$D_{G2G1S2}$ = GRB2-GAB1-SHP2 diffusivity
$D_{G1}$ = GAB1 diffusivity
$D_{G1S2}$ = GAB1-SHP2 diffusivity
$D_{S2}$ = SHP2 diffusivity

$k_{S,i}$ = SFK inactivation
$k_{S,a}$ = SFK activation
$k_{G1,f}$ = GAB1 binding, forward
$k_{G1,r}$ = GAB1 binding, reverse
$k_{G1p}$ = GAB1 phosphorylation
$k_{G1dp}$ = GAB1 dephosphorylation
$k_{G2,f}$ = GRB2 binding, forward
$k_{G2,r}$ = GRB2 binding, reverse
$k_{S2,f}$ = SHP2 binding, forward
$k_{S2,r}$ = SHP2 binding, reverse
$k_{E,f}$ = EGF binding, forward
$k_{E,r}$ = EGF binding, reverse
$k_{dE,f}$ = EGFR dimerization, forward
$k_{dE,r}$ = EGFR dimerization, reverse
$k_{catE}$ = EGFR phosphorylation
$k_{dp}$ = EGFR dephosphorylation
$[EGF]$ = EGF concentration

Equations and boundary conditions: All model equations were written and discretized in spherical coordinates assuming rotational symmetry about the center of a spherical cell. Thus, only the radial component is considered for model equations. $r$ = 0 corresponds to the cell center, and $r$ = $R$ corresponds to the cell surface. $t$ = 0 corresponds to the initial time prior to EGF treatment.

$C_{SFK,i}$ conservation equation, boundary conditions, and initial condition:

$$\frac{\partial C_{SFK,i}}{\partial t} = D_{SFK}\frac{1}{r^2}\frac{\partial}{\partial r}\left(r^2\frac{\partial C_{SFK,i}}{\partial r}\right) + k_{S,i}C_{SFK,a} \quad \text{(S1.1)}$$

$$D_{SFK}\frac{\partial C_{SFK,i}}{\partial r}(R,t) = -k_{S,a}C_{SFK,i}C_{pE,tot} \quad \text{(S1.2)}$$

$$D_{SFK}\frac{\partial C_{SFK,i}}{\partial r}(0,t) = 0 \quad \text{(S1.3)}$$

$$C_{SFK,i}(r,0) = C_{o,SFK} \quad \text{(S1.4)}$$

Conservation equation and initial and boundary conditions for $C_{SFK,a}$:

$$\frac{\partial C_{SFK,a}}{\partial t} = D_{SFK}\frac{1}{r^2}\frac{\partial}{\partial r}\left(r^2\frac{\partial C_{SFK,a}}{\partial r}\right) - k_{S,i}C_{SFK,a} \quad \text{(S1.5)}$$

$$D_{SFK}\frac{\partial C_{SFK,a}}{\partial r}(R,t) = k_{S,a}C_{SFK,i}C_{pE,tot} \quad \text{(S1.6)}$$

$$D_{SFK}\frac{\partial C_{SFK,a}}{\partial r}(0,t)=0 \tag{S1.7}$$

$$C_{SFK,a}(r,0)=0 \tag{S1.8}$$

Conservation equation and initial and boundary conditions for $C_{G1}$:

$$\frac{\partial C_{G1}}{\partial t}=D_{G1}\frac{1}{r^2}\frac{\partial}{\partial r}\left(r^2\frac{\partial C_{G1}}{\partial r}\right)-k_{G1,f}C_{G1}C_{G2}+k_{G1,r}C_{G2G1}-k_{G1p}C_{SFK,a}C_{G1}+k_{G1dp}C_{PG1} \tag{S1.9}$$

$$D_{G1}\frac{\partial C_{G1}}{\partial r}(R,t)=k_{G1,r}C_{EG2G1}-k_{G1,f}C_{G1}C_{EG2} \tag{S1.10}$$

$$D_{G1}\frac{\partial C_{G1}}{\partial r}(0,t)=0 \tag{S1.11}$$

$$C_{G1}(r,0)=C_{o,GAB1} \tag{S1.12}$$

Conservation equation and initial and boundary conditions for $C_{PG1}$:

$$\frac{\partial C_{PG1}}{\partial t}=D_{G1}\frac{1}{r^2}\frac{\partial}{\partial r}\left(r^2\frac{\partial C_{PG1}}{\partial r}\right)-k_{G1,f}C_{PG1}C_{G2}+k_{G1,r}C_{G2PG1}+k_{G1p}C_{SFK,a}C_{G1}-$$
$$k_{G1dp}C_{PG1}-k_{S2,f}C_{PG1}C_{S2}+k_{S2,r}C_{PG1S} \tag{S1.13}$$

$$D_{G1}\frac{\partial C_{PG1}}{\partial r}(R,t)=k_{G1,r}C_{EG2PG1}-k_{G1,f}C_{PG1}C_{EG2} \tag{S1.14}$$

$$D_{G1}\frac{\partial C_{PG1}}{\partial r}(0,t)=0 \tag{S1.15}$$

$$C_{PG1}(r,0)=0 \tag{S1.16}$$

Conservation equation and initial and boundary conditions for $C_{G2}$:

$$\frac{\partial C_{G2}}{\partial t}=D_{G2}\frac{1}{r^2}\frac{\partial}{\partial r}\left(r^2\frac{\partial C_{G2}}{\partial r}\right)-k_{G1,f}C_{G1}C_{G2}+k_{G1,r}C_{G2G1}-k_{G1,f}C_{PG1}C_{G2}+$$
$$k_{G1,r}C_{G2PG1}-k_{G1,f}C_{G2}C_{PG1S}+k_{G1,r}C_{G2PG1S} \tag{S1.17}$$

$$D_{G2}\frac{\partial C_{G2}}{\partial r}\left(R,t\right)=k_{G2,r}C_{EG2}-k_{G2,f}C_{G2}C_{E} \tag{S1.18}$$

$$D_{G2}\frac{\partial C_{G2}}{\partial r}\left(0,t\right)=0 \tag{S1.19}$$

$$C_{G2}\left(r,0\right)=C_{o,GRB2} \tag{S1.20}$$

Conservation equation and initial and boundary conditions for $C_{G2G1}$:

$$\frac{\partial C_{G2G1}}{\partial t}=D_{G2G1}\frac{1}{r^2}\frac{\partial}{\partial r}\left(r^2\frac{\partial C_{G2G1}}{\partial r}\right)+k_{G1,f}C_{G1}C_{G2}-k_{G1,r}C_{G2G1}-$$
$$k_{G1p}C_{SFK,a}C_{G2G1}+k_{G1dp}C_{G2PG1} \tag{S1.21}$$

$$D_{G2G1}\frac{\partial C_{G2G1}}{\partial r}\left(R,t\right)=k_{G2,r}C_{EG2G1}-k_{G2,f}C_{G2G1}C_{E} \tag{S1.22}$$

$$D_{G2G1}\frac{\partial C_{G2G1}}{\partial r}\left(0,t\right)=0 \tag{S1.23}$$

$$C_{G2G1}\left(r,0\right)=0 \tag{S1.24}$$

Conservation equation and initial and boundary conditions for $C_{G2PG1}$:

$$\frac{\partial C_{G2PG1}}{\partial t}=D_{G2G1}\frac{1}{r^2}\frac{\partial}{\partial r}\left(r^2\frac{\partial C_{G2PG1}}{\partial r}\right)+k_{G1,f}C_{PG1}C_{G2}-k_{G1,r}C_{G2PG1}+k_{G1p}C_{SFK,a}C_{G2G1}-$$
$$k_{G1dp}C_{G2PG1}-k_{S2,f}C_{G2PG1}C_{S}+k_{S2,r}C_{G2PG1S} \tag{S1.25}$$

$$D_{G2G1}\frac{\partial C_{G2PG1}}{\partial r}\left(R,t\right)=k_{G2,r}C_{EG2PG1}-k_{G2,f}C_{G2PG1}C_{E} \tag{S1.26}$$

$$D_{G2G1}\frac{\partial C_{G2PG1}}{\partial r}\left(0,t\right)=0 \tag{S1.27}$$

$$C_{G2PG1}\left(r,0\right)=0 \tag{S1.28}$$

Conservation equation and initial and boundary conditions for $C_{S2}$:

$$\frac{\partial C_{S2}}{\partial t} = D_{S2}\frac{1}{r^2}\frac{\partial}{\partial r}\left(r^2\frac{\partial C_{S2}}{\partial r}\right) - k_{S2,f}C_{PG1}C_{S2} + k_{S2,r}C_{PG1S} - k_{S2,f}C_{G2PG1}C_{S2} + k_{S2,r}C_{G2PG1S} \tag{S1.29}$$

$$D_{S2}\frac{\partial C_{S2}}{\partial r}(R,t) = -k_{S2,f}C_{S2}C_{EG2PG1} + k_{S2,r}C_{EG2PG1S} \tag{S1.30}$$

$$D_{S2}\frac{\partial C_{S2}}{\partial r}(0,t) = 0 \tag{S1.31}$$

$$C_S(r,0) = C_{o,SHP2} \tag{S1.32}$$

Conservation equation and initial and boundary conditions for $C_{PG1S}$:

$$\frac{\partial C_{PG1S}}{\partial t} = D_{G1S2}\frac{1}{r^2}\frac{\partial}{\partial r}\left(r^2\frac{\partial C_{PG1S}}{\partial r}\right) + k_{S2,f}C_{PG1}C_{S2} - k_{S2,r}C_{PG1S} - k_{G1,f}C_{G2}C_{PG1S} + k_{G1,r}C_{G2PG1S} \tag{S1.33}$$

$$D_{G1S2}\frac{\partial C_{PG1S}}{\partial r}(R,t) = -k_{G1,f}C_{PG1S}C_{EG2} + k_{G1,r}C_{EG2PG1S} \tag{S1.34}$$

$$D_{G1S2}\frac{\partial C_{PG1S}}{\partial r}(0,t) = 0 \tag{S1.35}$$

$$C_{PG1S}(r,0) = 0 \tag{S1.36}$$

Conservation equation and initial and boundary conditions for $C_{G2PG1S}$:

$$\frac{\partial C_{G2PG1S}}{\partial t} = D_{G2G1S2}\frac{1}{r^2}\frac{\partial}{\partial r}\left(r^2\frac{\partial C_{G2PG1S}}{\partial r}\right) + k_{S2,f}C_{G2PG1}C_{S2} - k_{S2,r}C_{G2PG1S} + k_{G1,f}C_{G2}C_{PG1S} - k_{G1,r}C_{G2PG1S} \tag{S1.37}$$

$$D_{G2G1S2}\frac{\partial C_{G2PG1S}}{\partial r}(R,t) = -k_{G2,f}C_{G2PG1S}C_E + k_{G2,r}C_{EG2PG1S} \tag{S1.38}$$

$$D_{G2G1S2}\frac{\partial C_{G2PG1S}}{\partial r}(0,t) = 0 \tag{S1.39}$$

$$C_{G2PG1S}\left(r,0\right)=0 \tag{S1.40}$$

Conservation equation and initial condition for $C_{mE}$:

$$\frac{dC_{mE}}{dt}=-k_{E,f}C_{mE}\left[EGF\right]+k_{E,r}C_{mES} \tag{S1.41}$$

$$C_{mE}\left(0\right)=C_{o,EGFR} \tag{S1.42}$$

Conservation equation and initial condition for $C_{mES}$:

$$\frac{dC_{mES}}{dt}=k_{E,f}C_{o,EGF}C_{mE}-k_{E,r}C_{mES}-2k_{dE,f}C_{mES}{}^{2}+2k_{dE,r}C_{mESmES} \tag{S1.43}$$

$$C_{mES}\left(0\right)=0 \tag{S1.44}$$

Conservation equation and initial condition for $C_{mESmES}$:

$$\frac{dC_{mESmES}}{dt}=k_{dE,f}C_{mES}{}^{2}-k_{dE,r}C_{mESmES}-k_{catE}C_{mESmES}+k_{dp}C_{E} \tag{S1.45}$$

$$C_{mESmES}\left(0\right)=0 \tag{S1.46}$$

Conservation equation and initial condition for $C_{E}$:

$$\frac{dC_{E}}{dt}=-k_{G2,f}C_{G2}C_{E}+k_{G2,r}C_{EG2}-k_{G2,f}C_{G2G1}C_{E}+k_{G2,r}C_{EG2G1}-k_{G2,f}C_{G2PG1}C_{E}+ \\ k_{G2,r}C_{EG2PG1}-k_{G2,f}C_{G2PG1S}C_{E}+k_{G2,r}C_{EG2PG1S}+k_{catE}C_{mESmES}-k_{dp}C_{E} \tag{S1.47}$$

$$C_{E}\left(0\right)=0 \tag{S1.48}$$

Conservation equation and initial condition for $C_{EG2}$:

$$\frac{dC_{EG2}}{dt}=-k_{G1,f}C_{G1}C_{EG2}+k_{G1,r}C_{EG2G1}-k_{G1,f}C_{PG1}C_{EG2}+k_{G1,r}C_{EG2PG1}+ \\ k_{G2,f}C_{G2}C_{E}-k_{G2,r}C_{EG2}-k_{G1,f}C_{PG1S}C_{EG2}+k_{G1,r}C_{EG2PG1S} \tag{S1.49}$$

$$C_{EG2}\left(0\right)=0 \tag{S1.50}$$

Conservation equation and initial condition for $C_{EG2G1}$:

$$\frac{dC_{EG2G1}}{dt} = k_{G1,f} C_{G1} C_{EG2} - k_{G1,r} C_{EG2G1} + k_{G2,f} C_{G2G1} C_E - k_{G2,r} C_{EG2G1} \tag{S1.51}$$

$$C_{EG2G1}(0) = 0 \tag{S1.52}$$

Conservation equation and initial condition for $C_{EG2PG1}$:

$$\frac{dC_{EG2PG1}}{dt} = k_{G1,f} C_{PG1} C_{EG2} - k_{G1,r} C_{EG2PG1} + k_{G2,f} C_{G2PG1} C_E - k_{G2,r} C_{EG2PG1} - k_{S2,f} C_{S2} C_{EG2PG1} + k_{S2,r} C_{EG2PG1S} \tag{S1.53}$$

$$C_{EG2PG1}(0) = 0 \tag{S1.54}$$

Conservation equation and initial condition for $C_{EG2PG1S}$:

$$\frac{dC_{EG2PG1S}}{dt} = k_{S2,f} C_{S2} C_{EG2PG1} - k_{S2,r} C_{EG2PG1S} + k_{G1,f} C_{PG1S} C_{EG2} - k_{G2,r} C_{EG2PG1S} + k_{G2,f} C_{G2PG1S} C_E - k_{G2,r} C_{EG2PG1S} \tag{S1.55}$$

$$C_{EG2PG1S}(0) = 0 \tag{S1.56}$$

**Finite difference method: Ordinary differential equations**

Ordinary differential equations were discretized using a forward explicit finite difference approximation as follows:

$$\frac{dC}{dt} = k_1 C \tag{S1.57}$$

$$\frac{C_{i+1} - C_i}{h} = k_1 C_i \tag{S1.58}$$

where $h$ is the step size between two discretized time points ($h = t_{i+1} - t_i$). This discretized equation can be rearranged to solve for $C$ at time point $t_{i+1}$ ($C_{i+1}$), given that values for $C$ at time point $t_i$ ($C_i$) are known:

$$C_{i+1} = \left(h \cdot k_1 + 1\right) C_i \tag{S1.59}$$

**Finite difference method: Partial differential equations**

Partial differential equations were discretized using a forward explicit finite difference approximation for first-order time derivatives; an explicit first-order central finite difference approximation for first-order spatial derivatives; and a central explicit finite difference approximation for second-order spatial derivatives as follows:

$$\frac{\partial C}{\partial t} = D \frac{1}{r^2} \frac{\partial}{\partial r}\left( r^2 \frac{\partial C}{\partial r} \right) = D\left( \frac{2}{r} \frac{\partial C}{\partial r} + \frac{\partial^2 C}{\partial r^2} \right) \tag{S1.60}$$

$$\frac{C_{i+1,j} - C_{i,j}}{h} = D\left( \frac{2}{r_j} \frac{C_{i,j+1} - C_{i,j-1}}{2k} + \frac{C_{i,j+1} - 2C_{i,j} + C_{i,j-1}}{k^2} \right) \tag{S1.61}$$

where $h$ is the step size between two discretized time points ($h = t_{i+1} - t_i$) and $k$ is the step size between two discretized space points ($k = r_{j+1} - r_j$). This discretized equation can be rearranged to solve for $C$ at time point $t_{i+1}$ and space point $r_j$ ($C_{i+1,j}$), given that values for $C_i$ at space points $r_{j-1}$, $r_j$, and $r_{j+1}$ at time point $t_i$ are known ($C_{i,j-1}$, $C_{i,j}$, and $C_{i,j+1}$, respectively).

$$C_{i+1,j} = hD\left( \frac{1}{r_j} \frac{C_{i,j+1} - C_{i,j-1}}{k} + \frac{C_{i,j+1} - 2C_{i,j} + C_{i,j-1}}{k^2} \right) + hC_{i,j} \tag{S1.62}$$

To simultaneously solve the discretized equations for membrane-associated species, which are coupled with the discretized boundary conditions for cytosolic species, a semi-implicit finite difference method was utilized, shown here for an example involving the discretized equations for arbitrary membrane-associated species $A$ and $B$ and the discretized boundary condition at the cell membrane for one cytosolic species ($C$):

Discretization for $A$:

$$\frac{dA}{dt} = -C \cdot A + B \tag{S1.63}$$

$$\frac{A_{i+1} - A_i}{h} = -C_{i+1,N} \cdot A_i + B_i \tag{S1.64}$$

Discretization for *B*:

$$\frac{dB}{dt} = C \cdot A - B \tag{S1.65}$$

$$\frac{B_{i+1} - B_i}{h} = C_{i+1,N} \cdot A_i - B_i \tag{S1.66}$$

Discretization for *C*'s membrane boundary condition:

$$D_C \frac{\partial C}{\partial r}\left(R,t\right) = C \cdot A - B \tag{S1.67}$$

$$D \frac{C_{i+1,N} - C_{i+1,N-1}}{k} = C_{i+1,N} \cdot A_{i+1} - B_{i+1} \tag{S1.68}$$

Due to the fact that $A_{i+1}$, $B_{i+1}$, and $C_{i+1,N}$ are all unknown, initial guesses are provided for $A_{i+1}$ and $B_{i+1}$ to solve for $C_{i+1,N}$, which is then used to solve for $A_{i+1}$ and $B_{i+1}$ according to the discretized equations above. These updated values are then used to re-solve for $C_{i+1,N}$, and this process is iterated until the fractional error between initial guesses and updated values for $A_{i+1}$, $B_{i+1}$, and $C_{i+1,N}$ (illustrated in Eq. S1.69 below for $A_{i+1}$) is less than the desired tolerance:

$$\left| 1 - \frac{A_{i+1,0}}{A_{i+1,f}} \right| < \text{tol} \tag{S1.69}$$

where $A_{i+1,0}$ is the initial guess and $A_{i+1,f}$ is the updated value from evaluating the model equations.

Discretization of boundary conditions for cytosolic species at the cell center results in the requirement for the concentration of a given cytosolic species to be equivalent at the cell center node and the space node immediately adjacent to it, as illustrated below:

$$D\frac{dC}{dr}(0)=0 \tag{S1.70}$$

$$D\frac{C_{i+1}-C_i}{k}=0 \tag{S1.71}$$

$$C_{i+1}=C_i \tag{S1.72}$$

Once all partial and ordinary differential equations for cytosolic and membrane-associated species, respectively, were discretized using the finite difference schemes above, they were then solved as follows:

1) Concentrations of all species were defined for the initial time.
2) The model was advanced one time step, and the concentrations of all cytosolic species were solved for in the bulk (i.e., everywhere except the cell center and cell surface boundaries).
3) At the same time step, the concentrations of all cytosolic species at the cell center were solved for using the appropriate boundary conditions at the cell center.
4) At the same time step, the concentrations of all cytosolic species at the cell surface were solved for using the appropriate boundary conditions at the cell surface, where initial guesses for concentrations of all membrane-associated species were provided.
5) The concentrations of all membrane-associated species were solved for using the concentrations of cytosolic species at the cell surface calculated in Step 5.
6) Steps 5 and 6 were repeated, where the initial guesses for the concentrations of membrane-associated species in Step 5 were updated using the concentrations calculated in the previous iteration of Step 6. This process was iterated until the concentrations obtained in Steps 5 and 6 converged within the desired error tolerance.
7) Steps 3-7 were repeated until the final timepoint was reached.

#### Appendix S2: Steady-state model equations and simplifications.

##### Derivation of reduced model equations

Additional model calculations were performed with a simplified set of model equations in which EGFR and GRB2 were neglected. Neglecting these two species was in part justified by the model sensitivity analysis (Fig. 3A), which indicated that the abundance and persistence of GAB1-SHP2 complexes throughout the cell was relatively insensitive to changes in the binding and unbinding rate constants of GAB1 to GRB2 ($k_{G1,f}$, $k_{G1,r}$) and GRB2 to EGFR ($k_{G2,f}$, $k_{G2,r}$). The original set of model PDEs was thus reduced to a set of six PDEs describing active and inactive SFKs, GAB1, pGAB1, SHP2, and pGAB1-SHP2. All model equations were then converted to their steady-state forms by setting the time derivatives in each equation to zero. The concentration of phosphorylated EGFR, which is present in the membrane boundary conditions for active and inactive SFKs, was set to the predicted value of total phospho-EGFR from the solution to the full PDE model after 5 minutes of a 10 ng/mL EGF treatment.

Applying the above simplifications results in the following conservation equations and boundary conditions for the remaining cytosolic species:

$C_{SFK,i}$ conservation equation and boundary conditions:

$$D_{SFK}\frac{1}{r^2}\frac{d}{dr}\left(r^2\frac{dC_{SFK,i}}{dr}\right)+k_{S,i}C_{SFK,a}=0 \tag{S2.1}$$

$$D_{SFK}\frac{dC_{SFK,i}}{dr}\left(R\right)=-k_{S,a}C_{pE,tot}C_{SFK,i} \tag{S2.2}$$

$$\frac{dC_{SFK,i}}{dr}\left(0\right)=0 \tag{S2.3}$$

Conservation equation and boundary conditions for $C_{SFK,a}$ at steady state:

$$D_{SFK}\frac{1}{r^2}\frac{d}{dr}\left(r^2\frac{dC_{SFK,a}}{dr}\right)-k_{S,i}C_{SFK,a}=0 \tag{S2.4}$$

$$D_{SFK}\frac{dC_{SFK,a}}{dr}(R)=k_{S,a}C_{pE,tot}C_{SFK,i} \tag{S2.5}$$

$$\frac{dC_{SFK,a}}{dr}(0)=0 \tag{S2.6}$$

Conservation equation and initial and boundary conditions for $C_{G1}$ at steady state:

$$0=D_{G1}\frac{1}{r^2}\frac{d}{dr}\left(r^2\frac{dC_{G1}}{dr}\right)-k_{G1p}C_{SFK,a}C_{G1}+k_{G1dp}C_{PG1} \tag{S2.7}$$

$$\frac{dC_{G1}}{dr}(R)=0 \tag{S2.8}$$

$$\frac{dC_{G1}}{dr}(0)=0 \tag{S2.9}$$

Conservation equation and initial and boundary conditions for $C_{PG1}$ at steady state:

$$0=D_{G1}\frac{1}{r^2}\frac{d}{dr}\left(r^2\frac{dC_{PG1}}{dr}\right)+k_{G1p}C_{SFK,a}C_{G1}-k_{G1dp}C_{PG1}-k_{S2,f}C_{PG1}C_{S2}+k_{S2,r}C_{PG1S} \tag{S2.10}$$

$$\frac{dC_{PG1}}{dr}(R)=0 \tag{S2.11}$$

$$\frac{dC_{PG1}}{dr}(0)=0 \tag{S2.12}$$

Conservation equation and initial and boundary conditions for $C_{PG1S}$ at steady state:

$$0=D_{G1S2}\frac{1}{r^2}\frac{d}{dr}\left(r^2\frac{dC_{PG1S}}{dr}\right)+k_{S2,f}C_{PG1}C_{S2}-k_{S2,r}C_{PG1S} \tag{S2.13}$$

$$\frac{dC_{PG1S}}{dr}(R)=0 \tag{S2.14}$$

$$\frac{dC_{PG1S}}{dr}(0)=0 \qquad \text{(S2.15)}$$

Conservation equation and initial and boundary conditions for $C_{S2}$ at steady state:

$$0=D_{S2}\frac{1}{r^2}\frac{d}{dr}\left(r^2\frac{dC_{S2}}{dr}\right)-k_{S2,f}C_{PG1}C_{S2}+k_{S2,r}C_{PG1S} \qquad \text{(S2.16)}$$

$$\frac{dC_{S2}}{dr}(R)=0 \qquad \text{(S2.17)}$$

$$\frac{dC_{S2}}{dr}(0)=0 \qquad \text{(S2.18)}$$

Steady-state solutions for active and inactive SFKs were obtained analytically, as described in the next section. Steady-state solutions to the four PDEs above were obtained numerically using a modified finite difference approach (also described in the next section) and the analytical solutions for active and inactive SFKs. Additional solutions were obtained by further reducing the set of four ODEs to a system of two ODEs and to a single ODE using a conserved scalar approach. These simplifications are outlined in the next sections.

#### Analytical solutions for active and inactive SFKs at steady state

The conservation equations for active and inactive SFKs can be readily solved to obtain analytical solutions for the concentration profiles of each species at steady state. The conservation equations for these species are coupled, but this can be resolved by adding the equations together and using the mass balance for total SFKs to obtain a single ordinary differential equation (ODE).

The steady-state conservation equation for active SFKs can be rearranged as

$$\frac{1}{r^2}\frac{d}{dr}\left(r^2\frac{dC_{SFK,a}}{dr}\right)-\frac{k_{S,i}}{D_{SFK}}C_{SFK,a}=0 \qquad \text{(S2.19)}$$

Eq. S2.19 is a form of the modified spherical Bessel's equation with $n = 0$ and $m = \left( \frac{k_{S,i}}{D_{SFK}} \right)^{1/2}$.

The general solution to this equation is

$$C_{SFK,a}\left(r\right) = \alpha \frac{\sinh mr}{mr} + \beta \frac{\cosh mr}{mr} \tag{S2.20}$$

Differentiating Eq. S2.20 and applying the no-flux boundary condition for active SFKs at $r = 0$ yields $\beta = 0$ and

$$C_{SFK,a}\left(r\right) = \alpha \frac{\sinh mr}{mr} \tag{S2.21}$$

To solve for $\alpha$, the reactive boundary condition at $r = R$ must be rewritten in terms of $C_{SFK,a}$ by substituting for $C_{SFK,i}$. The relation between $C_{SFK,a}$ and $C_{SFK,i}$ can be obtained by first adding the conservation equations for active and inactive SFKs to obtain

$$\frac{D_{SFK}}{r^2} \frac{d}{dr}\left( r^2 \frac{dC_{SFK,a}}{dr} \right) + \frac{D_{SFK}}{r^2} \frac{d}{dr}\left( r^2 \frac{dC_{SFK,i}}{dr} \right) = 0 \tag{S2.22}$$

Dividing by $\frac{D_{SFK}}{r^2}$ and integrating twice gives

$$C_{SFK,a} + C_{SFK,i} = c_1 r + c_2 \tag{S2.23}$$

where $c_1$ and $c_2$ are unknown constants. Differentiating Eq. S2.23 and applying the no-flux boundary conditions at $r = 0$ gives $c_1 = 0$ and

$$C_{SFK,a} + C_{SFK,i} = c_2 \tag{S2.24}$$

The mass balance for total SFKs indicates that $c_2 = C_{o,SFK}$. Substituting into Eq. S2.24 and rearranging gives

$$C_{SFK,i} = C_{o,SFK} - C_{SFK,a} \tag{S2.25}$$

The membrane boundary condition for active SFKs can then be written as

$$\frac{dC_{SFK,a}}{dr}(R)=\frac{k_{S,a}C_{pE,tot}}{D_{SFK}}\left(C_{o,SFK}-C_{SFK,a}\big|_{r=R}\right) \qquad \text{(S2.26)}$$

where $C_{pE,tot}$ is a known constant at steady state (taken from the 5-min time point of the full PDE solution). Evaluating the derivative of Eq. S2.21 at $r = R$, equating it to Eq. S2.26, and solving for α gives

$$\alpha=\frac{k_{S,a}C_{pE,tot}C_{o,SFK}}{D_{SFK}}\left[\frac{\cosh mR}{R}+\frac{\sinh mR}{mR}\left(\frac{k_{S,a}C_{pE,tot}}{D_S}-\frac{1}{R}\right)\right]^{-1} \qquad \text{(S2.27)}$$

Finally, substituting for α in Eq. S2.21 gives the steady-state solution for active SFKs,

$$C_{SFK,a}(r)=\frac{k_{S,a}C_{pE,tot}C_{o,SFK}}{D_{SFK}}\left[\frac{\cosh mR}{R}+\frac{\sinh mR}{mR}\left(\frac{k_{S,a}C_{pE,tot}}{D_S}-\frac{1}{R}\right)\right]^{-1}\frac{\sinh mr}{mr} \qquad \text{(S2.28)}$$

The solution at $r = 0$ can be obtained by applying L'Hôpital's rule to Eq. S2.28 in the limit $r \rightarrow 0$, which yields

$$C_{SFK,a}(0)=\alpha \qquad \text{(S2.29)}$$

**Model simplification using conserved scalars**

The four simplified conservation equations describing GAB1, pGAB1, pGAB1-SHP2, and SHP2 can be further reduced to a set of two ODEs using conserved scalar quantities. At steady state, the equations for GAB1, pGAB1, and pGAB1-SHP2 can be added together to give

$$D_{G1}\left[\frac{d}{dr}\left(r^2\frac{dC_{G1}}{dr}\right)+\frac{d}{dr}\left(r^2\frac{dC_{PG1}}{dr}\right)\right]+D_{G1S2}\frac{d}{dr}\left(r^2\frac{dC_{PG1S}}{dr}\right)=0 \qquad \text{(S2.30)}$$

Integrating twice and applying the no-flux boundary conditions gives

$$D_{G1}\left(C_{G1}+C_{PG1}\right)+D_{G1S2}C_{PG1S}=c_3 \qquad \text{(S2.31)}$$

Neglecting differences in diffusivity, $c_3$ is equal to the total concentration of GAB1, $C_{o,GAB1}$:

$$C_{G1} + C_{PG1} + C_{PG1S} = C_{o,GAB1} \tag{S2.32}$$

Adding the conservation equations for pGAB1-SHP2 and SHP2 and integrating likewise gives

$$D_{S2}C_{S2} + D_{G1S2}C_{PG1S} = c_4 \tag{S2.33}$$

Again neglecting differences in diffusivities, $c_4$ is equal to the total concentration of SHP2, $C_{o,SHP2}$:

$$C_{S2} + C_{PG1S} = C_{o,SHP2} \tag{S2.34}$$

Eqs. S2.32 and S2.34 can be used to reduce number of equations that must be solved numerically to two. Substituting for the concentrations of GAB1 and SHP2, the conservation equations for pGAB1 and GAB1-SHP2 become

$$\begin{aligned}\frac{\partial C_{PG1}}{\partial t} = D_{G1}\frac{1}{r^2}\frac{\partial}{\partial r}\left(r^2\frac{\partial C_{PG1}}{\partial r}\right) + k_{G1p}C_{SFK,a}\left(C_{o,G1} - C_{PG1} - C_{PG1S}\right) - k_{G1dp}C_{PG1} - \\ k_{S2,f}C_{PG1}\left(C_{o,G1} - C_{PG1S}\right) + k_{S2,r}C_{PG1S}\end{aligned} \tag{S2.35}$$

$$\frac{\partial C_{PG1S}}{\partial t} = D_{G1S2}\frac{1}{r^2}\frac{\partial}{\partial r}\left(r^2\frac{\partial C_{PG1S}}{\partial r}\right) + k_{S2,f}C_{PG1}\left(C_{o,G1} - C_{PG1S}\right) - k_{S2,r}C_{PG1S} \tag{S2.36}$$

Eqs. S2.35 and S2.36 are then solved using the modified finite difference approach outlined at the end of this supplement, with the concentration profiles for GAB1 and SHP2 obtained using Eqs. S2.32 and S2.34.

**<u>Further model simplification using pGAB1-SHP2 binding equilibrium</u>**

If pGAB1 and SHP2 are at or near binding equilibrium, then Eqs. S2.35 and S2.36 can be reduced to a single differential equation. At equilibrium,

$$\frac{k_{S2,on}}{k_{S2,off}} = \frac{C_{PG1S}}{C_{PG1}C_{S2}} \tag{S2.37}$$

Using Eqs. S2.31, S2.33, and S2.36 to substitute for $C_{S2}$ and $C_{PG1S}$, the conservation equation for pGAB1 can be written as

$$D_{G1}\frac{1}{r^2}\frac{d}{dr}\left(r^2\frac{dC_{PG1}}{dr}\right)+k_{G1p}C_{SFK,a}\left[C_{o,G1}-C_{PG1}-C_{o,G1}\left(\frac{k_{S2,off}}{k_{S2,on}C_{PG1}}+1\right)^{-1}\right]-k_{G1dp}C_{PG1}=0 \quad \text{(S2.38)}$$

Eq. S2.38 is then solved using the finite differences method described below.

**Modified steady-state finite difference scheme**

The steady-state conservation equations and boundary conditions for GAB1, pGAB1, pGAB1-SHP2, and SHP2 were discretized in space using the same procedure as outlined in Appendix S1. Concentration profiles for active and inactive SFKs were obtained using the analytical solutions, which are outlined in the next section. Mass balance constraints for GAB1 and SHP2 were placed on the system accordingly:

$$0=\frac{3}{R^3}\int_0^R\left(C_{G1}+C_{PG1}+C_{PG1S}\right)r^2dr-C_{o,GAB1} \quad \text{(S2.39)}$$

$$0=\frac{3}{R^3}\int_0^R\left(C_{S2}+C_{PG1S}\right)r^2dr-C_{o,S2} \quad \text{(S2.40)}$$

where the integrals in Eqs. S2.39 and S2.40 were approximated with the trapezoidal rule. The final result is a system of nonlinear algebraic equations of the form $f(x) = 0$ which was solved using the *fsolve* function in MATLAB.

#### Appendix S3: Scaling and order-of-magnitude derivations of SFK and GAB1-SHP2 length scale relationships.

##### Estimation of active SFK and GAB1-SHP2 length scales

Estimates for the length scales over which active SFKs and GAB1-SHP2 persist from the cell membrane were estimated using standard scaling analysis methods, the full details of which are described by Deen[1]. The analysis was carried out assuming the system described in Appendix S1 has reached steady state, which corresponds with the final time point in the results shown in Fig. 2A and Fig. S1A. Because the diffusivities of GAB1-SHP2 and GRB2-GAB1-SHP2 are nearly identical (Table 1), these two species were considered identical for the purposes of this analysis. Thus, the length scale for GAB1-SHP2 is also taken as the length scale for GRB2-GAB1-SHP2.

The length scale of SFK activation can be derived from the steady-state conservation equation,

$$D_{SFK}\frac{1}{r^2}\frac{d}{dr}\left(r^2\frac{dC_{SFK,a}}{dr}\right)-k_{S,i}C_{SFK,a}=0 \qquad \text{(S3.1)}$$

Let the dimensionless concentration for active SFKs, $\theta$, be defined as

$$\theta=\frac{C_{SFK,a}}{C_{o,SFK}} \qquad \text{(S3.2)}$$

where $C_{o,SFK}$ is the initial concentration of inactive SFKs. The differential of the dimensionless concentration is given by

$$d\theta=\frac{1}{C_{o,SFK}}dC_{SFK,a} \qquad \text{(S3.3)}$$

Similarly, let the dimensionless radial coordinate, $\eta$, and its differential be defined as

$$\eta = \frac{r}{R} \quad \text{(S3.4)}$$

$$d\eta = \frac{1}{R}dr \quad \text{(S3.5)}$$

where $R$ is the cell radius. Substituting Eqs. S3.2-S3.5 into Eq. S3.1 and rearranging give

$$\frac{1}{\eta^2}\frac{d}{d\eta}\left(\eta^2\frac{d\theta}{d\eta}\right) = \frac{k_{S,i}R^2}{D_{SFK}}\theta \quad \text{(S3.6)}$$

The leading ratio on the right-hand side of Eq. S3.6 is known as the Damköhler number, Da, which describes the balance between the inactivation and diffusion of active SFKs within the cell. Substituting in Da, Eq. S3.6 is rewritten as

$$\frac{1}{\eta^2}\frac{d}{d\eta}\left(\eta^2\frac{d\theta}{d\eta}\right) = \mathrm{Da}\theta \quad \text{(S3.7)}$$

In cases where Da >> 1, Eq. S3.7 will be improperly scaled. To avoid this and maintain all terms ~1, $\eta$ must be rescaled using

$$\chi = \eta \mathrm{Da}^{1/2} = r\left(\frac{k_{S,i}}{D_{SFK}}\right)^{1/2} \quad \text{(S3.8)}$$

where $\chi$ is now the properly scaled and dimensionless radial coordinate. The true length scale for the radial coordinate and active SFKs is therefore given by,

$$\delta_{SFK,a} \sim \left(\frac{D_{SFK}}{k_{S,i}}\right)^{1/2} \quad \text{(S3.9)}$$

This is the distance over which large changes in the concentration of active SFKs are expected to occur within a cell.

Using the same scaling analysis approach to derive a length scale GAB1-SHP2 complexes is complicated by the presence of multiple kinetic terms in the conservation equations for pGAB1-SHP2 and GRB2-pGAB1-SHP2. An alternative approach is to estimate and sum the length scales associated with the processes that cooperate to maintain GAB1-SHP2 complexes. These processes are: (i) SFK-mediated GAB1 phosphorylation, (ii) GAB1-SHP2 dissociation, and (iii) GAB1 dephosphorylation. Estimating the length scale in this way assumes that complexes form wherever GAB1 is being actively phosphorylated and wherever it remains phosphorylated and that intact complexes require a finite amount of time to dissociate. Using the same approach and reasoning to derive $\delta_{SFK,a}$, the length scale associated with GAB1-SHP2 complex dissociation, $\delta_{dis}$, can be written as

$$\delta_{dis} \sim \left( \frac{D_{G1S2}}{k_{S2,off}} \right)^{1/2} \tag{S3.10}$$

The length scale for GAB1 dephosphorylation, $\delta_{dep}$, describes the balance between GAB1 dephosphorylation and diffusion and is written as

$$\delta_{dep} \sim \left( \frac{D_{G1}}{k_{G1dp}} \right)^{1/2} \tag{S3.11}$$

Since SHP2 may still bind to pGAB1 over $\delta_{dep}$ to reform the GAB1-SHP2 complex, it may be reasoned that GAB1-SHP2 likewise persists over this length scale. Thus, the length scale over which large changes in the concentration of GAB1-SHP2 are expected to occur can be estimated by summing Eqs. S3.9-S3.11 to obtain

$$\delta_{G1S2} \sim \left( \frac{D_{SFK}}{k_{S,i}} \right)^{1/2} + \left( \frac{D_{G1S2}}{k_{S2,off}} \right)^{1/2} + \left( \frac{D_{G1}}{k_{G1dp}} \right)^{1/2} \tag{S3.12}$$

#### Appendix S4: Estimation of Protein and Protein Complex Diffusivities.

Diffusivities for each cytosolic protein monomer or complex were calculated by adjusting the diffusivity of tubulin[2] based on differences in molecular weight between the model species of interest and tubulin; correlating those differences with reported hydrodynamic radii (Stokes radii) of several protein standards[3]; and then linearly interpolating hydrodynamic radii for tubulin and the model species based on those standards. First, the hydrodynamic radius of tubulin was interpolated using standards reported by Erickson[3]. The mass of tubulin was taken to be about 50 kDa. Using this molecular weight, the hydrodynamic radius of tubulin was linearly interpolated as 3.18 nm using the masses and hydrodynamic radii of ovalbumin hen egg and albumin beef serum. The same procedure was repeated for each of the model species using the protein standards reported by Erickson[3].

Protein and complex diffusivities were then estimated by adjusting the diffusivity of tubulin based on differences in the hydrodynamic radii of tubulin and the model species. From measurements made by Pepperkok *et al.*[2], the diffusivity of tubulin was taken to be approximately 87 μm²/min. Assuming that the relationship between the hydrodynamic radius and diffusivity of a protein is linear, the diffusivity of model proteins and complexes was then estimated using

$$D_i = \frac{D_{tub} R_{S,tub}}{R_{S,i}} \tag{S4.1}$$

where $D_i$ is the diffusivity of protein or complex $i$, $D_{tub}$ is the diffusivity of tubulin, $R_{S,i}$ is the Stokes radius of the protein/complex $i$, and $R_{S,tub}$ is the Stokes radius of tubulin. The results of these calculations for each model protein or complex are summarized in Table S2.

#### Supplementary Figures and Tables

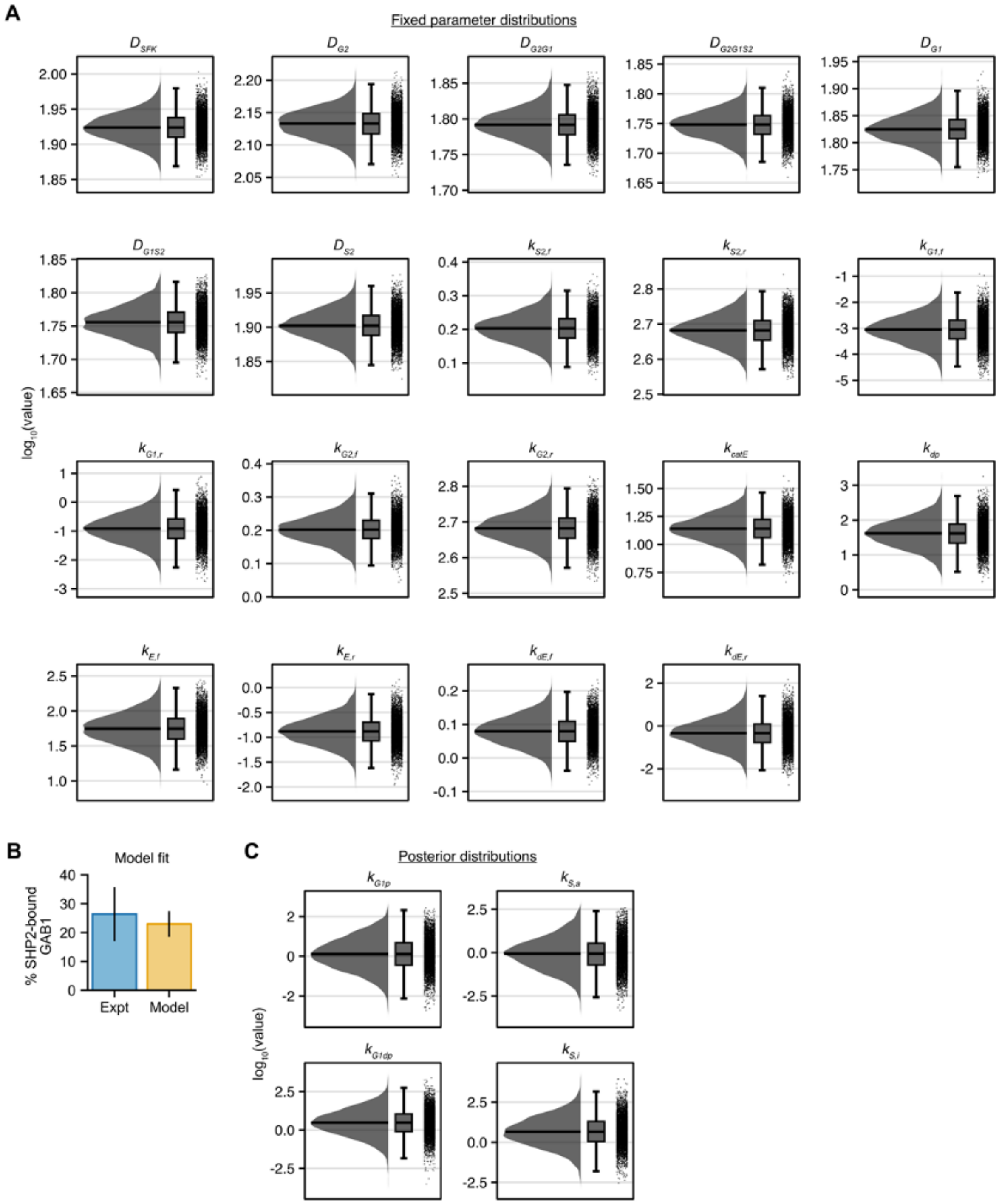


**Fig. S1. The fitted reaction-diffusion model matches experimental measurements of GAB1-SHP2 association.** (**A**) Raincloud plots are shown for all fixed (non-fitted) model parameters based on 5,000 samples from each parameter's distribution. The median of each distribution is indicated as a solid black line. (**B**) Experimental measurements for the fraction of SHP2-bound GAB1 from H1666 cells treated with 10 ng/mL EGF[4] were compared against fitted model ensemble predictions for 5 min of 10 ng/mL EGF, as described in Section 2-3. Experimental error bar indicates mean ± s.e.m. The error bar on model predictions indicates the 68.4% credible interval (roughly equivalent to one standard deviation) around the median model prediction calculated from the full set of 5,000 parameter posterior samples. (**C**) Posterior distributions are shown for the fitted model parameters based on all 5,000 posterior samples. The median of each distribution is indicated as a solid black line.

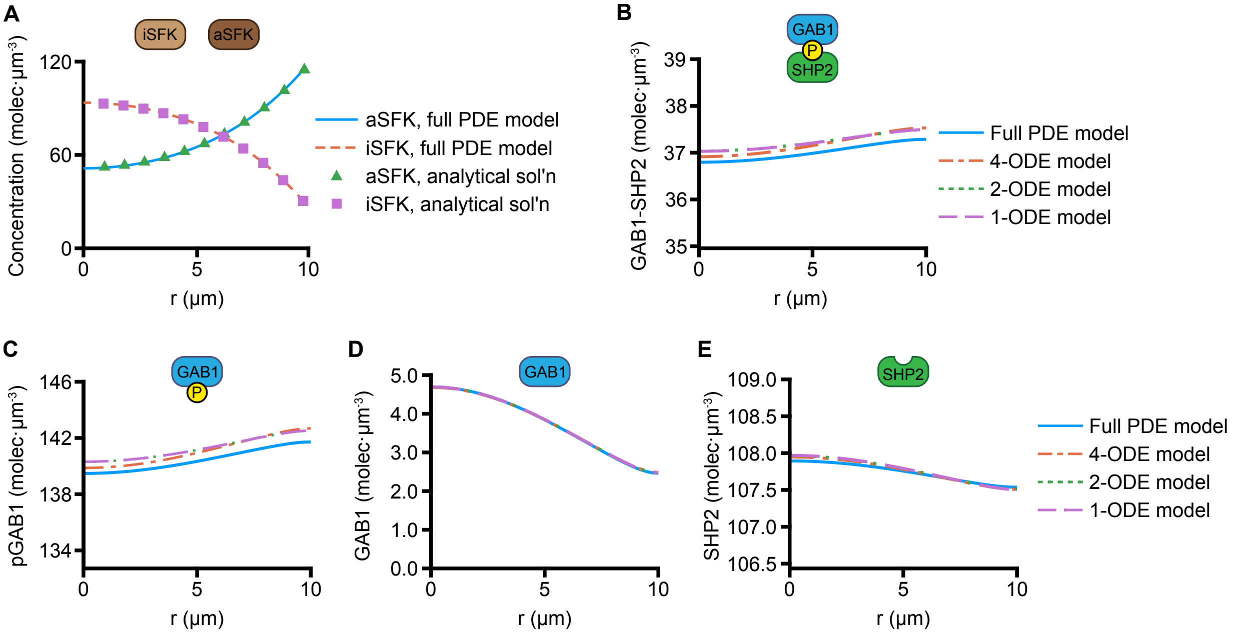


**Fig. S2. Steady-state model predictions predict that GAB1-SHP2 complexes are evenly distributed throughout cells in response to EGF treatment.** (**A**) Concentration profiles for active and inactive SFKs after a 5-min 10 ng/mL EGF treatment from the full reaction-diffusion model as a function of distance from the cell center *r* were compared to the analytical, steady-state solutions obtained by fixing the concentration of phosphorylated EGFR to the full model value at *t* = 5 min and solving the steady-state conservation equations for active and inactive SFKs. (**B-E**) Concentration profiles were calculated with the full PDE model for total GAB1-SHP2, total pGAB1, GAB1, and SHP2 as a function of *r* and compared to predictions from the steady-state reduced model without EGFR and GRB2 binding (4-ODE model), the reduced model simplified with conserved scalars (2-ODE model), and the reduced model simplified with conserved scalars and GAB1-SHP2 binding equilibrium (1 ODE model). Predictions from the full PDE model were taken from the 5-min time point of a 10 ng/mL EGF treatment. Predictions from the 4-ODE, 2-ODE, and 1-ODE models were obtained using a modified finite difference scheme without time steps. See *Methods* and Appendix S2 for details. Base model parameter values were used for all calculations as defined in *Methods*.

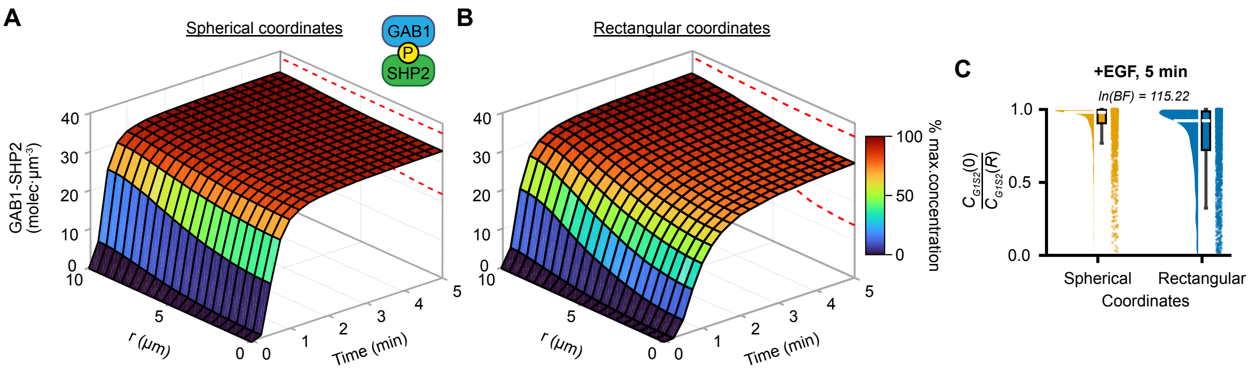


**Fig. S3. Comparison of model predictions in spherical versus rectangular coordinates.** (**A-B**) Model ensemble predictions are shown for the concentration of GABl-SHP2 complexes in response to 10 ng/mL EGF as a function of distance from the cell center ($r$) and time for model equations in spherical or rectangular coordinates using 2,000 draws from model parameter distributions. Surfaces represent the median of ensemble predictions, and dashed red lines enclose the 68.2% credible interval centered around the median at $t = 5$ min. (**C**) Model predictions for the ratio of GAB1-SHP2 concentrations at the cell center to the cell surface ($C_{G1S2}(0)/C_{G1S2}(R)$) were computed for the model ensembles in **A** and **B** at $t = 5$ min. The natural logarithm of the Bayes factor (BF) indicates strong evidence in favor of the alternative hypothesis that the difference in the distribution means is different than zero (i.e., strong evidence that there is a difference in the predicted ratio between spherical and rectangular coordinates).

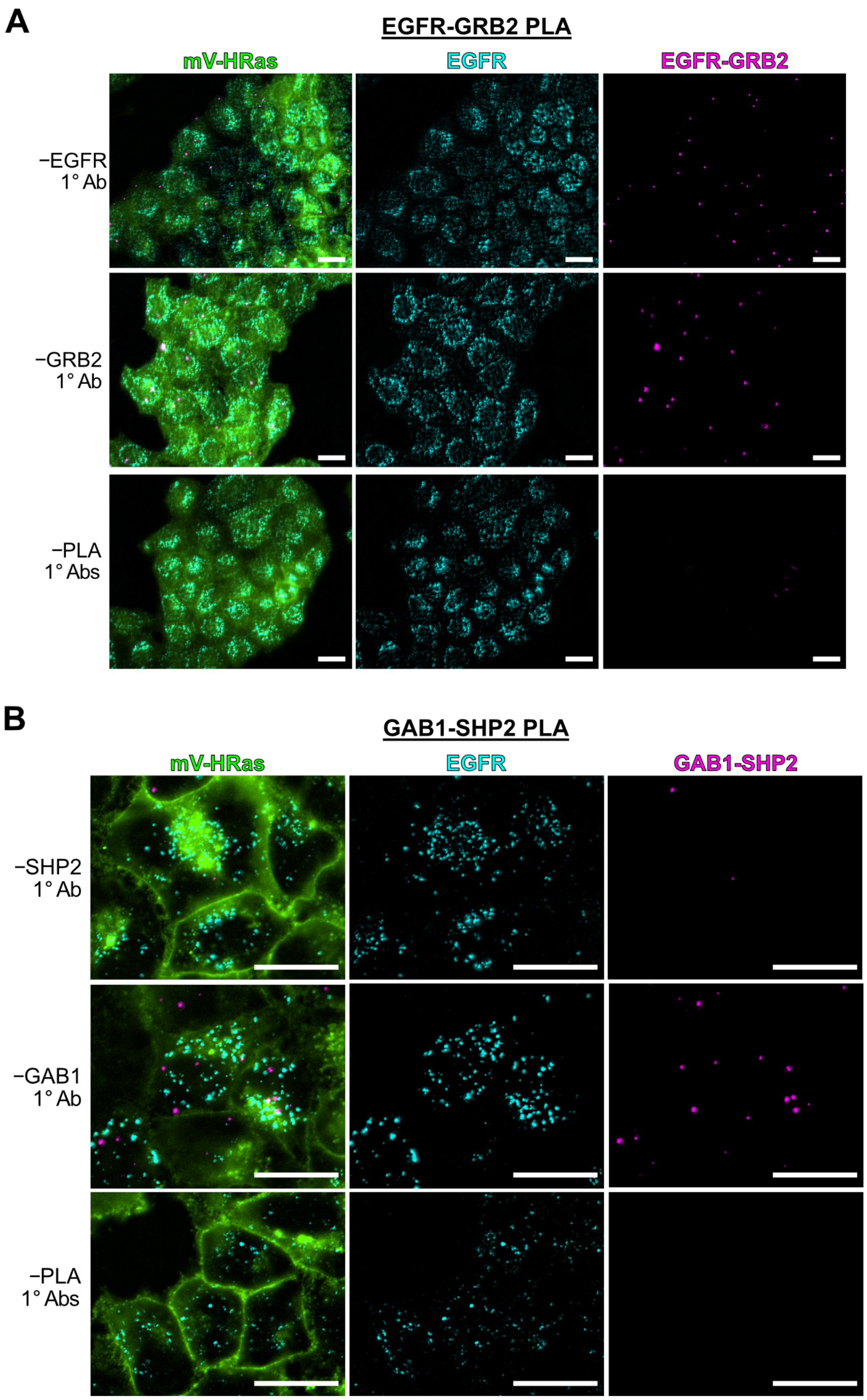


**Fig. S4. Proximity ligation assay primary antibody controls.** (**A**) Images of EGFR-GRB2 PLA primary antibody control samples. HeLa/mVenus-HRAS that were treated with 10 ng/mL EGF for 7 min, fixed, and stained for EGFR and EGFR-GRB2 by IF and PLA, respectively, without individual PLA primary antibodies as indicated. Scale bars, 20 μm. Images are representative of n = 3 independent coverslips per condition. (**B**) Images of GAB1-SHP2 PLA primary antibody control samples. HeLa/mVenus-HRAS that were treated with 10 ng/mL EGF for 15 min, fixed, and stained for EGFR and GAB1-SHP2 by IF and PLA, respectively, without individual PLA primary antibodies as indicated. Maximum-intensity *z*-projections from epifluorescence imaging are shown. Scale bars, 20 μm. Images are representative of n = 5 independent coverslips per condition.

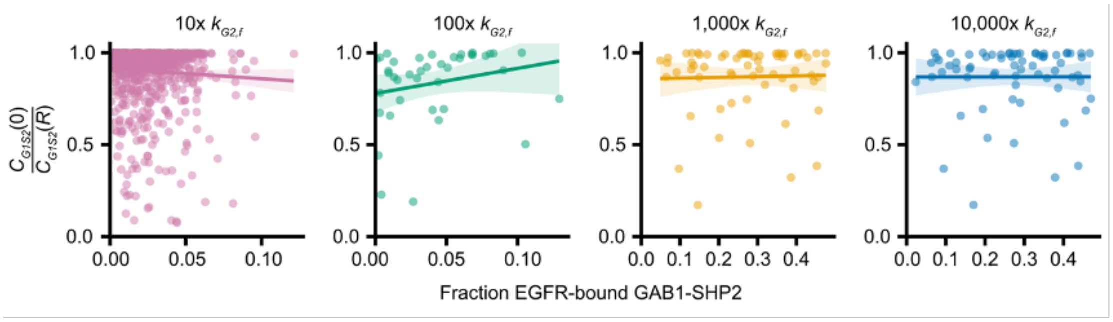


**Fig. S5. Increasing GAB1-SHP2 binding to EGFR negligibly affects the length scale of complex persistence.** Model predictions for the center-to-surface ratio of GAB1-SHP2 concentrations ($C_{G1S2}(0)/C_{G1S2}(R)$) were performed for 1,000 samples from model parameter ensembles and when multiplying the value for the rate constant of EGFR-GRB2 forward binding ($k_{G2,f}$) in each parameter set by the indicated values above each plot panel. Note that many model simulations became unstable when making these changes, which is why not all panels contain the same number of points. Linear trendlines indicating the relationship between the center-to-surface ratio and the fraction of EGFR-bound GAB1-SHP2 are indicated on each plot with shading indicating trendline 95% confidence intervals.

**Table S1. Parameter values used to generate fixed and prior parameter distributions.**

| Parameter (units) | Description | Reported values | Uncertainty | Source |
|---|---|---|---|---|
| $k_{E,f}$ (μM$^{-1}$ min$^{-1}$) | EGF binding to EGFR, forward | 63 | ± 19 (s.d.) | [5] |
| $k_{E,r}$ (min$^{-1}$) | EGF binding to EGFR, reverse | 0.16 | ± 0.05 (s.d.) | [5] |
| $k_{dE,f}$ (μm$^{2}$ molec$^{-1}$ min$^{-1}$) | EGFR dimerization, forward | 1.2 | N.A.—assumed ×/÷ 1.1 | [4] |
| $K_{D,dim}$ (molec μm$^{-2}$) | EGFR dimer dissociation constant | 0.38 | ×/÷ 4.67 (estimated multiplicative s.d.) | [6] |
| $k_{catE}$ (min$^{-1}$) | EGFR phosphorylation, EGF-occupied dimer | 14.4 (Y992)<br>17.4 (Y1068)<br>7.2 (Y1086)<br>12.9 (Y1114)<br>13.1 (Y1148)<br>15.1 (Y1173) | ± 0.5 (s.e.m, n = 4)<br>± 0.6 (s.e.m, n = 4)<br>± 0.3 (s.e.m, n = 4)<br>± 0.4 (s.e.m, n = 4)<br>± 0.4 (s.e.m, n = 4)<br>± 0.2 (s.e.m, n = 4) | [7] |
| $k_{dp}$ (min$^{-1}$) | EGFR dephosphorylation | 8.0<br>40.2 (Y1068)<br>52.8 (Y992)<br>36 (Y1148)<br>127.2 (Y1173) | N.A.—assumed ×/÷ 1.1<br>± 2.76 (s.e.m, assumed n = 2)<br>± 9.6 (s.e.m, assumed n = 2)<br>± 14 (s.e.m, assumed n = 2)<br>± 37.8 (s.e.m, assumed n = 2) | [4]<br>[8]<br>[8]<br>[8]<br>[8] |
| $k_{S,a}$ (μm$^{3}$ molec$^{-1}$ min$^{-1}$) | SFK activation | 0.42 | N.A.—assumed ×/÷ 1.1 | [4] |
| $k_{S,i}$ (min$^{-1}$) | SFK inactivation | 9.5 | N.A.—assumed ×/÷ 1.1 | [4] |
| $k_{G2,f}$ (μm$^{3}$ molec$^{-1}$ min$^{-1}$) | GRB2 binding to EGFR, forward | 1.594 | ×/÷ 1.1 (estimated) | [9] |
| $k_{G2,r}$ (min$^{-1}$) | GRB2 binding to EGFR, reverse | 480 | ×/÷ 1.1 (estimated) | [9] |
| $K_{D,G2}$ (molec μm$^{-3}$) | GRB2 binding to EGFR, dissociation | 60.22 | ×/÷ 3 (estimated) | [9] |
| $k_{G1,f}$ (μm$^{3}$ molec$^{-1}$ min$^{-1}$) | GAB1 binding to GRB2, forward | $2.4 \times 10^{3}$<br>$6.4 \times 10^{4}$<br>$9.5 \times 10^{4}$<br>$1.1 \times 10^{3}$<br>$7.8 \times 10^{3}$<br>$1.5 \times 10^{4}$<br>$1.3 \times 10^{3}$<br>$2.4 \times 10^{4}$<br>$0.9 \times 10^{3}$ | $\pm 0.1 \times 10^{3}$<br>$\pm 0.1 \times 10^{4}$<br>$\pm 0.1 \times 10^{4}$<br>$\pm 7.0 \times 10^{3}$<br>$\pm 0.1 \times 10^{3}$<br>$\pm 0.2 \times 10^{4}$<br>$\pm 0.2 \times 10^{3}$<br>$\pm 0.3 \times 10^{4}$<br>$\pm 0.1 \times 10^{3}$ | [10] |
| $k_{G1,r}$ (min$^{-1}$) | GAB1 binding to GRB2, reverse | $3.9 \times 10^{-2}$<br>$1.9 \times 10^{-3}$<br>$2.2 \times 10^{-3}$<br>$3.0 \times 10^{-3}$<br>$9.9 \times 10^{-4}$<br>$2.2 \times 10^{-3}$<br>$1.6 \times 10^{-3}$<br>$3.2 \times 10^{-3}$<br>$1.6 \times 10^{-3}$ | $\pm 0.2 \times 10^{-2}$<br>$\pm 0.2 \times 10^{-3}$<br>$\pm 0.1 \times 10^{-3}$<br>$\pm 0.1 \times 10^{-3}$<br>$\pm 0.2 \times 10^{-4}$<br>$\pm 0.3 \times 10^{-3}$<br>$\pm 0.3 \times 10^{-3}$<br>$\pm 0.3 \times 10^{-3}$<br>$\pm 0.04 \times 10^{-3}$ | [10] |

| | | | | |
|---|---|---|---|---|
| $k_{G1p}$ ($\mu m^3$ $molec^{-1}$ $min^{-1}$) | GAB1 phosphorylation | 0.42 | N.A.—assumed ×/÷ 1.1 | [4] |
| $k_{G1dp}$ ($min^{-1}$) | GAB1 dephosphorylation | 9.5 | N.A.—assumed ×/÷ 1.1 | [4] |
| $k_{S2,f}$ ($\mu m^3$ $molec^{-1}$ $min^{-1}$) | SHP2 binding to phosphorylated GAB1, forward | 1.594 | ×/÷ 1.1 (estimated) | [9] |
| $k_{S2,r}$ ($min^{-1}$) | SHP2 binding to phosphorylated GAB1, reverse | 480 | ×/÷ 1.1 (estimated) | [9] |
| $K_{D,S2}$ (molec $\mu m^{-3}$) | SHP2 binding to phosphorylated GAB1, dissociation | 60.22 | ×/÷ 3 (estimated) | [9] |
| $D_{SFK}$ ($\mu m^2$ $min^{-1}$) | Diffusivity of SFK molecules | $8.2\times10^1$ | Assumed ×/÷ 1.1 | [2,3] |
| $D_{G2}$ ($\mu m^2$ $min^{-1}$) | Diffusivity of GRB2 molecules | $1.3\times10^2$ | Assumed ×/÷ 1.1 | [2,3] |
| $D_{G2G1}$ ($\mu m^2$ $min^{-1}$) | Diffusivity of GRB2-GAB1 molecules | $6.1\times10^1$ | Assumed ×/÷ 1.1 | [2,3] |
| $D_{G2G1S2}$ ($\mu m^2$ $min^{-1}$) | Diffusivity of GRB2-GAB1-SHP2 [illegible] | $5.5\times10^1$ | Assumed ×/÷ 1.1 | [2,3] |
| $D_{G1}$ ($\mu m^2$ $min^{-1}$) | Diffusivity of GAB1 molecules | $6.6\times10^1$ | Assumed ×/÷ 1.1 | [2,3] |
| $D_{G1S2}$ ($\mu m^2$ $min^{-1}$) | Diffusivity of GAB1-SHP2 molecules | $5.6\times10^1$ | Assumed ×/÷ 1.1 | [2,3] |
| $D_{S2}$ ($\mu m^2$ $min^{-1}$) | Diffusivity of SHP2 molecules | $7.8\times10^1$ | Assumed ×/÷ 1.1 | [2,3] |

**Table S2: Masses, hydrodynamic radii, and diffusivities of model proteins.** $R_S$ is the hydrodynamic (or Stokes) radius in nm, and $D$ is diffusivity in ($\mu m^2$ $min^{-1}$).

| Protein or complex | Molecular weight (Da) | $R_S$ (nm) | $D$ ($\mu m^2$ $min^{-1}$) |
|---|---|---|---|
| Tubulin | 50,000 | 3.18 | 87 |
| SFK | 59,835 | 3.37 | 82 |
| GRB2 | 25,206 | 2.07 | 130 |
| GAB1 | 115000 | 4.20 | 66 |
| SHP2 | 68,436 | 3.53 | 78 |
| GRB2-GAB1 | 140,206 | 4.56 | 61 |
| GAB1-SHP2 | 183,436 | 4.93 | 56 |
| GRB2-GAB1-SHP2 | 208,642 | 5.05 | 55 |

#### SUPPLEMENTARY REFERENCES

1. Deen, W.M. (1998). Analysis of Transport Phenomena, 2 Edition (Oxford University Press New York).
2. Pepperkok, R., Bre, M.H., Davoust, J., and Kreis, T.E. (1990). Microtubules are stabilized in confluent epithelial cells but not in fibroblasts. J Cell Biol *111*, 3003-3012. 10.1083/jcb.111.6.3003.
3. Erickson, H.P. (2009). Size and Shape of Protein Molecules at the Nanometer Level Determined by Sedimentation, Gel Filtration, and Electron Microscopy. Biol Proced Online *11*, 32-51. 10.1007/s12575-009-9008-x.
4. Furcht, C.M., Buonato, J.M., and Lazzara, M.J. (2015). EGFR-activated Src family kinases maintain GAB1-SHP2 complexes distal from EGFR. Science Signaling *8*. 10.1126/scisignal.2005697.
5. French, A.R., Tadaki, D.K., Niyogi, S.K., and Lauffenburger, D.A. (1995). Intracellular Trafficking of Epidermal Growth-Factor Family Ligands Is Directly Influenced by the Ph Sensitivity of the Receptor-Ligand Interaction. J Biol Chem *270*, 4334-4340. DOI 10.1074/jbc.270.9.4334.
6. Hajdu, T., Váradi, T., Rebenku, I., Kovács, T., Szöllösi, J., and Nagy, P. (2020). Comprehensive Model for Epidermal Growth Factor Receptor Ligand Binding Involving Conformational States of the Extracellular and the Kinase Domains. Front Cell Dev Biol *8*, 776. 10.3389/fcell.2020.00776.
7. Fan, Y.X., Wong, L., Deb, T.B., and Johnson, G.R. (2004). Ligand regulates epidermal growth factor receptor kinase specificity - Activation increases preference for GAB1 and SHC versus autophosphorylation sites. J Biol Chem *279*, 38143-38150. 10.1074/jbc.M405760200.
8. Young, M.W. (2019). Dephosphorylation of Epidermal Growth Factor Receptor by Protein Tyrosine Phosphatase 1B. The FASEB journal *33*.
9. Morimatsu, M., Takagi, H., Ota, K.G., Iwamoto, R., Yanagida, T., and Sako, Y. (2007). Multiple-state reactions between the epidermal growth factor receotor and Grb2 as observed by using single-molecule analysis. P Natl Acad Sci USA *104*, 18013-18018. 10.1073/pnas.0701330104.
10. Kiyatkin, A., Aksamitiene, E., Markevich, N.I., Borisov, N.M., Hoek, J.B., and Kholodenko, B.N. (2006). Scaffolding protein Grb2-associated binder 1 sustains epidermal growth factor-induced mitogenic and survival signaling by multiple positive feedback loops. J Biol Chem *281*, 19925-19938. 10.1074/jbc.M600482200.
